# Microbes use transporters to regulate the release of metabolites based on value

**DOI:** 10.1101/2025.08.19.671024

**Authors:** Snorre Sulheim, Gunn Broli, Peter F. Doubleday, Alisson Gillon, Julien S. Luneau, Prajwal Padmanabha, Andrew Quinn, Eric Ulrich, Margaret A. Vogel, Nicola Zamboni, Philipp Engel, Sara Mitri

## Abstract

Microbial interactions are shaped by the exchange of metabolites, ranging from intermediates in central carbon metabolism to amino acids, vitamins and fermentation by-products. While specific metabolic processes such as overflow metabolism are well documented, a general principle explaining the chemical diversity of metabolite release and the variation in individual release rates remains elusive. Here, using a combination of computational and experimental approaches on seven different microbes, we show that release rates are negatively correlated with the value of metabolites, and that this relationship explains more variability in release rates than other frequently associated factors. By measuring metabolite release from 67 *E. coli* mutants lacking individual transport genes, we find that transporters both cause and counteract net metabolite release. These findings suggest that metabolite release constitutes a fitness cost that microbes have evolved to reduce, while revealing the underlying transport mechanism. This unifying explanation reshapes our understanding of how microbial interactions emerge and persist.

## Introduction

Metabolites are frequently exchanged between microbes in natural communities^1^. Such cross-feeding plays a key role in maintaining community diversity and function^2–6^. While certain occurrences of cross-feeding arise from extracellular metabolism such as polymer degradation by extracellular enzymes^6,7^, many observations of cross-feeding stem from the release of intracellular metabolites into the environment by microbial community members^8–10^. Recent *in vitro* quantifications of microbial cultures have demonstrated that this release includes high-energy glycolytic intermediates, amino acids, vitamins and organic acids^11–15^. The ubiquity of metabolite release may help explain the high frequency of auxotrophs isolated from natural environments that may be supported by co-cultivated partners^4,15–19^. However, why so many diverse intracellular metabolites are being released and what explains differences in release rates remain unanswered.

Previous efforts to answer why microbes release intracellular metabolites have revealed fundamental principles of metabolism, including rate-yield trade-offs and constraints on proteome allocation^20–24^, mechanisms of pathway regulation and maintenance of homeostasis^25^, optimal catabolic pathway lengths in different environments^26,27^ and redox balancing^28,29^. These explanations are context-, metabolite- or species-specific. Cell lysis and passive diffusion across the cell membrane are mechanisms that apply more broadly across diverse taxa and environments, but it remains unclear whether these mechanisms alone can explain observed rates^12,30–34^. Thus, a unifying principle for metabolite release that applies across the spectrum of chemical diversity, microbial species and environments, and is able to explain variations in release rates is lacking.

Here, we approach the problem by asking: does metabolite release generally confer a benefit or impose a cost to the producer? Low or negligible biosynthetic costs have previously been linked to metabolite release^35–37^, and the trade-off between growth and production is a well-documented consequence of mass-balance^38,39^. However, whether biosynthetic costs systematically shape exometabolome diversity and release rates has not been tested. While recent work has explored how metabolite release can be beneficial, e.g. by alleviating costs associated with molecular noise^33,40^, we here propose and test an alternative hypothesis that metabolite release comes with a cost that microbes have evolved to reduce.

## Results

### Metabolite release rates are negatively correlated with metabolite value

If metabolite release generally imposes a fitness cost, and natural selection acts to minimize such costs, then net release rates of specific metabolites should be negatively correlated with their relative importance, or value, to the microbe (Fig. 1A). Release of more valuable metabolites – that require more energy to produce – should have a more severe negative impact on fitness and be more strongly selected against.

**Figure 1:**
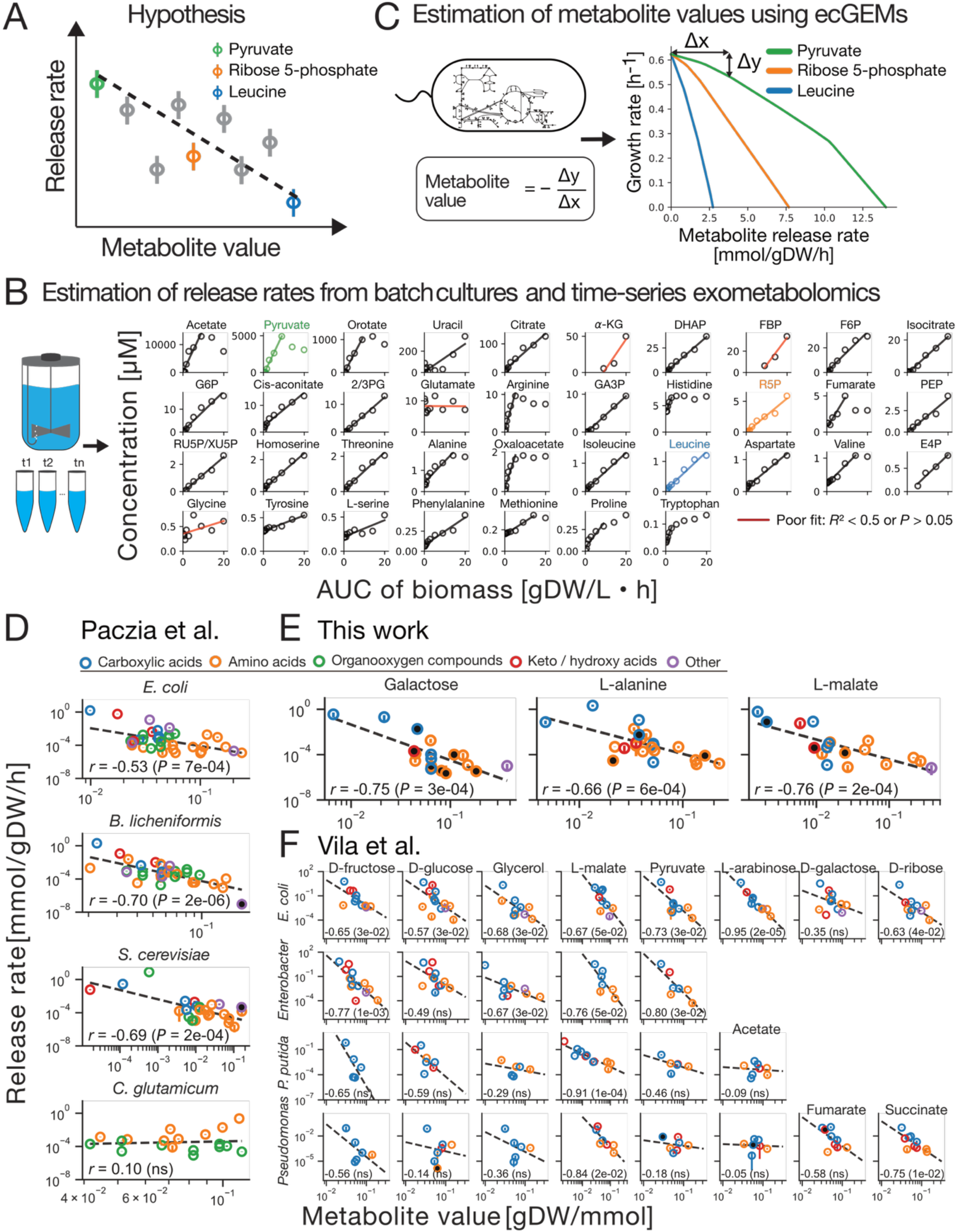
Correlation between microbial metabolite release rates and metabolite values. A) Hypothesis: if metabolite release generally constitutes a fitness cost that microbes have evolved to reduce, metabolite release rates should be negatively correlated with metabolite values. Three metabolites are used to conceptually illustrate their relative position based on their estimated release rate and metabolite value, see panels B and C. B) Metabolite release rates were estimated from time-series exometabolomics data from batch cultures by linear regression of extracellular concentrations vs area under the curve (AUC) of biomass. This panel shows extracellular concentrations of 37 metabolites measured during exponential phase of *E. coli* cultivated in glucose medium by Paczia et al. (2012), and the linear regressions used to estimate net release rates. Linear regressions with an *R*^2^ < 0.5 or *P* > 0.05 (two-sided Wald test) are drawn in red. Full time-series, including the stationary phase and linear regression fits for *E. coli*, *B. licheniformis*, *C. glutamicum* and *S. cerevisiae* are shown in Figs. S1-S4. Metabolite abbreviations: α-KG: alpha-ketoglutarate, DHAP: dihydroxyacetone phosphate, FBP: fructose 1,6-bisphosphate, F6P: fructose 6-phosphate, G6P: glucose 6-phosphate, 2/3PG: 2/3-phosphoglycerate, GA3P: glyceraldehyde 3-phosphate, R5P: ribose 5-phosphate, PEP: phosphoenolpyruvate, RU5P/XU5P: ribulose 5-phosphate/xylulose 5-phosphate, E4P: erythrose 4-phosphate. C) We used enzyme-constrained genome-scale metabolic models to estimate metabolite values, quantified as the negative change in growth rate (Δy) on metabolite release (Δx) close to optimal growth. D-F) Three different datasets were used to test our hypothesis: D) Paczia et al. (2012), including *S. cerevisiae*, *B. licheniformis*, *C. glutamicum* and *E. coli* on glucose^12^, E) *E. coli* on three other carbon sources (this work), and F) Vila et al. (2023) covering a smaller number of metabolites on *E. coli, P. putida* and two environmental strains of the genera *Enterobacter* and *Pseudomonas* on up to 8 different carbon sources^49^. The scatter plots show mean estimated rates ± standard errors vs estimated metabolite values. Black-filled circles are datapoints where the lower error bar is omitted because mean minus standard error is less than 0 and cannot be included on a log-scale. Axis labels and color legend are shared for panels D-F. Annotations in panels D-F show the Pearson correlation and associated *P* value in parentheses.

To quantify metabolite release rates across different species, we first analyzed an existing time-series exometabolome dataset with high temporal resolution and absolute quantification of 19 to 37 metabolites collected from batch cultures of *E. coli*, *B. licheniformis, S. cerevisiae* and *C. glutamicum* in high glucose concentrations^12^. We estimated net uptake and release rates for all metabolites using simple linear regression, modeling extracellular metabolite concentration as a linear function of the area under the growth curve (AUC), until the point where the extracellular concentration saturated or the exponential growth phase ended (Methods, Fig. 1B, and Figs. S1-S4). The initial dynamics of 44-95% of metabolites were well described by this simple linear model (two-sided Wald test, *P* < 0.05, *R*^2^ > 0.5, Figs. S1-S4).

To estimate metabolite value, we used previously reconstructed enzyme-constrained genome-scale metabolic models (ecGEMs)^41–43^. Genome-scale metabolic models (GEMs) are mathematical representations of a species’ metabolic capacity that can for example be used to predict optimal metabolic phenotypes across various environments^44,45^. The additional enzyme constraints account for the total enzyme pool limitation in cells^21^, enabling prediction of relevant trade-offs and temporal dynamics, e.g. acetate overflow in *E. coli*^42^. Various measures of metabolite value have been proposed in the past^37,46,47^. Here we define metabolite value as the negative change in growth rate on metabolite release close to optimal growth as predicted by enzyme-constrained GEMs (Fig. 1C). This measure explicitly reflects the fitness penalty on growth rate associated with the marginal release of a metabolite, and it can easily be used for different species across different environmental contexts.

With these two measures, we could now test our hypothesis that release rate should be negatively correlated with metabolite value. Indeed, this holds for *E. coli, S. cerevisiae and B. licheniformis* in glucose medium (Pearson *r*, *P* < 0.001, Fig. 1D). This did not hold for *C. glutamicum* (Fig. 1D), possibly because the engineering of its central carbon metabolism for enhanced lysine production has also affected the release of other metabolites^48^. The diversity of metabolites measured for *C. glutamicum* was also more limited than for the other species. However, when we restricted the analysis for the other species to the same compound classes (amino acids and organooxygen compounds), the correlation remained negative for all three species, although weaker and no longer significant for *E. coli* (Pearson *r* = −0.63 to −0.23).

To test whether our hypothesis held beyond simple glucose environments, we conducted new well-controlled batch cultivations of *E. coli* in minimal media with galactose, L-malate or L-alanine as the single carbon source. These carbon sources were chosen to span the variability of *E. coli* metabolism and avoid biasing the results towards a particular class of compounds or metabolic flux patterns (Methods, Fig. S5), given that similar carbon sources have been shown to produce similar exometabolome patterns^49^. Triplicate batch bioreactor experiments were performed under strictly controlled aerobic conditions with active regulation of pH and dissolved oxygen (Fig. S6). Relative extracellular concentrations of 98 metabolites mapping to the *E. coli* metabolome were then quantified in three exponential-phase samples and one sample from stationary phase. Of these metabolites, 45-57% were released at significant rates during exponential phase (two-sided Wald test, FDR < 0.05, Figs. S7-S10), consistent with previous findings showing that microbes generate rich exometabolomes^12,13^. Absolute quantification was only achieved for a subset of the metabolites, and ultimately, we were able to compare absolute release rates with estimated metabolite values for 27 different metabolites. Again, we found significant negative correlations for all three conditions (Pearson *r*, *P* < 0.001, Fig. 1E).

Finally, we analyzed a third dataset that in addition to *E. coli* and *Pseudomonas putida* contained environmental isolates from the genera *Enterobacter* and *Pseudomonas*^49^. Unlike the previous datasets, these experiments were conducted without pH and oxygen control, offering an opportunity to test the hypothesis under more variable conditions. Despite these differences, we again found consistent negative correlations (Fig. 1F), suggesting that the relationship between metabolite value and release rate is robust across species, nutrient conditions, and experimental platforms. Importantly, the ecGEMs used here were reconstructed by different research groups using different pipelines, suggesting that our results are not dependent on the choices made during model reconstruction. This conclusion was also supported by consistent results when metabolite values were estimated using four other *E. coli* GEMs (Fig. S11).

### Metabolite value is the most important factor explaining variability in metabolite release rates

The consistent negative correlation between metabolite release rates and metabolite value encouraged us to ask how well this factor explains release variability compared to other metabolic or physicochemical properties previously associated with metabolite release^40,50^. We focused on the data from bioreactor batch cultures of *E. coli* with glucose, galactose, L-malate or L-alanine as the carbon source, where conditions were well-defined, the measured metabolites were diverse and reference values for intracellular metabolite concentrations were available^51–54^. We first built univariate linear models to quantify the importance of these factors: metabolite value, intracellular concentration, compound class, octanol-water partitioning (log *P*), molecular weight, topological polar surface area, charge, hydrogen bond acceptor and donor count, rotatable bond count, turnover and carbon source (Fig. 2A). Metabolite value explained 43% of the variability, more than any other factor (Fig. 2B). Given that release rates are constrained by multiple factors such as intracellular concentration, passive diffusion rates and transporter expression levels, it is remarkable that such a substantial effect can be explained by metabolite value alone. In contrast, only 8% of the variability was explained by intracellular concentration. The structure-based classification of compound class also had good predictive power (29% explained), primarily because amino acids tended to be released at lower rates than carboxylic and keto/hydroxy acids (Fig. 2A). However, these two factors are clearly confounded as class-level differences mirrored significant differences in metabolite values (Fig. 2C). The number of rotatable bonds, log *P* and the number of hydrogen bond donors, each related to membrane permeability^55,56^, were also predictive of metabolite release, suggesting passive diffusion as a relevant mechanism. One might expect the effect of intracellular concentration to increase with membrane permeability, but we found at most a non-significant trend in this direction (interaction tests; all adjusted *P* values ≥ 0.21; see Methods). Similarly, we found little support for the noise-averaging cooperation hypothesis^40^, which predicts that release rates should decrease with metabolite turnover because low-turnover metabolites are more susceptible to noise and therefore more useful to share within the population.

**Figure 2:**
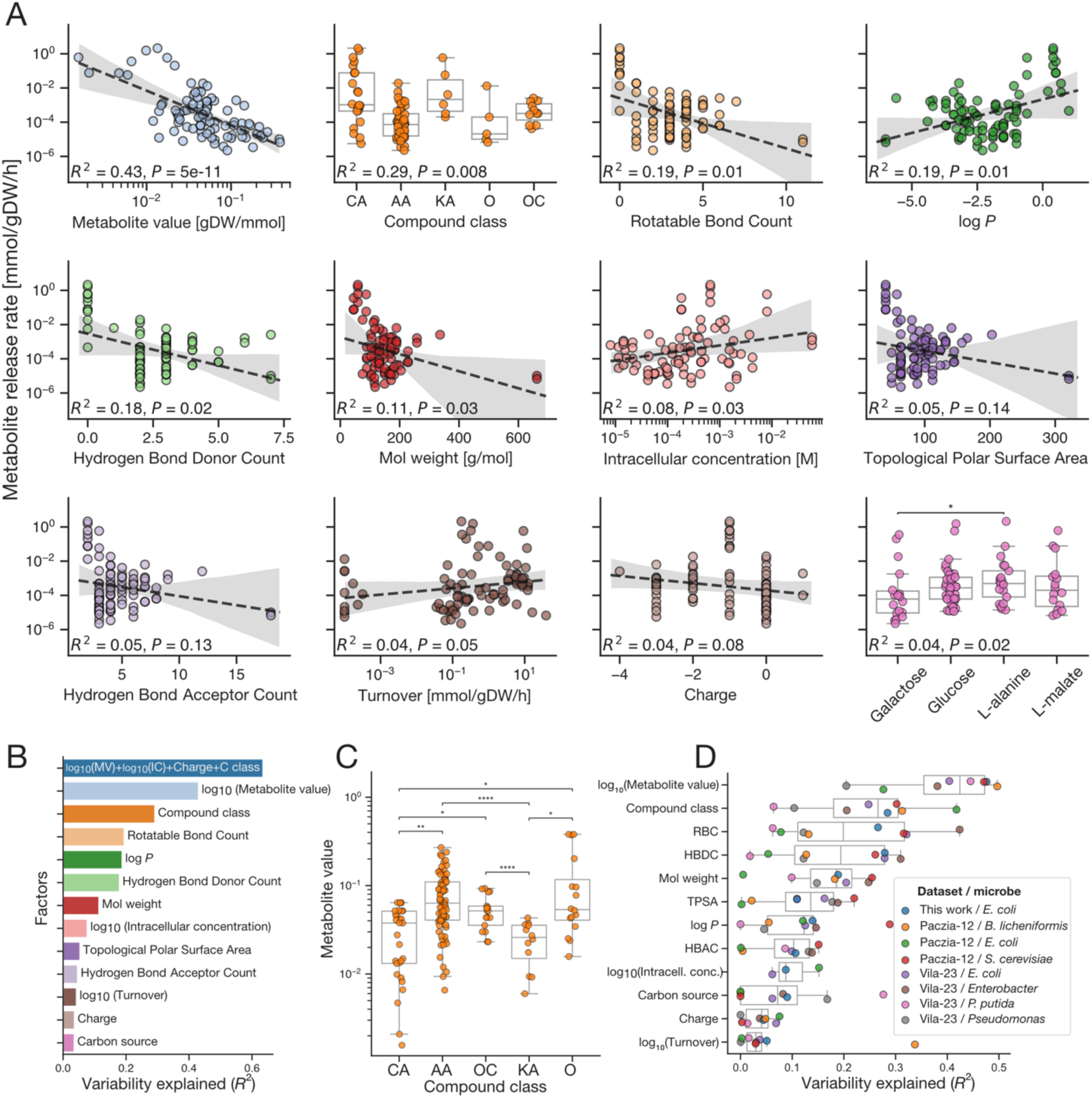
The importance of different factors in explaining metabolite release rates. A) Among the tested chemical and metabolic factors, log-scaled metabolite value explains the most variability in log-scaled *E. coli* release rates (n = 92 observations of 48 metabolites). The tested factors are shown in order of how much variability (*R*^2^) in log-scaled metabolite release rates they explain as a univariate linear model, shown in panel B. Because several metabolites were measured on multiple carbon sources, all P values were calculated from linear models with standard errors clustered by metabolite. *For continuous factors, P values test the regression slope, and shaded areas show 95% confidence intervals bootstrapped over metabolites. For compound class and carbon source, P values are from a cluster-robust Wald test of the overall group effect* and *pairwise comparisons are cluster-robust Wald tests with Benjamini–Hochberg correction (*: P ≤ 0.05; **: P ≤ 0.01; ***: P ≤ 0.001; ****: P ≤ 0.0001; non-significant comparisons not shown)*. Compound class abbreviations: CA, Carboxylic acids; AA, Amino acids; KA, Keto/hydroxy acids; OC, Organooxygen compounds; O, Other. B) The best linear model, ranked by BIC score, is a four-factor model that explains 63% of the variability in log-scaled release rates. C) Differences in metabolite values between compound classes (n = 153 observations of 53 metabolites). Abbreviations and statistical test as in A. D) Log-scaled metabolite value consistently ranks among the most important factors when evaluated separately for each species in each dataset. Factor abbreviations: RBC, Rotatable Bond Count; HBDC, Hydrogen Bond Donor Count; TPSA, Topological Polar Surface Area; HBAC, Hydrogen Bond Acceptor Count.

When we asked how well the best linear model could predict metabolite release rates, we found that 63% of the variability could be explained by metabolite value, compound class, intracellular concentration and charge (Fig. 2B). However, predictive accuracy declined when applied to out-of-sample conditions (45%, *P* = 0.04, 52.9 ± 3.8% expected from random model, Fig. S12) or out-of-sample metabolites (50%, *P* < 0.001, 55.2 ± 0.7% expected from random model, Fig. S13), likely due to correlations among the factors, which warrants caution when interpreting factors’ relative importance (Fig. S14). Nevertheless, excluding metabolite value as a factor reduced model performance on out-of-sample conditions (10% decline in median *R*^2^, *P* = 0.003, one-sided Mann-Whitney U, N = 20, Fig. S15), and model quality was largely dependent on including metabolite value or compound class (Fig. S16). Furthermore, we found differences in metabolite value between conditions to be negatively correlated with differences in release rate (Pearson *r, P =* 0.003, Fig. S17). This not only supports the relevance of this factor in predicting out-of-sample rates but also raises the question of whether microbes adapt their release rates to context-dependent costs. Finally, when we estimated each factor’s importance separately for each microbe in each dataset (except *C. glutamicum*), metabolite value still emerged as the most robust explanatory factor (Fig. 2D).

Cell lysis is commonly considered a key mechanism for metabolite release^32,33^. To measure its importance in our experiments, we used *E. coli* batch cultures with paired sampling of intra- and extracellular metabolites, as well as flow cytometry with live/dead staining to quantify the fraction of lysed cells. In general, less than 10% of extracellular concentrations could be explained by the release of intracellular metabolites following cell lysis, and often much less (Fig. 3A). We then extended this analysis by comparing our collection of *E. coli* extracellular metabolite concentrations to published intracellular concentrations^51–54^. To avoid asserting equal lysis rates across conditions, we asked how large a fraction of the cell population would need to be lysed to account for the extracellular levels, considering only the contribution from intracellular metabolite pools (Fig. 3B). Only 10.4% (157/1508) of the data points fall within the range of measured lysis fraction (0.2-2.3%, mean ± std across conditions = 0.6 ± 0.6%) or below, and 56.3% (849/1508) are above the limit where more than the whole cell population is required. Glutamate, glutamine and NAD are the only metabolites for which most data points can be accounted for by cell lysis, due to high intracellular concentrations and moderate to low extracellular levels (Fig. S18A-B). Furthermore, the fraction of lysed cells needed to explain extracellular concentrations decreased with metabolite value (Spearman *ρ* = −0.35, *P* = 0.016; Fig. S18C), suggesting that lysis may be a relatively more important route for the release of high-value metabolites.

**Figure 3:**
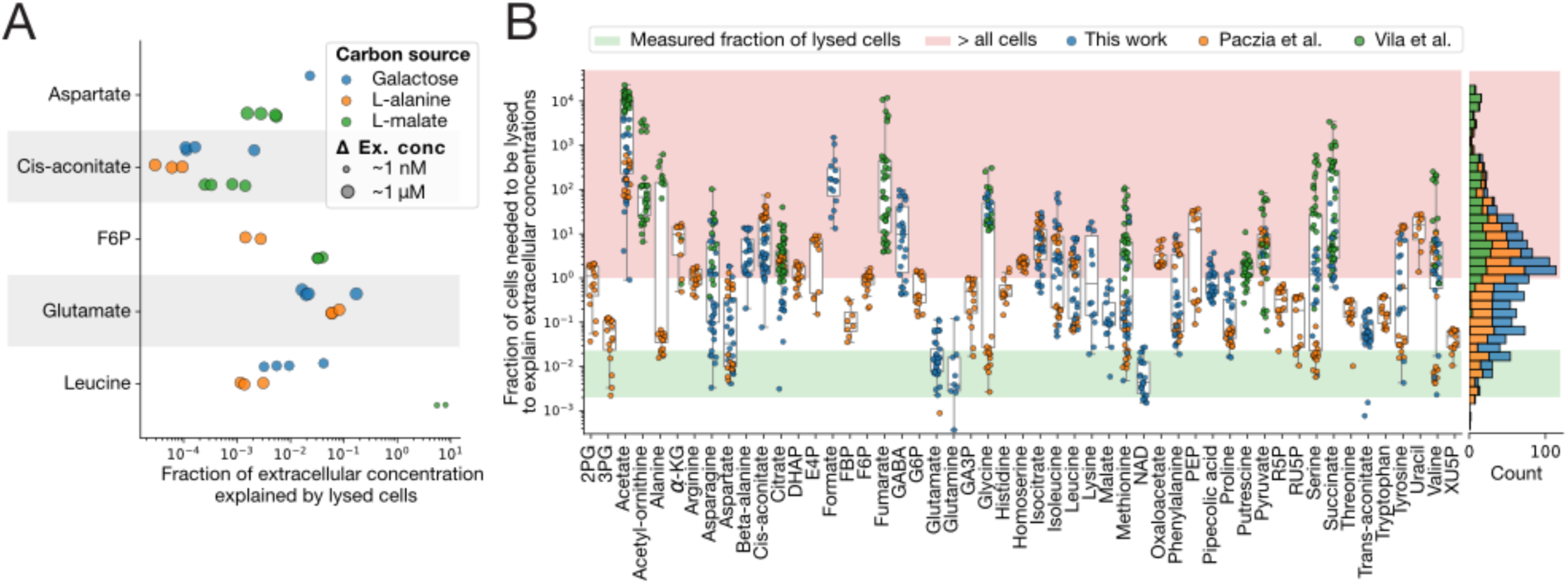
Contribution from cell lysis and associated release of intracellular metabolites to extracellular concentrations. A) Using paired intra- and extracellular metabolite samples from the exponential phase of *E. coli* batch cultures, we quantified the fraction of extracellular metabolite concentrations that can be explained by cell lysis. Point size reflects order-of-magnitude increase in extracellular concentration from the 2-hour sample to the late-exponential phase sample (OD_600_ ≈ 1.1), and all values except leucine in the L-malate condition were in the μM range. B) We expanded this analysis to all metabolite measurements in our *E. coli* datasets by using literature values for intracellular concentrations (Methods). For each metabolite, we estimated the fraction of cell population that would need to be lysed for the intracellular concentrations released by cell lysis to explain the measured extracellular concentrations. The green region covers the range of measured fractions of lysed cells from A), and the red region is where more than the whole cell population is required to explain extracellular concentrations. Abbreviations: 2PG: 2- phosphoglycerate, 3PG: 3-phosphoglycerate, α-KG: alpha-ketoglutarate, DHAP: dihydroxyacetone phosphate, E4P: erythrose 4- phosphate, FBP: fructose 1,6-bisphosphate, F6P: fructose 6-phosphate, G6P: glucose 6-phosphate, PEP: phosphoenolpyruvate, R5P: ribose 5-phosphate, RU5P: ribulose 5-phosphate, XU5P: xylulose 5-phosphate.

Degradation of cell debris may also contribute to extracellular metabolite pools^57^. In *E. coli,* this degradation is dependent on the presence of Lon protease in the cell lysate^58^. However, the benefit of cell debris degradation by Lon was only significant well beyond exponential phase, and only 19 ± 6% of oligopeptides (6-10 amino acids) in the lysate were catabolized by Lon over 20 hours^58^. Thus, in our dataset in which 90% of the samples were collected before 26 hours (67% in exponential phase), we expect only a fraction of the proteome from lysed cells to contribute to the extracellular concentrations of amino acids. When we included 10% proteome degradation in our analysis, only the extracellular levels of tryptophan and threonine additionally became fully explained by cell lysis (Fig. S19). However, apart from amino acids, the metabolites in Fig. 3B are not typical degradation products. Overall, these results suggest that cell lysis and proteome degradation are insufficient to explain extracellular levels of most metabolites – consistent with previous work^12,32^.

### Metabolic disruption and evolution change metabolite release rates

Given that gene deletions substantially perturb intracellular metabolite concentrations^59^, we reasoned that the lack of negative correlation between metabolite value and release rate in *C. glutamicum* (Fig. 1D) could be a result of its engineered metabolism^48^. We hypothesized that modifying central metabolic fluxes in *E. coli* would systematically alter metabolite release rates. To test this and quantify the sensitivity of metabolite release patterns to modifications in core metabolism, we profiled six *E. coli* knockout (KO) strains targeting distinct central pathways (Fig. 4, A and B) with demonstrated effects on intracellular metabolite pools^59^ and predicted impacts on intracellular flux patterns (Fig. S20). In line with our expectations, 13-44% (11-37 of 85) of the late-exponential extracellular metabolite concentrations differed significantly from the wild type across the KO strains (Welch’s t-test, FDR < 0.05, Fig. 4, C and D). In contrast, deleting *lacA* – a negative control not involved in central carbon metabolism – altered only 3.5% of metabolite concentrations (Fig. 4C). Each KO strain displayed a distinct exometabolome profile (Fig. 4E), with the most pronounced differences often observed for metabolites proximal to the deleted reaction (Figs. S21-S23), consistent with intracellular effects of enzyme-deletions^59^. The WT reference for these KO strains from the KEIO collection^60^ – *E. coli* BW25113 – had release rates largely comparable to those of *E. coli* MG1655 used in the experiments presented in Fig. 1 (Fig. S24).

**Figure 4:**
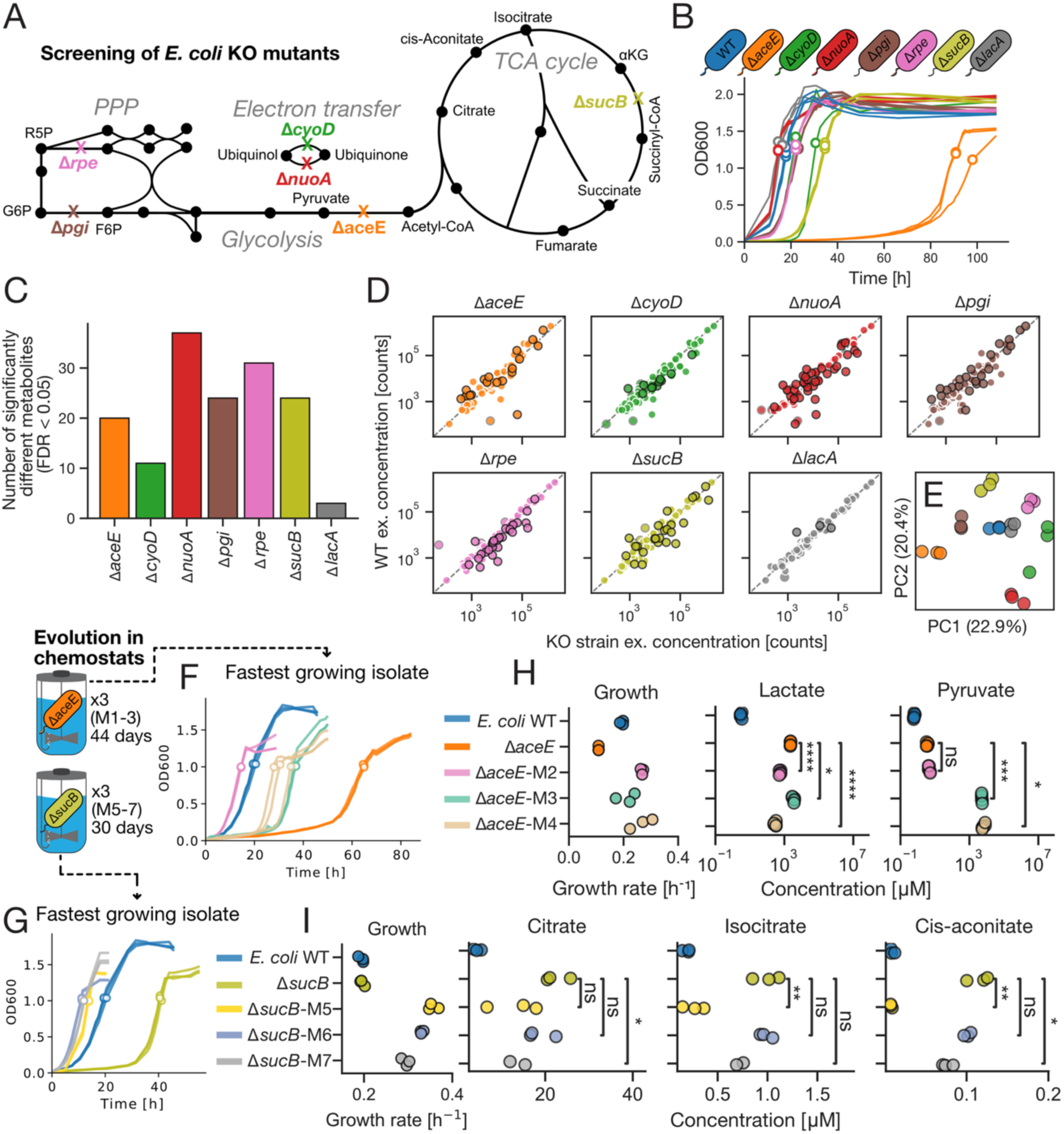
Effect of gene knockouts on metabolite release rates and the effect of subsequent evolution to higher fitness. A) A simplified metabolic network showing the location of the different gene knockouts associated with each strain. B) Growth curves of selected KO mutants, with sampling time-points indicated by open circles. C) The number of significantly different extracellular metabolite concentrations (late exponential phase, OD_600_∼1.2-1.3) compared to WT. D) Comparison of extracellular metabolite concentrations in late exponential phase between KO mutants and WT. Significant differences are marked by solid black outlines (Welch’s t-test, FDR < 0.05). Metabolites not detected in both strains are not shown. E) A PCA plot of exometabolome profiles, showing that each strain has a distinct pattern. F and G) Batch culture growth curves and sampling timepoints for isolates from the endpoint of the evolution experiment for Δ*aceE* and Δ*sucB*, respectively. H and I) Growth rates and extracellular metabolite concentrations for metabolites near the deleted reaction, for Δ*aceE* and Δ*sucB*, respectively. Statistical significance was assessed using Welch’s t-test.

Our central hypothesis for the negative correlation between metabolite value and release rate is that bacteria face a trade-off: the cost of preventing metabolite release versus the energy lost through that release. If this trade-off is under strong selection, then disrupting the balance (e.g. via a gene knockout) should trigger rapid evolutionary compensation to restore it. To test this, we evolved the Δ*aceE* and Δ*sucB* mutants – chosen for their severe growth defects and distinct exometabolome characteristics – for ∼100 (replicates M2–M4) and ∼200 generations (M5–M7) in galactose medium, respectively (Fig. S25). We evolved the strains in chemostats, which select for faster growth at limiting resource concentrations through either increased substrate affinity or increased maximum growth rate^61^. Because release of costly metabolites wastes resources and thereby reduces growth rate, we expected such release to be selected against. Chemostats also avoid the temporal dynamics of batch cultures, where metabolites released during exponential growth can be re-consumed in stationary phase, potentially selecting for efficient scavengers rather than reduced release. The dilution rate was adjusted several times during the first half of the experiment in response to changes in growth dynamics and in an attempt to increase the number of generations, but was kept constant for long periods, including the full second half of the experiment (Fig. S25B). From each chemostat endpoint, we isolated three strains and selected from each replicate the fastest growing clone (Fig. 4, F and G). These clones showed 94-146% (Δ*aceE*) and 50-82% (Δ*sucB*) growth rate increases relative to their ancestors (Fig. 4, H and I). To assess changes in metabolite release, we measured extracellular concentrations of five metabolites per strain at the end of exponential growth (Fig. 4, F and G). These metabolites were selected because their release was strongly perturbed in the ancestors (Figs. S21-S23). We targeted both metabolites proximal to the deleted reaction (pyruvate and lactate for Δ*aceE*; citrate, isocitrate and *cis*-aconitate for Δ*sucB*) where we expected the strongest effects, and metabolites across different chemical classes located elsewhere in the metabolic network to probe more widespread effects. Lactate levels were significantly reduced in 2/3 Δ*aceE* isolates (Fig. 4H), which also exhibited the highest growth rates. The reductions were, however, not sufficient to reach WT concentrations. In contrast, all three Δ*aceE* isolates showed increased pyruvate release, contrary to expectations. These results were qualitatively invariant to growth rate normalization (Fig. S26). For the Δ*sucB* isolates, we found in general small reductions in extracellular concentrations (Fig. 4I), but some of these differences changed qualitatively upon growth rate normalization (Fig. S27). For the metabolites elsewhere in the metabolic network, the outcomes were variable (Figs. S28 and S29).

To estimate the fitness contribution of reducing metabolite release, we used ecGEMs to predict relative growth rate penalties associated with the observed release rates, using the ancestors’ mean growth rates as references. In the Δ*aceE* ancestor, lactate release is predicted to reduce fitness by 23% whereas pyruvate release has negligible impact (0.02%). The reduced release of lactate in the isolates from chemostats M2 and M4 is predicted to reduce the negative fitness effect to only 16% and 7%, respectively. These contributions are small compared to the observed fitness increase (94-146%). The pattern is similar for the Δ*sucB* isolates: the estimated net fitness reduction due to the release of citrate, isocitrate and *cis*-aconitate in the ancestor is only 0.37%, far from sufficient to explain the improved growth rates.

To better understand the reasons for increased growth and changed release rates, we identified sequence variants in the selected fast-growing strains and mutations that had fixed in the chemostat populations (Tables S1–S4). Aside from isolate Δ*aceE*-M4, which likely hypermutated due to a *mutT* mutation, the evolved Δ*aceE* (M2–M4) and Δ*sucB* (M5–M7) isolates harbored few sequence variants. Recurrent mutations were associated with either galactose uptake (e.g. the regulator *galS* in M2, M4 and M7) or the modulation of fluxes proximal to the deleted reaction (e.g. *poxB* in M2; *ldhA* in M4; and *icd, aceK* and *sdhA* in M5–M7). Fixed non-synonymous variants in the chemostat populations supported these observations; for example, *ldhA* and *galS* fixed in M2 and *ilvG* fixed in M3. The Δ*sucB* populations harbored many more fixed mutations, and *galS* and *aceK* were again identified among those shared across replicates. Mutations in *galS*, which encodes a repressor of the high-affinity mgl galactose transporter, are frequently selected in *E. coli* evolved in glucose-limited chemostats^61^, suggesting that selection in our experiment acted largely on substrate uptake.

Together, these results show that metabolite release rates do change as microbes evolve, but the outcome only partially aligns with our expectation that the release of costly metabolites would be reduced. Our interpretation of this mismatch is that selection favored mutations affecting more dominant drivers of fitness, such as substrate uptake, whose benefits outweighed the comparatively small fitness gains from reduced release – leading to inconsistent changes in metabolite release across replicates.

### Transporters cause and counteract metabolite release

Our findings so far suggest that microbes have evolved to release metabolites at rates that scale inversely with their value and that release rates are sensitive to changes in intracellular fluxes. However, the role of metabolite transporters in regulating metabolite release remains unclear. To assess the underlying mechanism, we profiled the exometabolomes of 67 *E. coli* KO mutants from the KEIO collection, each strain devoid of a gene associated with active or passive transport across the inner or outer membrane (Table S5). These genes were selected to span the diversity of transporters, covering major functional classes (ABC, PTS, symporter, antiporter, porin, and channel), a variety of substrates (amino acids, sugars, nucleosides, carboxylic acids, and others), and expected direction (import, export, or both). We cultivated each KO mutant and the WT in glucose minimal medium and quantified changes in extracellular metabolite concentrations over the course of batch growth using flow-injection analysis time-of-flight mass spectrometry (FIA- TOF MS)^62^. The final dataset comprises 2277 unique samples collected at 10 timepoints and analyzed in quadruplicate, yielding 9089 mass spectrometry measurements after quality control. We could annotate *m/z* ion peaks to 333 metabolites in the *E. coli* metabolome. Four strains were excluded from downstream analyses due to severely impaired growth or failed strain quality control (Methods). We quantified the *effect* of each transporter on each metabolite as the ratio between the mean KO-WT difference and the within-strain variability (Fig. 5A). Statistical significance for all 15848 strain-metabolite *effects* was assessed using a permutation test (Methods). The dataset is made available for interactive exploration at <u>keio.unil.ch</u>.

**Figure 5:**
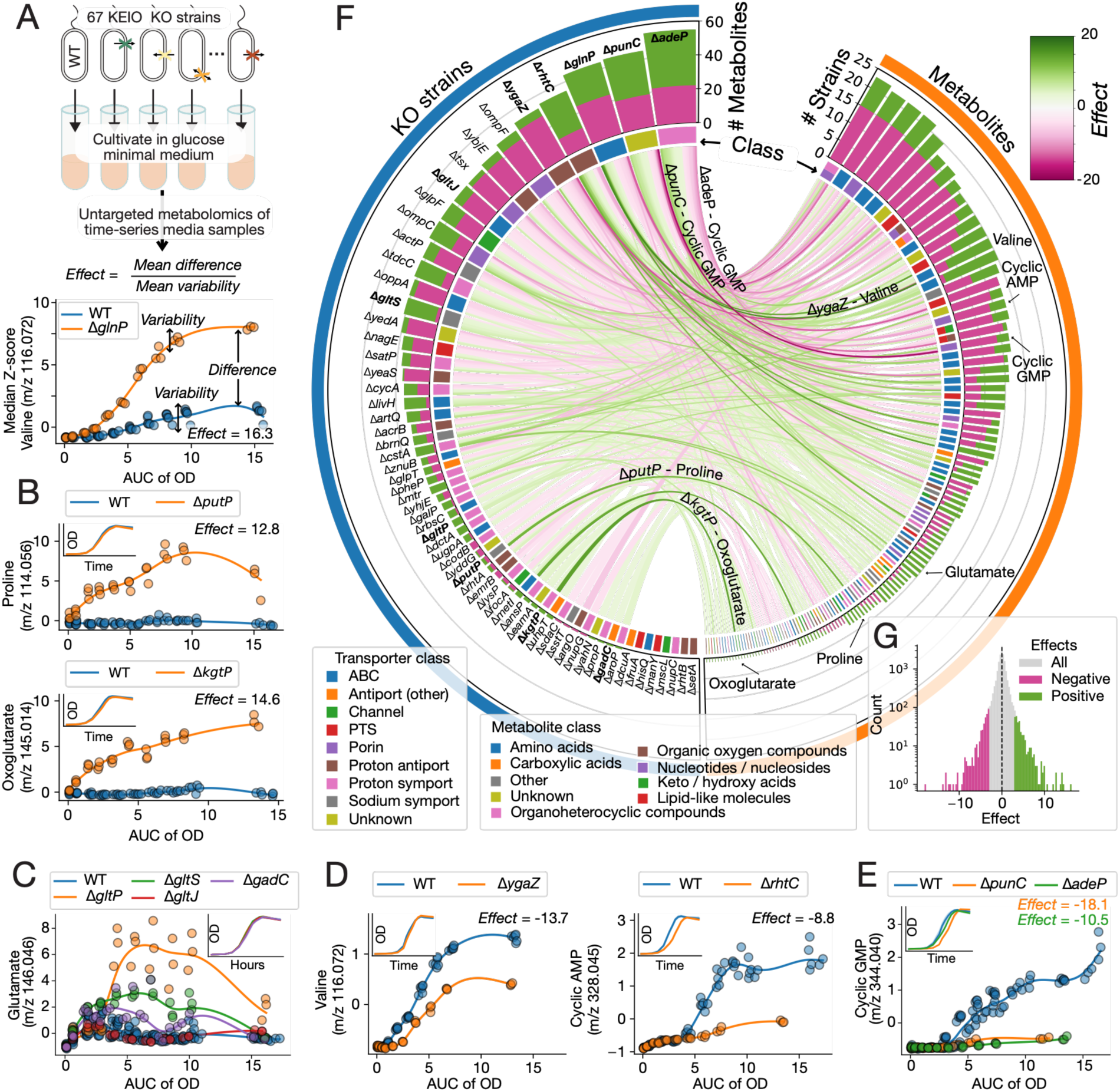
Transporters cause and counteract metabolite release. A) Approach: We screened 67 *E. coli* KO strains in batch culture and analyzed the temporal dynamics of extracellular metabolites for the 63 strains retained after quality control (Methods). We quantified the effect of each strain on each metabolite by computing the ratio between the mean KO-WT difference and the within-strain variability (Methods). B-E) Each panel shows the extracellular concentration of one metabolite (Z-score, median of four technical replicates) for *E. coli* BW25113 (WT) and 1-4 KO mutants across a batch culture, with the area under the curve of the optical density (AUC of OD) on the x-axis. Small inset panels compare the growth curves of the respective strains (mean OD, three technical replicates). F) Overview of significant, large effects (|*Effect*| > 3, *P* < 0.005, permutation test). The left bar chart displays the number of positive (green) and negative (pink) large effects across all 63 KO strains, and the color label indicates the class of the deleted transporter. The right bar chart displays the number of positive (green) and negative (pink) effects on each of the metabolites affected in one or more of the KO strains (122 in total, |*Effect*| > 3, *P* < 0.005, permutation test). The color label indicates the classification of all metabolites in *E. coli* mapping to the corresponding m/z value, which in some cases maps to two different classes. The chord diagram displays which metabolites are affected (and the strength) in each KO strain. Metabolites / strains displayed in panels B-E are annotated and highlighted in bold, and the strongest effects are also annotated on the links in the chord diagram. G) Distribution of all effects across all KO strains and metabolites.

A positive effect implies that removing a transporter increases the net release of the respective metabolite in the KO strain compared to the WT, while a negative effect implies the opposite. We found a strong positive effect in 2.2% (343/15848) of the strain-metabolite pairs (*Effect* > 3*, P* < 0.005). In many cases, the strongest effect matched the annotated function of the transporter (Fig. 5B). For example, deletion of the high-affinity proline/sodium symporter gene *putP* strongly increased proline release, suggesting that PutP counteracts proline release in the WT by re- importing proline lost via other mechanisms. This mechanism has already been described for *E. coli* strains selected for proline secretion^63^. Similarly, deletion of the oxoglutarate importer *kgtP* most strongly increased oxoglutarate release. For glutamate, three of the four annotated transporters exhibited positive effects, suggesting that they collectively counteract its release (*P*

< 0.005, Fig. 5C). This active re-import potentially explains why glutamate is released at moderate, and sometimes not significant, rates despite its high intracellular concentration. However, we also observed strong and sometimes diverse effects extending beyond the metabolites annotated to the respective transporters in EcoCyc^64^. For example, deletion of the glutamine importer *glnP* strongly increased the extracellular concentration of 20 metabolites (*Effect* > 3, *P* < 0.005), including valine, hydroxybutanoic acid, aminobutanoic acid, isoleucine/leucine, alanine, and glutamine (Fig. S30).

Strong negative effects were found in 2.4% (379/15848) of the strain-metabolite pairs (*Effect* < - 3, *P* < 0.005), showing that transporters frequently cause metabolite release. Some of these effects were linked to transporters primarily associated with metabolite export, such as YgaZ, which exports valine, and RhtC, which mediates threonine efflux (Fig. 5D)^65,66^. However, we observed equally strong effects for transporters that are important for metabolite uptake such as the purine and adenosine transporters PunC and AdeP, respectively (Fig. 5E)^67–69^. Similarly, the “exporters” YgaZ and RhtC showed positive effects comparable in magnitude to their negative ones (Fig. S30).

To test whether the measured effects of transporter deletion on metabolite release are ecologically relevant, we co-cultivated the WT and 2–4 KO strains with a proline, methionine or isoleucine auxotroph. Strains with measured positive effects on amino acid release generally supported auxotroph growth better than WT (4/5 significantly). Of the two strains with negative effects, Δ*tsx* did not differ significantly from the WT. Unexpectedly, Δ*ygaZ* supported growth of the isoleucine auxotroph better than the WT, despite its negative effect on isoleucine/leucine release (Fig. S31). Altogether, these results suggest that the transporter-mediated metabolite effects identified in our screen can translate into ecologically relevant outcomes for microbial interactions, but further experiments are needed to understand when such effects are predictive.

Among the 63 KO strains, 55 strongly influenced one or more metabolites (|*Effect*| > 3, *P* < 0.005, Fig. 5F). Of these, 29 strains strongly influenced 10 or more metabolites, and 37 showed both positive and negative effects. While the balance between positive and negative effects differed between strains, the overall dataset showed a striking symmetry with almost equally strong and numerous positive and negative effects (Fig. 5G). From the metabolite perspective, 122 metabolites were affected by one or more strains, and 57 of these were affected in both directions (|*Effect*| > 3, *P* < 0.005, Fig. 5F). We found no significant differences in the distribution of effect sizes across metabolite classes, nor any correlation between the value of a metabolite and how frequently it was affected by the deletion of a transporter (Fig. S32). Likewise, effect sizes were similarly distributed across transporter classes, and neither the number of affected metabolites nor the maximum effect size correlated with average transporter expression levels (Fig. S33)^70^.

These findings show that transporters are highly promiscuous, driving diverse metabolic changes^34,71^. We were unable to draw a clear distinction between exporters and importers, as most transporters featured both positive and negative effects. The resulting complexity of strain- metabolite connections suggests that trade-offs are common. Taken together, our results demonstrate that transporters expressed in *E. coli* are both a cause of metabolite release and a mechanism to counteract it. We suggest that *E. coli* has evolved to combine the activity of these importers and exporters to regulate the net release rates of different metabolites according to their value.

## Discussion

To better understand metabolic interactions between microbes, it is necessary to develop and test hypotheses for the fundamental principles that govern them. Here we propose that the broad release of intracellular metabolites is in general a loss of energy and that for each metabolite there is a trade-off between the cost of its release and the cost of preventing it. This hypothesis connects to previous work linking fitness costs to microbial traits, including amino acid uptake rates^14^, amino acid use and proteome composition^10,46^, interaction strengths between *E. coli* auxotrophs^10^, properties of extracellular enzymes^72^ and metabolic strategies^73^.

We find consistent support for our hypothesis: (i) metabolite values are negatively correlated with metabolite release rates across different species and nutrient environments, (ii) metabolite value explains the most variability in release rates among all tested factors, (iii) release rates cannot be explained by cell lysis alone, and (iv) knock-out of metabolite-transport genes frequently changes net metabolite release, providing a mechanistic explanation for the observed release and for how microbes can act to reduce the net release of valuable metabolites. We emphasize that our measurements reflect net release rates, i.e. the balance of gross release and recapture, and that the negative correlation with metabolite value may therefore reflect selective investment in recapture of costly metabolites, rather than reduced gross secretion per se, or both. Also, our metabolite value estimates do not account for costs associated with recapture, so quantifying gross and net release rates and the associated cost of metabolite transport is a natural direction for future work. A final caveat is that all our growth assays were performed in monoculture. Metabolite release rates could differ in the presence of other species – for example, when reciprocal cross-feeding makes increased release beneficial to the producer itself^74,75^ – which could affect the patterns observed here. However, our results suggest that such cooperative interactions are rare and that release rates have primarily evolved to benefit the metabolite producer.

The complexity of KO strain–metabolite interactions (Fig. 5F) suggests why eliminating metabolite release entirely may be challenging, given the apparent trade-off between release and recapture, and the need to express several transporters at basal levels to sense and adapt to environmental changes^76–78^. Redundant transporters and competing influx and efflux similarly govern xenobiotic accumulation in *E. coli*^79,80^, highlighting the widespread nature of such transport complexity. The high frequency of metabolites with both significant positive and negative effects suggests that futile cycles might be common and raises the question of whether such cycles contribute to intracellular homeostasis, extending beyond their previously demonstrated role in metal ion regulation^81^. Furthermore, we find that release rates are sensitive to changes in intracellular metabolism caused by both gene knockouts and subsequent evolution. Yet on the timescale of our evolutionary experiments, changes in release rates were inconsistent across replicates and did not consistently reduce the release of costly metabolites. This may reflect alternative evolutionary paths that improved fitness independently of metabolite release, such as improved substrate uptake.

The large-scale exometabolomics screen of *E. coli* transporter knockout strains provides a unique resource for refining transporter annotations in databases and metabolic models, although the complexity might be challenging to distil. In parallel, our finding that simple linear models based on metabolite value can predict order-of-magnitude release rates offers a new avenue for such models to forecast microbial interactions. More broadly, our findings reshape how we think about metabolite release and the evolution of microbial cross-feeding. While it is commonly assumed that cross-feeding either emerges from extracellular enzymes or from the release of waste products^2,82,83^, our results suggest an alternative: microbes release metabolites because using, retaining or immediately recapturing them is not worth the investment. This increases the scope for commensal cross-feeding, which in this scenario can arise without invoking the evolution of cooperation^84^ and without being vulnerable to cheaters^36^.

## Acknowledgements

We thank all members of the Mitri lab at the University of Lausanne for discussions and feedback. We also thank people at the Department of Biotechnology and Nanomedicine at SINTEF for technical support and advice on the experimental design, and Vassily Hatzimanikatis and the Laboratory of Computational Systems Biotechnology at EPFL for feedback on the computational analyses. Furthermore, we would like to thank Simon van Vliet, Olga Schubert, Guilhem Panneau, Jean C. C. Vila, Milton Saier and Uwe Sauer for providing feedback, Olga Schubert and Kim Schlegel for providing the KEIO strains, Sebastian Burz, You Zheng Teo and Clément Vulin for support with the live/dead staining protocol, Estelle Pignon for providing *E. coli* auxotrophs and Vladimir Sentchilo for support with the flow cytometry and sampling for metabolomics analyses. We also thank the Metabolomics unit at UNIL, and specifically Hector G. Ayala and Julijana Ivanisevic, not only for conducting the LC-MS analyses, but also for providing guidance and recommendations on related matters. Special thanks to Florent Mazel and Daniel Segrè for feedback on the draft manuscript. Finally, we thank Lilja B. Thorfinnsdottir, Jean C. C. Vila and Stephan Noack for sharing of experimental data.

## Funding

Swiss National Science Foundation Swiss Postdoctoral Fellowship TMPFP3_217172 (SS)

National Center of Competence in Research Microbiomes grant SNF 51NF40_180575 (SM, SS, AG, MAV, JL, PP)

Faculty of Biology and Medicine, University of Lausanne (EU)

Swiss National Science Foundation Eccellenza grant PCEGP3_181272 (SM, MAV) SINTEF Industry (GB)

Research Council of Norway, SFI Industrial Biotechnology, grant no 309558 (GB)

## Author contributions

Conceptualization: SS, SM; Methodology: SS, PD, AQ, JSL, GB, MAV, EU; Investigation: SS, GB, PD, AG, EU, PP; Visualization: SS, PP, EU; Funding acquisition: SS, SM, PE; Project administration: SS, SM; Supervision: SS, SM; Writing – original draft: SS; Writing – review & editing: All authors

## Data and materials availability

Strains and isolates used in this study are available on request. Formatted data and code for data analyses and visualization are available on GitHub at https://github.com/Mitri-lab/metabolite-release. A permanent archive of this repository has been deposited to Zenodo (<u>10.5281/zenodo.21764846</u>). The raw sequencing data and metadata have been deposited to the NCBI SRA database under the BioProject ID PRJNA1270783. The raw FIA-TOF MS data are available on MassIVE under the dataset identifier MSV000097105.

## Supplementary Information

Supplementary Figures S1-S34 Supplementary Tables S1-S7

## Methods

### Strains

All experiments were conducted with either *E. coli* K-12 MG1655, *E. coli* K-12 BW25113 (wild-type ancestor for KEIO KO strains) or one of the 74 KO mutants selected from the KEIO collection^60^ (Table S6). The KEIO KO strains were verified by PCR with one primer binding inside the Kan^R^ cassette inserted during gene knock-out (Keio_K1_Rev for genes on the (-) strand; Keio_KT_RvComp_Fw for genes on the (+) strand) and one gene-specific primer binding downstream of the knocked-out gene (primers are listed in Table S6)^60,85^. For all experiments, the KEIO KO strains were used with the kanamycin resistance cassette in place.

### Precultures and growth media

For all experiments detailed below, the strains were precultured as described here unless otherwise stated: From glycerol stocks stored at -70 °C each strain was streaked out onto LB agar plates and incubated overnight (37 °C). Then, one colony of each strain was picked and used to inoculate liquid precultures (10 mL LB in 50 mL Erlenmeyer flasks) that were incubated overnight (37 °C, 200 rpm). The kanamycin-resistant KEIO KO strains and strains evolved from these were always precultured in liquid LB and streaked out onto LB agar plates with 25 µg/mL kanamycin, while *E. coli* K-12 MG1655 and *E. coli* K-12 BW25113 were always precultured in LB and streaked onto LB agar plates without selection. Unless otherwise noted, precultures were washed three times by centrifugation, with the supernatant removed after each step, and the cells resuspended in M9 medium with no carbon source to eliminate residual LB before inoculation. Unless otherwise stated, M9 medium was always prepared by mixing (per liter): 200 mL 5X M9 salts (M6030, Sigma-Aldrich, St. Louis, MO, USA), 2 mL 1 M MgSO_4_, 0.1 mL 1 M CaCl_2_, 1 mL 0.5 g/L FeSO_4_·7H_2_O, MQ water and appropriate amounts of a carbon source stock solution. The final M9 medium was then pH-adjusted to pH = 7.0, filter sterilized (0.22 μm) and stored at 4 °C until use.

### Selecting carbon sources for *E. coli* K-12 MG1655 batch cultures

To complement the existing exometabolome data from batch cultures of *E. coli* on glucose^12^, we aimed to select three carbon sources that differed in chemical class and caused variation in metabolic flux patterns. We used the ecGEM eciJO1366 of *E. coli* K-12 MG1655 without any modifications and parsimonious FBA (pFBA) to simulate optimal flux patterns across 104 different growth supporting metabolites^42,86^. We then performed a PCA of the predicted fluxes (as Boolean values) to compare flux patterns between the carbon sources (Fig. S5). We used HDBSCAN^87^ to identify clusters and selected from the different clusters in total 10 compounds that also differed in chemical class (sugars, amino acids and organic acids) and their entry point into central carbon metabolism (Fig. S5). We then screened *E. coli* growth on the selected carbon sources (plus glucose as a positive control) in triplicates over 48 hours in 96-well plates (Fig. S5). For this experiment, we prepared M9 medium with each of the 11 different carbon sources to a concentration scaled to 120 mM carbon atoms. From washed LB precultures, *E. coli* K-12 MG1655 was inoculated to OD_600_ = 0.05 in 200 μL of medium and cultivated with continuous shaking (double orbital, 425 cpm) in a BioTek Synergy H1 (Agilent Technologies, Winooski, VT, USA) plate reader at 37 °C for 48 hours. We ultimately chose galactose, L-alanine and L-malate, as they supported rapid growth, differed in chemical class, entered central metabolism at different locations and were grouped into different clusters based on flux patterns (Fig. S5).

### *E. coli* K-12 MG1655 batch cultures in bioreactors with different carbon sources

*E. coli* K-12 MG1655 was cultivated in lab-scale bioreactors using the three selected carbon sources. Their concentrations were normalized by the number of carbon atoms to a carbon atom concentration of 120 mM, corresponding to 20 mM galactose (G0625, Sigma-Aldrich), 30 mM L- malic acid (02290, Sigma-Aldrich) and 40 mM L-alanine (A7627, Sigma-Aldrich). M9 media supplemented with each of these three carbon sources were prepared as described above using 25X stock solutions.

The strain was streaked from a frozen glycerol vial onto an LB agar plate and incubated at 37 °C overnight. One colony was transferred to a 250 mL baffled shake flask containing 40 mL LB medium. After 7 hours, a culture volume corresponding to a start OD_600_ = 0.05 was used to inoculate a 500 mL baffled shake flask containing 100 mL M9 + 40 mM L-alanine. This flask was incubated for 36 h (200 rpm, 37 °C). For the two other carbon sources, the strain was incubated in LB for 17 h before inoculating a culture volume corresponding to OD_600_ = 0.05 in a 500 mL baffled shake flask containing 100 mL M9 + 20 mM galactose or M9 + 30 mM L-malic acid, respectively. The flasks were incubated for 24 h (200 rpm, 37 °C). Before inoculation to bioreactors, M9 precultures were centrifuged (3220 rcf, 10 min, 4 °C), supernatant removed, and pellet resuspended in M9 medium without carbon.

Batch cultivations were performed in 1 L DASGIP bioreactors (Eppendorf DASGIP, Jülich, Germany) with an initial volume of 600 mL M9 medium supplemented with either 20 mM galactose, 30 mM L-malic acid or 40 mM L-alanine, three replicates per condition. The reactors were equipped with two Rushton impellers and probes for measurements of dissolved oxygen (DO) and pH. Submerged aeration was maintained at 0.5 vvm, and agitation was cascaded between 400 and 800 rpm, controlled by a DO set point of 30%. pH set point was 7.0 and controlled using 2 M NaOH and 2 M HCl. Start OD_600_ was 0.05.

Sampling was performed regularly during the exponential growth phase with additional samples collected during stationary phase. OD_600_ was measured at all time points and supernatant stored: 2 mL culture was centrifuged (3220 rcf, 10 min, 4 °C), and supernatant frozen at -20 °C. At the end of exponential phase and at the end of the experiment, samples were collected for cell dry weight (CDW). 50 mL culture was centrifuged (3220 rcf, 10 min, 4 °C), supernatant removed, cell pellet resuspended in 50 mL MQ water, centrifuged, supernatant removed, cell pellet resuspended in 5-10 mL MQ water and transferred to a pre-weighed aluminum beaker and dried at 105 °C for 24 hours. Beakers were then weighed again and CDW calculated.

### Exometabolome analyses of time-series samples from *E. coli* K-12 MG1655 batch cultures with different carbon sources

Four samples from each of the nine batch cultures (three replicates of three different media) and media samples were analyzed with LC-MS and GC-MS to measure the extracellular concentration of metabolites at different time points throughout the experiment. The time points were selected based on time-series cultivation data to cover the exponential growth phase (three samples) and one stationary phase sample (Fig. S6). Samples from the different replicates were aligned based on the timing of the end of the exponential growth phase (Fig. S6).

LC-MS of selected bioreactor samples was conducted by the Metabolomics facility at UNIL using their method “high-coverage targeted analysis of polar metabolites”, following this procedure: Media samples were thawed on ice. An aliquot (20 µL) was extracted with the addition of ice-cold methanol (80 µL), vortexed (30 sec) and centrifuged (15000 rpm, 15 min, 4 °C). The supernatants were transferred to vials for liquid chromatography – tandem mass spectrometry (LC-MS/MS) analyses. The extracts were analyzed by hydrophilic interaction chromatography coupled to a tandem mass spectrometer (6496 iFunnel, Agilent Technologies) using multiple reaction monitoring - MRM approach in both positive and negative ionization modes, to maximize the polar metabolome coverage. The analytical conditions have been described in detail elsewhere^88,89^. Raw LC-MS/MS metabolome data were processed using the MassHunter Quantitative analysis software (Agilent Technologies, RRID:SCR_015040). The peak areas (or extracted ion chromatograms (EICs) for the monitored MRM transitions) were used for relative comparison of metabolite levels between different conditions or groups of samples. Data quality assessment, including signal drift correction, was performed using pooled quality control (QC) samples analyzed periodically throughout the entire batch.

Metabolite quantification was performed using stable isotope labeled internal standards and calibration curves following the signal drift correction. Data processing was done using MassHunter Quantitative analysis software. Peak area integration was manually curated, and concentrations were reported by selecting a 6-point portion of the linear standard curves relevant for the observed concentrations.

The relative LC-MS data were standardized before subsequent analyses and data processing. Both the relative and absolute LC-MS data were cleaned of obvious outliers. For the absolute data the outliers were caused by carry-over from previous samples leading to too high malate (six samples) and succinate (one sample) values. For the relative LC-MS data there were additional outliers for unknown reasons (39 values in total) affecting 14 metabolites in total. For several metabolites, there were more missing values (not detected/not quantified) in the absolute data than in the relative data. For a few metabolites (phenylalanine, proline, creatine), we leveraged good linear relationships between relative and absolute data (*R*^2^ > 0.8) to estimate the missing absolute values using a simple linear transformation.

GC-MS analysis of the same samples was conducted in the following way: Analytical standards were prepared as M9 medium with increasing concentrations of acetate, lactate, pyruvate, formate and propionate and stored at -70 °C until use. Samples from the *E. coli* bioreactor batch cultures and analytical standards were thawed on ice. Then, 100 μL was transferred to an Eppendorf tube and kept on a cold block (-20 °C) while adding 5 μL 11% HCl and 500 μL diethyl ether. Tubes were capped, vortexed on a thermoblock (2000 rpm, 10 min, 1 °C) and centrifuged (13000 rcf, 5 min, 4 °C). The top organic phase layer was pipetted to a glass vial and derivatized with 20 μL N-(t-butyldimethylsilyl)-N-methyltrifluoroacetamide (MTBSTFA) and vortexed briefly. Samples were then placed on a heat block (90 min, 35 °C) and kept at 16 °C until analysis. The samples were injected (1 μL) by a Pal3 autosampler onto an Agilent 8890-5977B GC-MSD (Agilent Technologies) with a VF-5MS (30 m x 0.25 mm x 0.25 μm) column. The samples were injected with a split ratio of 15:1, helium flow rate of 1 mL/min and inlet temperature of 230 °C. The temperature was held for 2 min at 50 °C, raised at 25 °C/min to 175 °C, 30 °C/min to 280 °C and held for 3.5 min. The MSD was run in scan mode from 40-500 Da. Analyte abundances were calculated using the MassHunter Quantitative analysis software. Absolute concentrations were calculated using standard curves.

Galactose concentrations were measured using the L-Arabinose/D-Galactose assay kit (K-ARGA, Megazyme, Bray, Ireland), following their rapid protocol for analyses in 96-well plates.

### Estimation of metabolite release rates of *E. coli*, *B. licheniformis*, *C. glutamicum* and *S. cerevisiae* in glucose medium

Time-series data on biomass (in OD_600_) and absolute extracellular metabolite concentrations from batch cultivations of *E. coli, B. licheniformis, C. glutamicum* and *S. cerevisiae* in high glucose concentrations (10-20 g/L)^12^ was kindly provided by the authors (mean and standard deviations). For metabolites reported as a sum of two metabolites, we attributed equal amounts to those metabolites to enable direct comparison with metabolite values estimated for individual metabolites (described below). This includes 2-phosphoglycerate/3-phosphoglycerate and ribulose 5-phosphate/xylulose 5-phosphate. To estimate the metabolite release and uptake rates of each species, we first converted the OD_600_ values to gDW/L. For *E. coli* we used the conversion factor we estimated from our own cell dry weight measurements in M9 galactose medium (0.346 gDW/L/OD_600_), which is comparable, but on the lower end of values found in the literature (0.36- 0.515 gDW/L/OD_600_)^90,91^. For the three other species, we used literature values: B. licheniformis (data from B. subtilis) 0.48 gDW/L/OD_600_^92^; *C. glutamicum* 0.27 gDW/L/OD_600_^93^; *S. cerevisiae* (mean of literature values) 0.746 gDW/L/OD_600_^90^. We then integrated the translated biomass time-series data to obtain biomass AUC data (*X_AUC_*(*t*) = ∫*^t^* *X*(*t*)*dt*). Assuming a constant specific net uptake or release rate *a_i_*, extracellular metabolite concentration *C_i_*(*t*) was modeled as a linear function of the integrated biomass: *C_i_*(*t*) = *a_i_* ⋅ *X_AUC_*(*t*) + *C*_0_. We leveraged this linear model together with the absolute exometabolome data to estimate the specific net rate *a_i_* for each metabolite and species using linear regression (Figs. S1–S4). Rates were only estimated for the exponential growth phase, and only until each metabolite either saturated or was clearly being re-consumed.

### Estimation of metabolite release rates of *E. coli* in galactose, L-malate and L-alanine

For our bioreactor batch cultivations of *E. coli* in galactose, L-malate and L-alanine, we used paired cell dry weight measurements and OD_600_ readings to calculate conversion factors used to translate OD_600_ values for all time points (galactose: 0.346 ± 0.016; L-malate: 0.279 ± 0.017; L-alanine: 0.296 ± 0.011; mean ± std in gDW/L/OD_600_). The translated time-series data were used to calculate the biomass area under the curve used to estimate metabolite release and uptake rates further described below.

We estimated the metabolite uptake or release rate by performing for each metabolite and each carbon source a linear regression of measured concentration (relative or absolute) vs biomass AUC using the three samples (timepoints T1-T3) acquired from the three replicate batch cultures during exponential phase plus the initial M9 medium (T0; Figs. S7-S9). Twenty-eight of the 126 unique metabolites were filtered out because they did not map to the *E. coli* Metabolome database (ECMDB)^53^. To avoid underestimating release rates for metabolites that potentially were being re-consumed at the end of the exponential phase, we discarded the last timepoint from the exponential phase in cases where the rate was positive (indicating metabolite release) but there was a significant reduction in extracellular concentration from T2-T3 (*P* < 0.05, one-sided t-test). For 18 (of 98) metabolites, we chose to not include T0 for at least one of the three conditions in the linear regression, either because of very large variability of T0 or because the trend from T0 to T1 was clearly different from the general trend from T1-T3. We only estimated the rate for metabolites where we had at least three data points from at least two different time points. For the metabolites where we had both relative and absolute data, we used the absolute data if there were enough absolute data points to get adequate slope estimates. For seven metabolites (glutamine, alpha-aminoadipate, serine, lysine, lactate, NAD and isocitrate) where both absolute and relative quantification was performed, we could estimate the rate more accurately on the relative dataset (because of more missing data points in the absolute dataset). However, there were sufficient absolute measurements to estimate the absolute spread (standard deviation) of these data points, enabling conversion of the slope value from Z-score to μmol/gDW/h.

### Estimation of metabolite release rates of *E. coli*, *P. putida,* an *Enterobacter* sp. and a *Pseudomonas* sp. in various carbon source environments

The dataset from Vila et al. (2023) was kindly provided by the authors^49^. The dataset contains exometabolome data from three timepoints (16, 28, and 48 hours) of *E. coli* MG1655, *P. putida* KT2440, and two environmental isolates of the genera *Enterobacter* and *Pseudomonas,* respectively^49^. All species were cultivated in M9 medium with one of five carbon sources (D- glucose, D-fructose, glycerol, pyruvate or L-malate). Additionally, the dataset includes exometabolome data from two timepoints (28 and 48 hours) of *E. coli* and the *Enterobacter* sp. on D-ribose, L-arabinose and D-galactose, and of *P. putida* and the *Pseudomonas* sp. on acetate, fumarate and succinate. To estimate metabolite release rates, we first used the plate reader growth curves to calculate biomass AUC after 16, 28 and 48 hours and to identify the end of exponential growth of each organism in each condition. The end of exponential phase was automatically identified as the first peak in the second derivative of the smoothed growth curve with an OD_600_ above 60% of maximum. We converted the AUC from OD_600_ to gDW/L using the same conversion factor for all the different carbon source environments: 0.36 gDW/L/OD_600_ for *E. coli* and the *Enterobacter* sp., mean of measured values and literature values^90,91^; 0.476 gDW/L/OD_600_ for *P. putida* and the *Pseudomonas* sp., mean of literature values^94,95^. Finally, we performed a linear regression between extracellular metabolite concentrations and AUC on both duplicates of each strain/carbon source combination including all datapoints until 2 hours after the end of exponential growth. The 2-hour buffer was included to account for some inaccuracy in the automatic detection. We only included rate estimates for metabolites that were quantified at least twice for each strain. As the dataset did not include an early timepoint or media measurement, we assumed the initial concentration of each metabolite to be 0. If a metabolite was detected at more than one timepoint, we discarded the estimated rate if there was a qualitative discrepancy in slope value when the slope was estimated with or without the T0 concentration assumed to be 0. If the carbon source was among the measured metabolites, this metabolite was excluded from rate estimation in those specific conditions.

### Genome-scale metabolic models

For *E. coli, C. glutamicum* and *S. cerevisiae* we used the previously developed enzyme-constrained genome-scale metabolic models (GEMs) eciJO1366^42^, ecCGL1^96^ and ecYeastGEM 8.3.4^41^, respectively. For *B. licheniformis* there was no available enzyme-constrained GEM, and we therefore opted to use the enzyme-constrained GEM ecBSU1 of the closely related species *B. subtilis*^97^, which was available at <u>github.com/tibbdc/ecBSU1</u>. A normal GEM for *B. licheniformis* is available^98^, but with this model we could not achieve any feasible flux balance solution. To simulate the environmental isolate of the genus *Enterobacter* we used the *E. coli* GEM eciJO1366. For the environmental isolate of the genus *Pseudomonas* and for *P. putida* we used the enzyme- constrained version of the *P. putida* GEM iJN1463^99^ automatically generated by the ECMpy2 method^43^, which we here refer to as eciJN1463.

We made a few minor changes to the published GEMs before predicting metabolite values. In the *E. coli* GEM eciJO1366 the proteome weight of the forward reaction *GALKr* (galactokinase) was extremely high (0.016), limiting the growth rate on galactose to much less than the growth rate we measured in M9 galactose medium. To enable eciJO1366 to reach the measured growth rate with galactose as the only carbon source, we replaced the proteome weight of the forward *GALKr* reaction with the proteome weight of the reverse *GALKr* reaction which was much smaller (0.00037). For the ecYeastGEM we only merged exchange reactions that were previously split to simplify its use. Both ecBSU1 and eciJN1463 were translated from json to sMOMENT format in xml^42^, modified by merging split exchange reactions and fixing metabolite compartment names and annotations. For ecBSU1 we also removed the enzyme constraint on the reaction “PPAm_num1” (inorganic diphosphatase) as it caused unrealistic uptake and release rates of phosphate and diphosphate, respectively. The *C. glutamicum* GEM ecCGL1 was also converted from the ECMpy format^43^ to the sMOMENT format^42^ but otherwise kept unchanged. All GEMs were used with their default biomass equations as objective when maximizing growth in all FBA or pFBA analyses. Reframed 1.5.3 (<u>github.com/cdanielmachado/reframed</u>) or COBRApy 0.29.1 (RRID:SCR_012096)^100^ were used to load, manipulate and run constraint-based analyses with Gurobi 10 (Gurobi Optimization, LLC) as the solver.

### Estimating metabolite value and turnover using genome-scale metabolic models

In constraint-based modeling, a shadow price is an estimate of how sensitive the objective function in a flux balance analysis (FBA) is to increased influx or efflux of a metabolite^101,102^. Hence, one can interpret the shadow price as *metabolite value* as it quantifies the impact on growth caused by the release of a metabolite (to have positive values we here define metabolite value as the shadow price multiplied by -1). A similar approach to measure the cost of metabolite release has been used to identify costless secretions^37^. The metabolite value can be predicted for a specific organism in a specific context by using a GEM of the corresponding organism (or closely related) and then constraining this GEM to the specific context (i.e. primarily defining metabolite uptake rates). While shadow prices are usually estimated directly by the solver used to run FBA^44^, the robustness of these values is often questioned because of numerical issues. To get more robust shadow price estimates we solved for each metabolite in each context a separate FBA, where we constrained the GEM to have an (additional) release of 0.01 mmol/gDW/h of that metabolite and quantified the change in the objective function, i.e. the change Δ*y* in growth rate caused by the forced metabolite release. The metabolite value was then quantified as Δ*y/0.01*. If the metabolite was predicted to be released by FBA, we added 0.01 mmol/gDW/h to the FBA- predicted value to estimate the rate. This was only relevant for acetate for *E. coli* in the L-malate condition, for ethanol, acetate, formate and pyruvate for *S. cerevisiae*, and for pyruvate for *P. putida* in the L-malate condition.

Using this approach, we estimated metabolite values for *E. coli* in batch cultures with glucose, galactose, L-malate or L-alanine as the carbon source by constraining the uptake of the corresponding exchange reaction in eciJO1366 to the uptake rate of the carbon source as estimated from the experimental data. The same approach was used for *C. glutamicum, B. licheniformis* and *S. cerevisiae* using the corresponding models. For *E. coli*, *P. putida* and the two environmental isolates of the genera *Enterobacter* and *Pseudomonas* we first extracted maximum growth rates for each organism in each environment from growth curves (using curveball^103^). Then, we used the respective GEMs to calculate the uptake rates of the corresponding carbon sources required to sustain the maximum growth rates. These uptake rates were then used to constrain the GEMs when estimating the metabolite values as described above. The turnover (i.e. the sum of positive fluxes producing an intracellular metabolite) was estimated by running parsimonious FBA^86^, using the same GEMs constrained in the same way as described above for metabolite values.

To validate that the negative correlation between log-transformed metabolite values and release rates was not dependent on the choice of GEM, we compared estimates from eciJO1366 with estimates from the four benchmark *E. coli* K-12 MG1655 GEMs iJR904^104^, iAF1260^105^, iJO1366^106^ and iML1515^107^ that are of increasing complexity. We then compared estimated metabolite values and estimated release rates using data from *E. coli* in bioreactors with glucose, galactose, L-malate or L-alanine as the sole carbon source, the same data used in Fig. 2A (Fig. S11). In the L- malate condition for the iJR904 model, we discarded the metabolite value for formate as an outlier because it was extremely small (<10^-12^), below the solver tolerance. Note that no metabolite values were negative and hence no other values were discarded upon log- transformation.

### Metabolite classification, chemical properties and intracellular concentrations

The charge and molecular weight of metabolites were obtained from eciJO1366^42^. All other chemical properties and InChIKeys were obtained from PubChem (RRID:SCR_004284) using the python interface PubChemPy (<u>github.com/mcs07/PubChemPy</u>). The InChIKeys were then used to classify metabolites into a chemical taxonomy using ClassyFire^108^. To get an appropriate level of detail of metabolic classification, we used the “Class” level category. However, we separated the class “Carboxylic acids and derivatives” into “Amino acids” and “Carboxylic acids” based on the “Subclass” level. We also merged the classes “Hydroxy acids and derivatives” and “Keto acids and derivatives” into the joint class “Keto / hydroxy acids”. Finally, metabolites that did not fall into any of the abovementioned categories nor the class “Organooxygen compounds” were joined into the category “Other”.

### Linear models for evaluating the importance of different factors for predicting metabolite release rates

Linear models were created and analyzed using statsmodels (RRID:SCR_016074)^109^. Compound class and carbon source were incorporated as categorical variables. Metabolites predicted to have zero turnover were given a value of 10^-4^, less than the minimum predicted turnover. Release rates with missing values in any of the compared factors were excluded prior to model fitting. Because several metabolites were measured on multiple carbon sources, P values were calculated using standard errors clustered by metabolite. For continuous factors, we tested the slope of univariate linear models. For categorical factors (compound class and carbon source), we used a cluster- robust Wald test of the overall group effect, followed by pairwise cluster-robust Wald tests with Benjamini–Hochberg correction. The same approach was used to compare metabolite values between compound classes (Fig. 2C) and to assess correlations in Fig. S14. To test whether the effect of intracellular concentration on release rate depended on membrane permeability, we fitted linear models including an interaction term between log-scaled intracellular concentration and each permeability-related descriptor separately (charge, molecular weight, log *P*, rotatable bond count, hydrogen bond donor count, hydrogen bond acceptor count and topological polar surface area). P values for the interaction terms were calculated using standard errors clustered by metabolite and corrected for multiple testing across descriptors using the Benjamini– Hochberg procedure. To evaluate the statistical significance of the out-of-sample predictions, we performed 10^4^ permutations of model fitting and out-of-sample estimations where the labels (either metabolite or condition) were shuffled on each permutation. We then calculated the fraction of out-of-sample *R*^2^ values from these permutations below the observed *R*^2^ value to estimate the *P* value.

### Cultivation and exometabolome analyses of KEIO knockout strains

One objective was to study how disruption of key metabolic fluxes would affect extracellular metabolite concentrations and how this would change if these strains were allowed to evolve in a simple nutrient environment. The KEIO collection is a collection of non-essential *E. coli* KO strains that we could use for this purpose^60^. To identify relevant KEIO KO strains, we used eciJO1366 and parsimonious FBA^86^ to predict the effect of a gene knockout on optimal metabolic fluxes, and previous metabolomics data^59^ of these strains to evaluate the effect on intracellular concentrations (Fig. S20). We eventually chose six strains (Δ*aceE*, Δ*cyoD*, Δ*nuoA*, Δ*pgi*, Δ*rpe*, Δ*sucB*) that were predicted to have different optimal flux distributions, had many significantly changed intracellular metabolite concentrations, and targeted different parts of key metabolic pathways (Fig. 4A). We included Δ*lacA* as a negative control expected to behave like the WT as done previously^110^.

The selected strains from the KEIO collection were precultured on LB agar and liquid LB as previously described, and glycerol stocks used for experiments described below were prepared by adding glycerol to a final concentration of 25%.

For exometabolome sampling, the KEIO strains and *E. coli* BW25113 were precultured as detailed above. They were then re-inoculated to OD_600_ = 0.05 in 10 mL M9 + 40 mM galactose (+ 25 µg/mL kanamycin for KEIO strains) in 50 mL Erlenmeyer flasks (200 rpm, 37 °C, 31 h). 1 mL culture was sampled into Eppendorf tubes and washed twice with M9 without carbon source by centrifugation (6000 rcf, 6 min, 4 °C). Ultimately, samples were resuspended in 0.5 mL M9 without carbon source and used to inoculate 200 μL M9 + 40 mM galactose (+25 µg/mL kanamycin for KEIO strains) in a flat bottom 96-well plate to OD_600_ = 0.05, and 6 mL M9 + 40 mM galactose (+25 µg/mL kanamycin for KEIO strains) in straight glass tubes (15.8 cm tall, 1.5 cm diameter) to OD_600_ = 0.005. The 96-well plate was incubated in a Synergy H1 plate reader with continuous shaking (double orbital, 425 cpm, 37 °C), and the glass tubes in a shaking incubator (200 rpm, 37 °C, 45° angle), both for 108 hours. The experiment in the well plate was used to monitor more closely the growth (as OD_600_) of these strains and to help with the timing of the exometabolome sampling. All strains were cultivated in three replicates. Two samples were collected during mid- exponential growth phase for all cultures, mostly to ensure that at least one sample from each replicate was obtained before the end of exponential phase. 1 mL culture volume was centrifuged (14000 rpm, 10 min, 4 °C), and supernatant stored at -70 °C for exometabolome analysis. The second sample for each replicate (Fig. 4B) was analyzed using LC-MS at the UNIL Metabolomics facility, following their protocol described above. Additionally, we analyzed three samples of pooled inocula to verify that any carry-over with the inoculum was much lower (or below detection limit) than the sample metabolite concentrations. Three replicate pooled samples were obtained by inoculating 6 mL M9 + 20 mM galactose medium with all the strains, each strain to an OD_600_ = 0.005, and subsequently handled as described above for the culture samples. Twenty- six (of 111) metabolites not mapping to ECMDB^53^ were filtered out prior to downstream analyses.

### Estimating distance between deleted reactions and measured metabolites

To estimate the distance, in terms of the number of reactions, between the metabolic reaction deleted in KEIO KO strains (Fig. 4A) and the quantified metabolites (Figs. S21-S23), we used the *E. coli* GEM iML1515 to create a bipartite metabolite-reaction network. The bipartite network was constrained to intracellular reactions and metabolites, and we excluded currency-metabolites and cofactors such as NAD, ATP, H_2_O, H^+^ etc. Therefore, we excluded the Δ*nuoA* and Δ*cyoD* strains from this analysis as they are concerned with electron transport using such metabolites and cofactors. The Δ*lacA* strain was also excluded from this analysis because the *lacA* gene is not included in iML1515. The shortest distance between each measured metabolite and the deleted reaction(s) was computed using networkX (RRID:SCR_016864)^111^.

### Chemostat evolution experiment with Δ*aceE* and Δ*sucB* in bioreactors

Two of the KEIO KO strains, Δ*aceE* and Δ*sucB*, were selected for the evolution experiment with continuous cultivation (chemostats) in bioreactors based on their reduced growth compared to their ancestor and difference in exometabolome patterns (Figs. 4A and S21-S23). The strains were streaked from glycerol vials onto LB + 25 µg/mL kanamycin agar plates and incubated at 37 °C overnight. 5-6 colonies were transferred to 500 mL baffled shake flasks containing 75 mL LB + 25 µg/mL kanamycin and incubated for 22 h (200 rpm, 37 °C).

Chemostat cultivations were performed in 1 L DASGIP bioreactors with a constant volume of 300 mL M9 + 20 mM galactose medium + 25 µg/mL kanamycin (prepared as previously described, but without pH adjustment because of practical challenges with the large volumes required, so pH ≈ 7.1). Cultivation was performed at 37 °C with a submerged airflow maintained at 0.5 vvm, and a cascaded agitation of 400-700 rpm, controlled by a DO set point of 30%. pH control was started at day 6 (Δ*aceE*) or day 20 (Δ*sucB*) with a set point of 7.1, controlled using 2 M NaOH (Fig. S25A).

The reactors (three replicates per strain) were inoculated to OD_600_ = 0.1. During the late stage of exponential growth, pumps were started to feed fresh culture medium and remove used medium. Medium was fed using a pre-calibrated peristaltic pump and used medium was removed by a steel pipe placed right above the culture surface. The steel pipe was coupled to a peristaltic pump operating at a flow rate approximately 1.5 times the medium inflow rate, to avoid volume accumulation. The set dilution rate (0–0.1 h⁻¹ for Δ*aceE* and 0–0.3 h⁻¹ for Δ*sucB*) was adjusted several times during the first half of the experiment in response to changes in growth dynamics and in an attempt to increase the number of generations, but was kept constant for long periods, including the full second half of the experiment (Fig. S25B). The actual dilution rate was monitored by keeping the flasks with fresh culture medium on scales. OD_600_ was measured daily, and larger samples were collected weekly. Culture was centrifuged (3220 rcf, 10 min, 4 °C), and supernatant and pellet stored at -70 °C. In addition, cultures were preserved at -70 °C as glycerol stocks. The number of generations was estimated from the measured dilution rates across the experiment plus the generations during the initial batch culture phase before the culture dilution was started.

### Cultivation of single colonies from evolution experiment and sampling of exometabolome

Culture samples from different bioreactors and timepoints collected in the chemostat experiment were streaked out onto LB + 25 µg/mL kanamycin agar plates and incubated overnight at 37 °C. Three colonies were picked and individually inoculated in 20 mL LB + 25 µg/mL kanamycin in 100 mL Erlenmeyer flasks and incubated (37 °C, 200 rpm, 16 h). 12 mL of the culture was then centrifuged (3220 rcf, 6 min, RT), and the pellets stored at -20 °C for sequencing. The isolates were stored as glycerol stocks at -70 °C made from these cultures.

2 mL samples from the precultures were washed as previously described, resuspended in M9 without carbon source, and used to inoculate a flat bottom 96-well plate to OD_600_ = 0.05 with 200 µL M9 + 25 mM galactose + 25 µg/mL kanamycin. The M9 medium was prepared as previously described, but to a pH of 7.1 as in the chemostats. The well plate was incubated in a Synergy H1 plate reader with continuous shaking (double orbital, 425 cpm, 37 °C, 120 h) and the OD_600_ was read every 20 minutes.

Based on these growth measurements we selected the fastest growing isolates from the last timepoint of each of the six bioreactors (three bioreactors per strain). These isolates, the initial KEIO KO strains Δ*aceE* and Δ*sucB*, and the KEIO ancestor *E. coli* BW25113 were precultured and subsequently washed as described above. The washed precultures were used to inoculate 10 mL M9 + 20 mM galactose to OD_600_ = 0.01 in straight glass tubes (15.8 cm tall, 1.5 cm diameter) in triplicates and incubated for 20-84 hours (200 rpm, 37 °C, 45° angle). 1 mL samples for exometabolome analysis were collected at OD_600_ ≈ 1 (Fig. 4, F and G), filtered using a 0.22 μm Millex-GV PVDF syringe filter (SLGVR33RS, MilliporeSigma, Darmstadt, Germany) into a 1.5 mL Eppendorf tube stacked in a cold block (-20 °C), and then stored at -70 °C until it was analyzed. The exometabolome LC-MS analyses were conducted by the UNIL Metabolomics facility, following the procedure described above.

### DNA extraction of samples from selected isolates and chemostat culture samples

The DNA extraction of 14 different samples from the chemostat cultures (#37A D44 M2 Pellet, #38A D44 M3 Pellet, #39A D44 M4 Pellet, #28B D30 M5 Pellet, #29B D30 M6 Pellet, #30B D30 M7 Pellet) and the selected isolates (sucB-M5-D30-4, sucB-M6-D30-6, sucB-M7-D30-4, sucB- Ancestor, aceE-M2-D44-2, aceE-M3-D44-3, aceE-M4-D44-1, aceE-Ancestor) was performed by adapting the protocol from the FastPure Bacteria DNA Isolation Mini Kit (DC103-01, Vazyme, Nanjing, China). 1 mL GA buffer was added to the thawed pellet samples, and 230 µL of these resuspensions were kept in Eppendorf tubes for DNA extraction. The rest was centrifuged (4000 rpm, 6 min, RT) and the pellet stored at -20 °C. Genomic DNA (gDNA) was extracted from the 230 µL sample following the manufacturer’s protocol. In the final step, elution of the DNA was performed twice with fresh elution buffer, eluting a total of 100 µL DNA. Extracted gDNA was stored at -70 °C until sequencing.

### Genomic sequencing processing

Genomic DNA from the selected isolates and chemostat samples was sequenced by Novogene (Planegg, Germany) using the Illumina NovaSeqX PE 150. The isolates and the bioreactor samples were sequenced to 1 Gb and 10 Gb sequencing depth, respectively. Good quality of both datasets was ensured using FastQC v. 0.12.1 (RRID:SCR_014583) before trimming the reads with fastp v. 0.24.0 (RRID:SCR_016962; parameters --qualified_quality_phred 20 --cut_front --cut_tail -- average_qual 20 --length_required 50)^112,113^. Both read sets were mapped against the *E. coli* BW25113 reference genome (NCBI RefSeq assembly GCF_000750555.1) with minimap2’s v. 2.28 (RRID:SCR_018550) default “sr” parameters^114^. The resulting alignments were filtered using SAMtools view (v. 1.20; RRID:SCR_002105; parameters -b -f 3 -q 60) to retain only properly paired reads mapped to the reference with a mapping quality score (MAPQ) ≥ 60^115^. Assessment of sequence coverage revealed a minimum average depth of 2017X for the chemostat culture samples and 190X for the sequenced isolates. For both datasets, variants were identified with FreeBayes v. 1.3.6 (RRID:SCR_010761; parameters --min-alternate-count 3 -p 1 --min-alternate- fraction 0.05 --pooled-continuous --haplotype-length 0) and annotated with SnpEff v. 5.0 (RRID:SCR_005191) using the BW25113 reference genome^116,117^. A custom python script was used to filter the annotated variants. For the isolates, variants with a frequency lower than 0.95 and a coverage lower than 100 were filtered out. For the chemostat culture samples, variants with a frequency lower than 0.05 and coverage lower than 100 were filtered out. A variant was considered fixed if its allele frequency was at least 0.9.

### Paired intracellular and extracellular metabolomics and live/dead staining

M9 medium with 20 mM galactose, 30 mM malate or 40 mM L-alanine was prepared as detailed above. *E. coli* K-12 MG1655 was precultured on LB agar plates and in liquid LB as detailed above. 1 mL preculture was washed as described above and used to start a second preculture in 20 mL M9 + 40 mM L-alanine, inoculated to OD_600_ = 0.05. Another LB preculture of *E. coli* K-12 MG1655 was used to inoculate a second preculture in M9 + 20 mM galactose and one in M9 + 30 mM malate in the same way 12 hours later. After 36/24 hours, 1 mL was collected from each of the three precultures and washed twice as detailed above. These washed precultures were used to inoculate 10 mL of the same media (M9 + either 20 mM galactose, 30 mM malate or 40 mM L- alanine) in straight glass tubes (15.8 cm tall, 1.5 cm diameter). These glass tubes were incubated for two days (200 rpm, 37 °C, 45° angle). Growth was monitored by measuring OD_600_ directly in the glass tubes (NANOCOLOR VIS II, MACHEREY-NAGEL, Düren, Germany) at 21 timepoints. The glass tubes were vortexed briefly before any OD reading or sampling.

A reference exometabolome 0.5 mL sample was collected after 2 hours following the procedure detailed below. Then, 2 mL samples for paired intra- and extracellular metabolome and live/dead analyses were collected in the late exponential phase (at OD_600_ ≈ 1.1, Fig. S34). Exometabolome samples were obtained by filtering 0.5 mL culture using a 0.22 μm Millex-GV PVDF syringe filter (SLGVR33RS, MilliporeSigma) into a 1.5 mL Eppendorf tube stacked in a cold block (-20 °C). The samples for intracellular metabolomics, i.e. cells, were obtained immediately after following this procedure: A 25 mm 0.22 μm Durapore PVDF filter (GVWP02500, MilliporeSigma) was placed in a new filter holder. A 5 mL syringe was connected to the inlet of the filter holder, and a vacuum pump (250 mbar) was connected to the outlet of the filter holder. Then, 2 mL of 37 °C MQ water was added to pre-wet the filter. Immediately after, 1 mL of culture sample was added to the syringe and allowed to pass through the filter. If the culture did not pass through solely based on the vacuum pump, light additional pressure was added by using the syringe itself to push the culture through the filter. This process took 10-30 seconds. Still using the same set-up, the filter was finally rinsed with 5 mL of 37 °C MQ water as previously recommended^54^. Two tweezers were quickly rinsed in MQ water and used to carefully transfer the filter from the filter holder to 2 mL lysis tubes filled with 1.4 mL of 80:20 methanol:water (v/v), pre-chilled to -20 °C and kept on a cold block also pre-chilled to -20 °C. All samples were then quickly transferred to -70 °C for storage. Live/dead staining was performed using a similar approach as previous work^118^: Briefly, the *live* sample was made by diluting 10 μL culture in 990 μL PBS and the *dead* sample was made by mixing 100 μL culture with 1 mL 70% isopropanol. Both samples were vortexed and incubated at room temperature for 1 h. Then, the isopropanol was removed from the dead sample by centrifugation (8000 rcf, 4 min, RT), pipetting off the supernatant, resuspending the pellet in 1 mL PBS and vortexing. This procedure was repeated once. Then, both the live and the dead sample were stained with propidium iodide (PI; P4170, Sigma-Aldrich) and SYBR green (Invitrogen S7563, Thermo Fisher Scientific, Carlsbad, CA, USA) with the following procedure: 48 μL sample (live or dead) was mixed in a 96-well plate with 49 μL PBS, 2 μL 0.5 mg/mL PI and 1 μL 100X SYBR green and incubated for 15 min in the dark. Then, 0.1X samples of both the stained live and the stained dead samples were created by diluting 10 μL of each sample in 90 μL of PBS. The stained live/dead samples were then analyzed immediately on a CytoFLEX S flow cytometer (Beckman Coulter, Brea, CA, USA) and the CytExpert software (v2.4.0.28, RRID:SCR_017217). The dead samples were used to make the gates required to quantify the fraction of dead cells in the live samples on the PE-H and FITC-H signals (Fig. S34, Table S7). The extra- and intracellular metabolomics samples were analyzed by the metabolomics facility at UNIL to quantify the five metabolites leucine, glutamate, aspartate, fructose 6-phosphate and *cis*-aconitate. These metabolites were chosen because they were chemically and metabolically diverse and likely to be quantifiable both intracellularly and extracellularly based on previous experience with similar samples. Also, as glutamate is the most abundant intracellular metabolite^54,119^, it serves as a best-case scenario in our attempt to test how much of the extracellular metabolite levels can be explained by cell lysis and the associated release of intracellular metabolites. Intracellular metabolite samples were first normalized by the amount of protein detected in the same sample and subsequently converted to absolute intracellular concentrations using the estimated protein density in *E. coli* (13.5 × 10^-8^ µg/µm³)^120^. The fructose 6-phosphate value for sample 1B, timepoint 1, was discarded as an outlier as it was one order of magnitude larger than any other fructose 6-phosphate measurements.

### Contribution from intracellular metabolites and cell lysis to extracellular concentrations

Approximate cell volume was estimated from measured OD_600_ and a previous estimate of OD to cell volume for *E. coli* of 3.6 μL/OD_600_/mL^121^. From approximate cell volume, measured absolute intracellular concentrations and cell lysis fractions we then calculated the corresponding change in extracellular concentrations. This potential contribution was then compared to the net change in extracellular concentrations between the T0 reference sample (after 2 hours) and the late exponential phase sample. If all T0 values were below the detection limit, we assumed the initial concentration of this metabolite to be 0. If only some values at T0 were below the detection limit we imputed these values to the mean of the detected values, giving conservative estimates for the net change in extracellular concentrations.

To identify typical biomass degradation products, we used the metabolites consumed by the biomass equation of the *E. coli* model iML1515^107^, although excluding the soluble pool (as categorized in^105^) since this should correspond to the intracellular concentrations already accounted for. Of the metabolites measured in our dataset (Fig. 3B), only the amino acids were present among the biomass components; therefore, protein depolymerization accounts for the entire contribution from biomass degradation in this analysis. To estimate a potential contribution from protein depolymerization we estimated cell density in gDW/L from OD_600_ (0.346 gDW/L/OD_600_) and from this specific amino acid density (in g/L) from recent estimates of *E. coli* amino acid mass fractions^122^. When mass fractions were attributed to pairs of amino acids (glutamate/glutamine and aspartate/asparagine), we assumed equal contributions. Amino acid density was converted from g/L to μM by dividing by the amino acids’ molecular weight minus the weight of a water molecule to account for the condensation reaction in protein polymerization. A similar procedure as described above was followed when we extended this analysis by using literature values for intracellular concentrations. To map extracellular concentrations in our dataset to literature values for intracellular concentrations, we used the following procedure: We first tried to map to literature values from cultivations in minimal medium with the same growth phase (exponential/stationary) and carbon source. If this was not possible, we prioritized mapping to data from minimal medium conditions of the correct growth phase. If concentrations from minimal medium conditions were not available, we mapped to data from complex growth media. When multiple intracellular concentrations mapped to our extracellular data point, we used the median of the intracellular concentrations. When extracellular concentrations were reported as two indistinguishable molecules, we attributed equal amounts to the two metabolites. This includes 2-phosphoglycerate/3-phosphoglycerate and ribulose 5-phosphate/xylulose 5-phosphate.

### Large-scale screening of metabolite release from transporter KO mutants

We selected 67 strains from the KEIO collection^60^ to assess if transporters (or porins/channels) play a role in metabolite release (Table S5). Each of these strains lacks a gene associated with metabolite transport. Functional annotations of these genes were obtained from EcoCyc (RRID:SCR_002433)^64^ and TCDB (RRID:SCR_004490)^123^. All strains were precultured and washed as described above and then inoculated to OD_600_=0.01 in 10 mL of 20 mM M9 glucose medium (pH=7.4) in straight glass tubes (15.8 cm tall, 1.5 cm diameter) for 20 hours (37 °C and 200 rpm). The glass tubes were angled 45° in the shaking incubator to improve mixing and aeration. The whole experiment was conducted in 5 separate batches. The KEIO reference strain *E. coli* BW25113 was included in 6 technical replicates in each batch. All other strains were cultivated in three technical replicates.

During the experiment, growth was monitored by measuring OD_600_ directly in the glass tubes on 1- or 2-hour intervals (NANOCOLOR VIS II, MACHEREY-NAGEL, Düren, Germany). The exometabolomes were sampled at 4, 8, 10, 11, 12, 13, 14, 15, 16 and 20 hours (+ at 23 and 24 hours for Δ*tolC*). These timepoints were selected to cover the whole exponential and stationary phase, with highest temporal resolution in the last part of the exponential phase and during the transition to stationary phase where we expected the largest changes in extracellular metabolite concentrations to occur. The exometabolome sampling was carried out by first vortexing each glass tube and then immediately collecting 150 μL into a 96-well filter plate (Millipore MultiScreen HTS GV Filter Plate, 0.22 µm Hydrophilic PVDF membrane, MSGVN22). The filter plate was stacked on top of a receiver plate and briefly centrifuged (1 min, 1000 rcf) to filter the cultures. The receiver plate and the filter plates were both immediately placed on an aluminum plate pre- chilled and kept on dry ice to cool down the samples quickly before the samples were moved to a -70 °C freezer.

The samples were measured by flow-injection analysis time-of-flight mass spectrometry (FIA-TOF MS) using an Agilent 6550 quadrupole time of flight mass spectrometer as previously described^62^. Samples were injected online utilizing an Agilent 1100 Series HPLC system (Agilent) coupled to a Gerstel MPS 3 autosampler (Gerstel). Each sample was injected four times. The mobile phase flow rate was set at 0.15 mL/min, with the isocratic phase composed of 60:40 (v/v) isopropanol and water buffered to a pH of 9 with 4 mM ammonium fluoride. The instrument was run in 4 GHz full- scan mode, collecting spectra between 50 and 1000 *m/z*. Online mass axis correction was performed with taurocholic acid and hexakis(1H,1H,3H-tetrafluoropropoxy)phosphazene. For these samples, masses were annotated by exact mass to within 1.0 ppm of the theoretical monoisotopic mass against *E. coli* KEGG metabolites via in-house software as previously described^62^. Processed mzML files for this dataset can be found at MassIVE: MSV000097105. Samples from all batches were initially analyzed in one continuous run (randomized within sampling plates) with four injections (technical replicates) of each sample. A standard was analyzed at regular intervals between samples. However, because of the large number of injections (>10000) instrument performance declined considerably during the analysis of batch 5, which was therefore re-analyzed later in a separate run. In some cases, the complexity of MS data (i.e. close neighboring peaks) yielded improperly centroided mass values which subsequently failed annotation to this stringent and highly conservative mass accuracy. This led to some differences in annotated metabolites between batches 1-4 and batch 5.

We conducted thorough quality control of both cultivation data and exometabolome data prior to downstream analyses. Three strains (Δ*tolC*, Δ*dppB*, and Δ*ptsG*) were discarded because of severely impaired growth, and one (Δ*mscS*) following strain verification checks that identified an ambiguous phenotypic profile. Manual inspection of both raw and total ion current (TIC)- normalized datasets confirmed that TIC normalization minimized technical noise; TIC-normalized peak areas were therefore selected for downstream analyses. First, we filtered the dataset by removing *m/z* features where none of the KEGG annotations mapped to ECMDB^53^. SMILES from ECMDB were then used to classify metabolites with ClassyFire^108^. We used primarily the superclass level to group metabolites, but we separated the class “Organic acids and derivatives” into “Amino acids”, “Carboxylic acids”, “Keto / hydroxy acids” and “Other organic acids”. To reduce complexity, we further merged the classes that mapped to few metabolites into a class named “Other”, i.e. “Benzenoids”, “Other organic acids”, “Phenylpropanoids and polyketides” and “Organic nitrogen compounds”. We then removed severe outliers, focusing on occurrences where multiple consecutive injections were affected, as these errors could propagate despite using the median rather than the mean of 4 injections in further data analyses. We standardized the data from batch 1-4 and batch 5 separately after removing outliers and before calculating the median value across the 4 injections from each sample.

To quantify the effect of each KO on each metabolite, we compared the time-series median standardized Z-score between the KO mutant and the WT, only using the data from the same batch. To calculate the spread of the data for the WT and KO mutant we first used SciPy (RRID:SCR_008058)^124^ to fit a smoothing spline (function *make_smoothing_spline,* with regularization parameter = 10) to all median Z-score values of each strain with the area under the curve of the OD_600_ readings on the x-axis (Fig. 5A), with weights inversely proportional to the standard deviation of each data point. We then calculated the spread as the mean absolute deviation from this smoothing spline. We calculated the difference between the KO strain and the WT as the mean distance between the two smoothing splines across x-axis locations corresponding to the underlying datapoints and within the x-range of both datasets. We then calculated the *Effect* as the ratio between the mean difference and the mean spread. To assess the significance of the estimated effects, we repeated this procedure over 1000 permutations where we shuffled the KO strain / WT label of all data points in each permutation. We then counted the number of permutations where we observed a higher / lower effect than in the actual data (for positive and negative effects).

### Amino acid auxotroph co-cultures

*E. coli* strains auxotrophic for proline (*E. coli* K-12 MG1655 Δ*proC* mCherry), isoleucine (*E. coli* K- 12 MG1655 Δ*ilvE*) and methionine (*E. coli* K-12 MG1655 Δ*metB* GFP) were kindly provided by Estelle Pignon^125,126^. The strains were transformed with the plasmid pEP28 providing chloramphenicol resistance^125^. We then cultivated each auxotroph in monoculture, with *E. coli* BW25113 and with 2-4 relevant transporter KO strains (proline: Δ*proP*, Δ*putP*; methionine: Δ*brnQ,* Δ*cstA,* Δ*metI,* Δ*mtr*; isoleucine: Δ*glnP*, Δ*tsx*, Δ*yeaS*, Δ*ygaZ*). Strains were precultured (with 25 μg/mL chloramphenicol / kanamycin for auxotrophs / KEIO KO strains) and washed as previously described, inoculated to OD_600_ = 0.001 in 20 mM glucose M9 medium (200 μL) and cultured in BioTek Synergy H1 platereader (24 hours, 37 °C) as previously described. After 24 hours, the auxotrophs’ abundances were quantified as CFUs/mL on LB+25 μg/mL chloramphenicol agar plates. Because methionine was not quantified in the batch 5 data, we had no information about the effect of Δ*metI* on methionine from the main dataset. However, not all metabolites were affected by the deteriorating mass-spec quality in the first run of batch 5. We therefore went back to the first run of batch 5, verified that the distribution of methionine peak intensities was similar in batch 5 as in batch 1-4, and then estimated the effect of Δ*metI* on methionine from these data (*Effect* = 2.56, *P < 0.005*).

