## Supplementary Figures for "Microbes use transporters to regulate the release of metabolites based on value"

Figures S1-S34

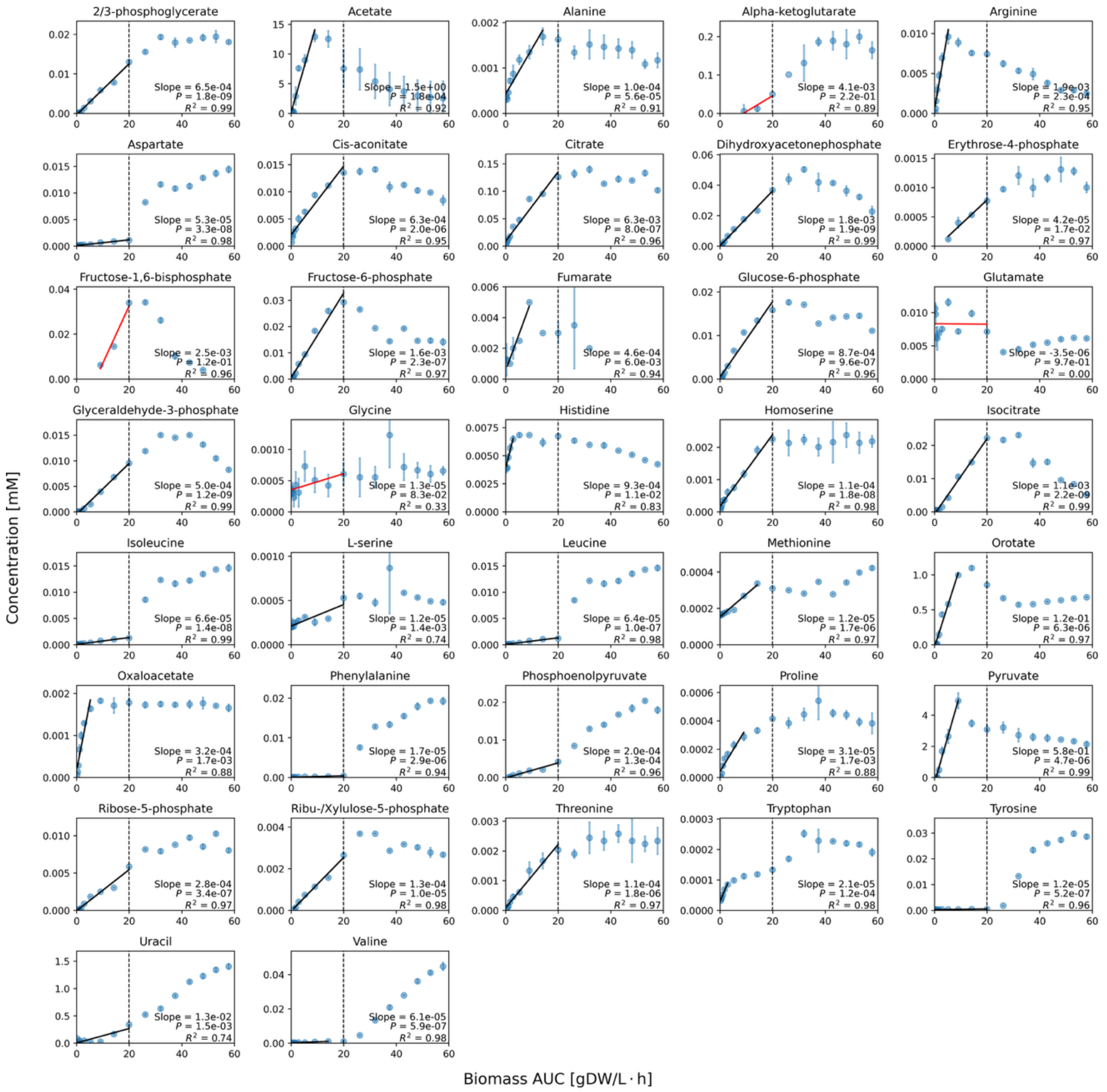


*Figure S1: Estimated rates of metabolite uptake or release in E. coli derived from extracellular metabolite concentrations[1]. Rates were calculated from metabolites with at least three data points within the exponential growth phase. To account for saturation effects or reconsumption, data points beyond the point of saturation in extracellular concentration were excluded from the rate estimation. The initial dynamics of 89% (33/37) of metabolites are well explained by a linear model ($R^2$ > 0.5 and $P$ < 0.05). Others are highlighted in red. The vertical dashed line marks the transition from exponential to stationary phase.*

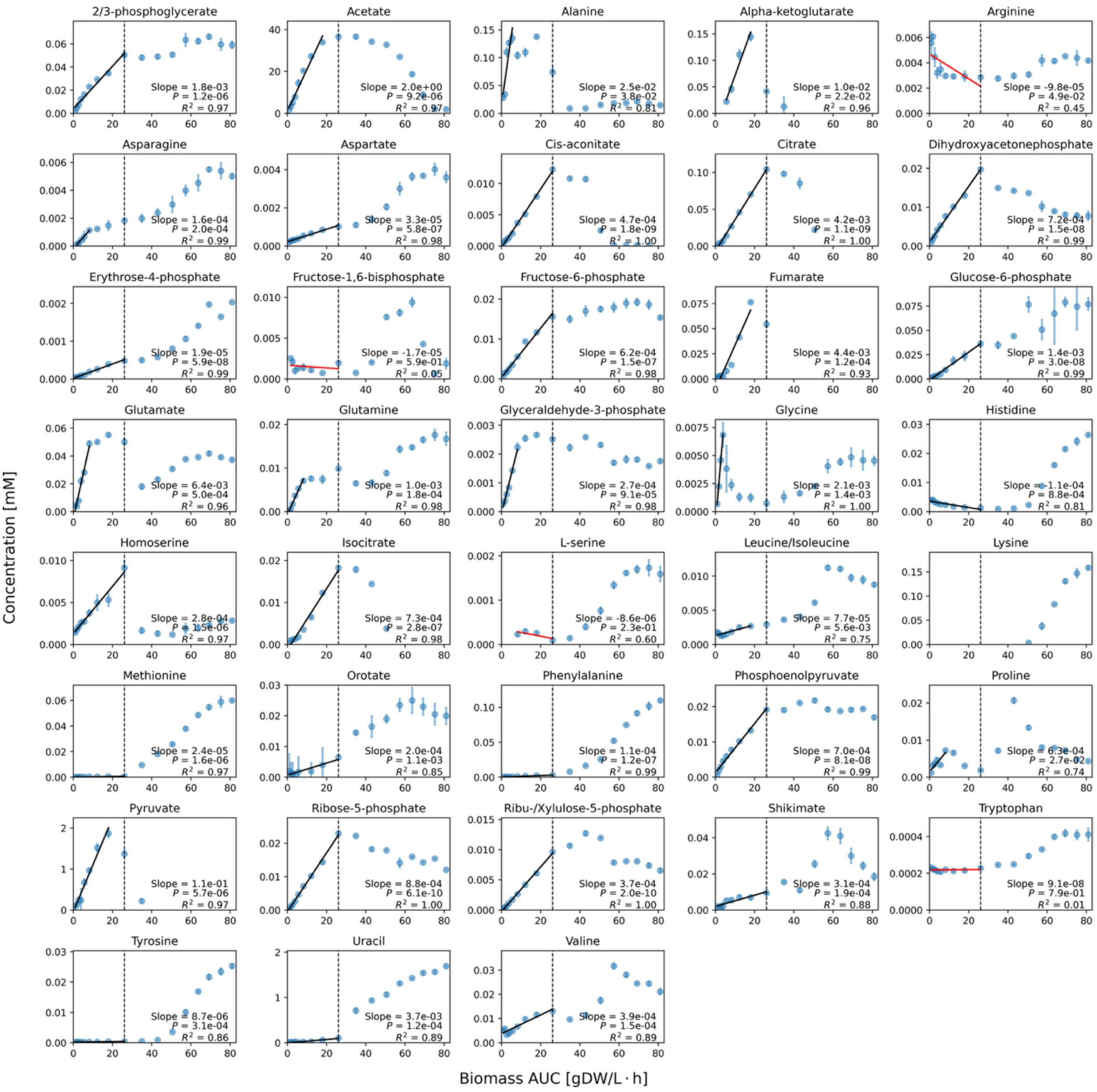


*Figure S2: Estimated rates of metabolite uptake or release in B. licheniformis derived from extracellular metabolite concentrations[1]. Rates were calculated from metabolites with at least three data points within the exponential growth phase. To account for saturation effects or reconsumption, data points beyond the point of saturation in extracellular concentration were excluded from the rate estimation. The initial dynamics of 89% (33/37) of metabolites are well explained by a linear model ($R^2 > 0.5$ and $P < 0.05$), not counting metabolites with too few data points to estimate the rate. Others are highlighted in red. The vertical dashed line marks the transition from exponential to stationary phase.*

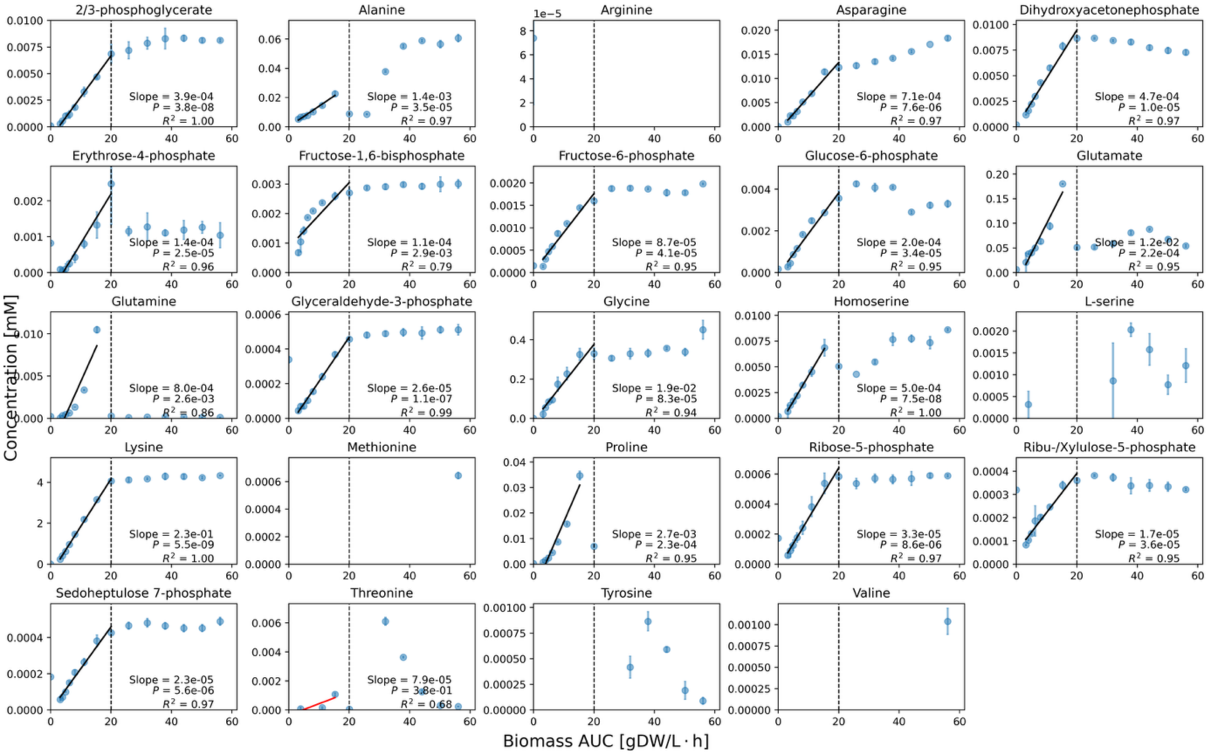


*Figure S3: Estimated rates of metabolite uptake or release in C. glutamicum derived from extracellular metabolite concentrations[1]. Rates were calculated from metabolites with at least three data points within the exponential growth phase. To account for saturation effects or reconsumption, data points beyond the point of saturation in extracellular concentration were excluded from the rate estimation. The initial dynamics of 95% (18/19) of metabolites are well explained by a linear model ($R^2 > 0.5$ and $P < 0.05$), not counting metabolites with too few data points to estimate the rate. Others are highlighted in red. The vertical dashed line marks the transition from exponential to stationary phase. We only show the first 24 hours of this data, and for the full timecourse (ca. 250 hours) we refer to the original publication[1].*

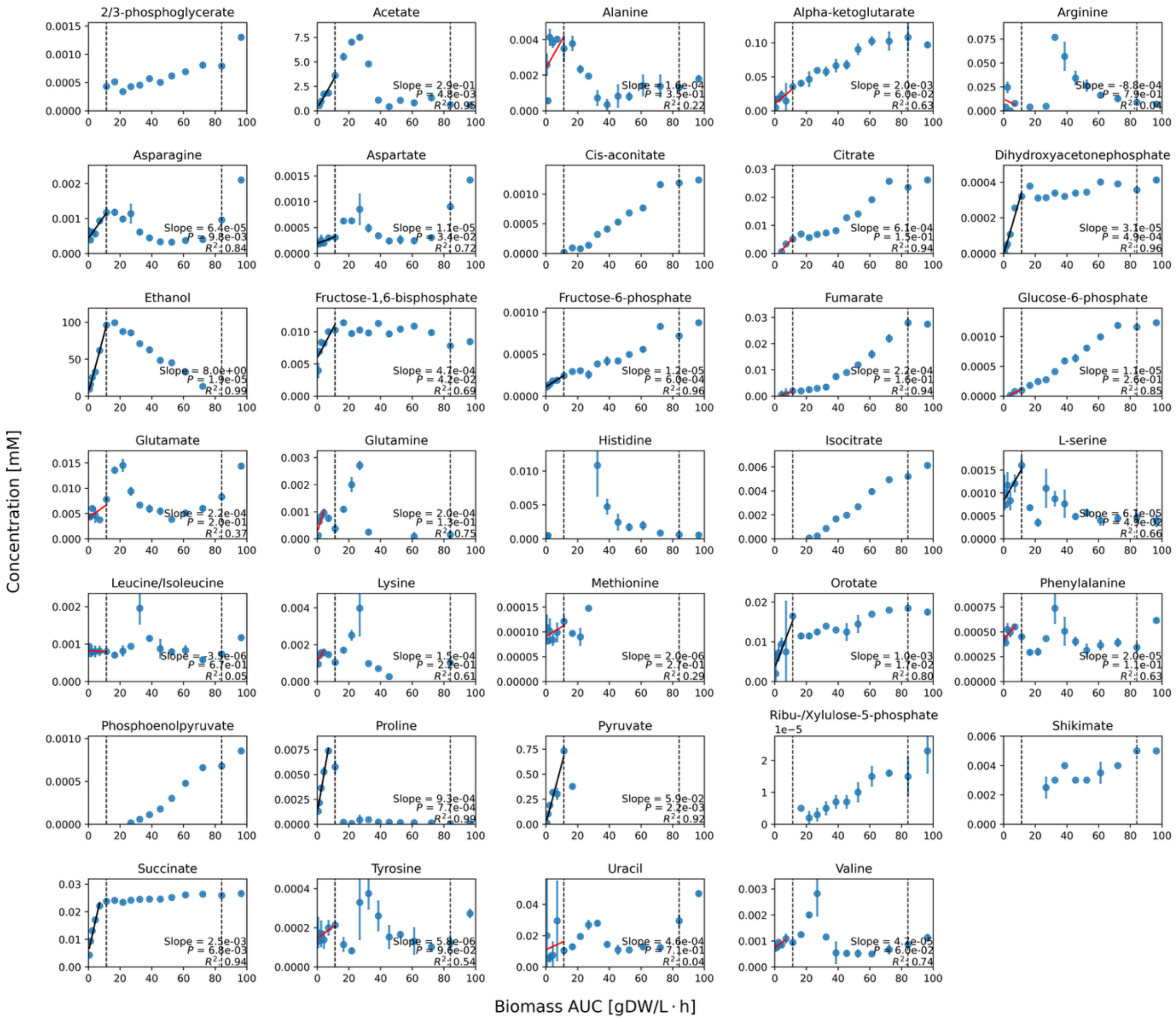


*Figure S4: Estimated rates of metabolite uptake or release in S. cerevisiae derived from extracellular metabolite concentrations[1]. Rates were calculated from metabolites with at least three data points, spanning the first exponential growth phase, before the diauxic shift where acetate and other metabolites are being reconsumed. To account for saturation effects or reconsumption, data points beyond the point of saturation in extracellular concentration were excluded from the rate estimation. The initial dynamics of 44% (12/27) of metabolites are well explained by a linear model ($R^2 > 0.5$ and $P < 0.05$), not counting metabolites with too few data points to estimate the rate. Others are highlighted in red. The first vertical dashed line shows the diauxic shift and the second dashed line marks the transition to stationary phase.*

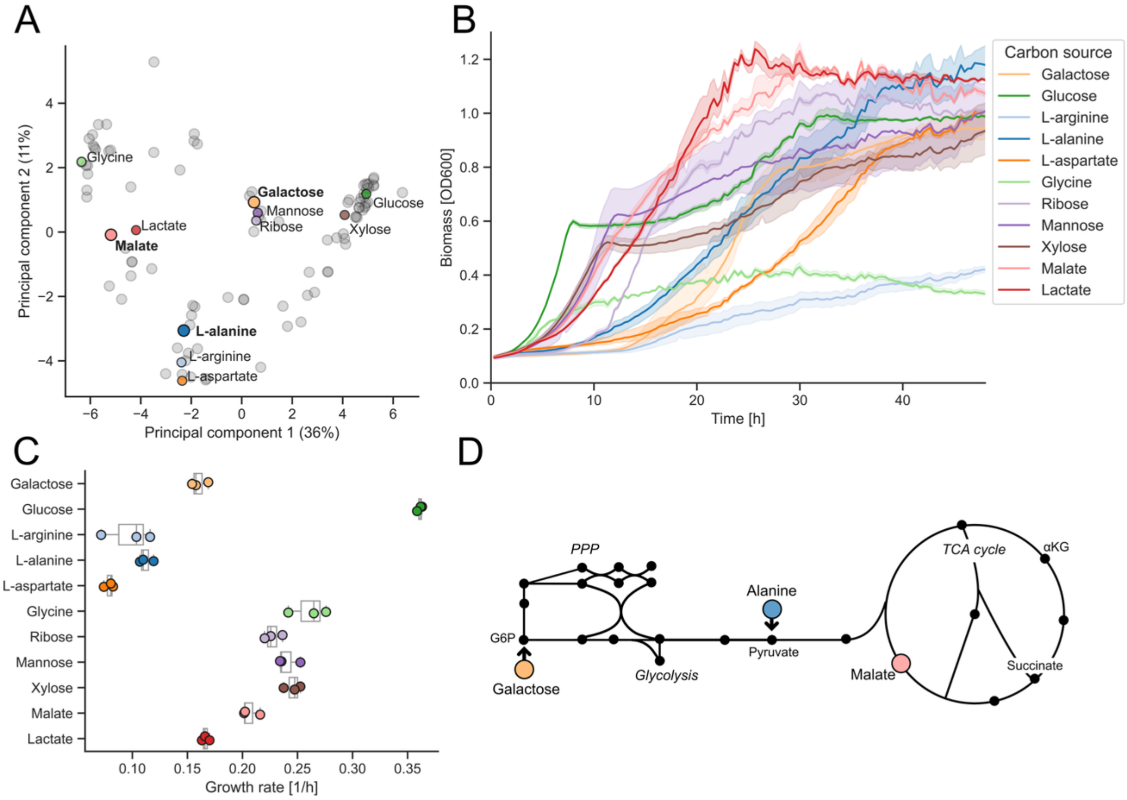


*Figure S5: Selection of carbon source for E. coli batch culture and exometabolome sampling experiments. A) A PCA of flux patterns predicted using pFBA and eciJO1366[2]. B) Growth curves for the screened carbon sources. C) The growth rate of E. coli on the different carbon sources. D) The entry point into metabolism of the selected carbon sources: L-alanine, L-malate and galactose.*

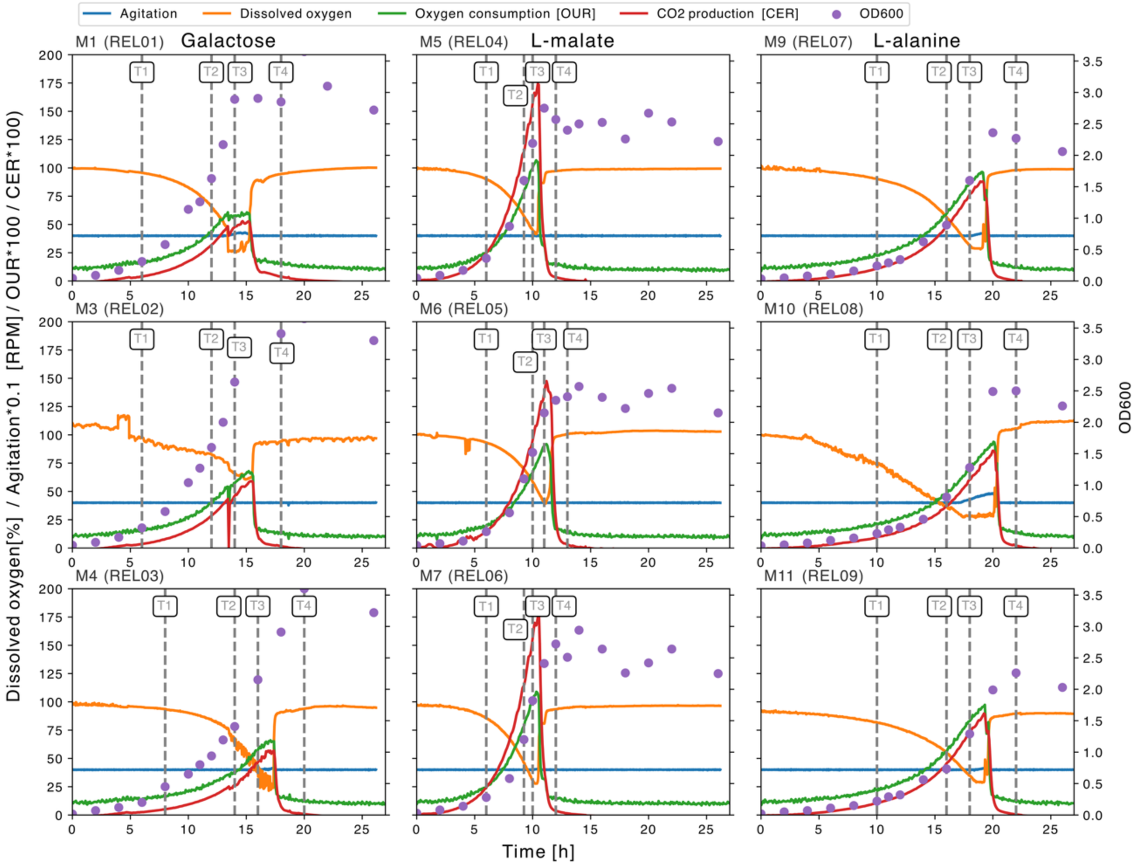


*Figure S6: Bioreactor data and sampling time for E. coli batch cultures in minimal M9 media with galactose, L-malate or L-alanine as the carbon source. Each column shows the three replicates for each carbon source. The timepoints for exometabolome sampling and analyses are annotated with vertical gray lines and labels. These selected exometabolome sampling timepoints are aligned with respect to the end of the exponential phase.*

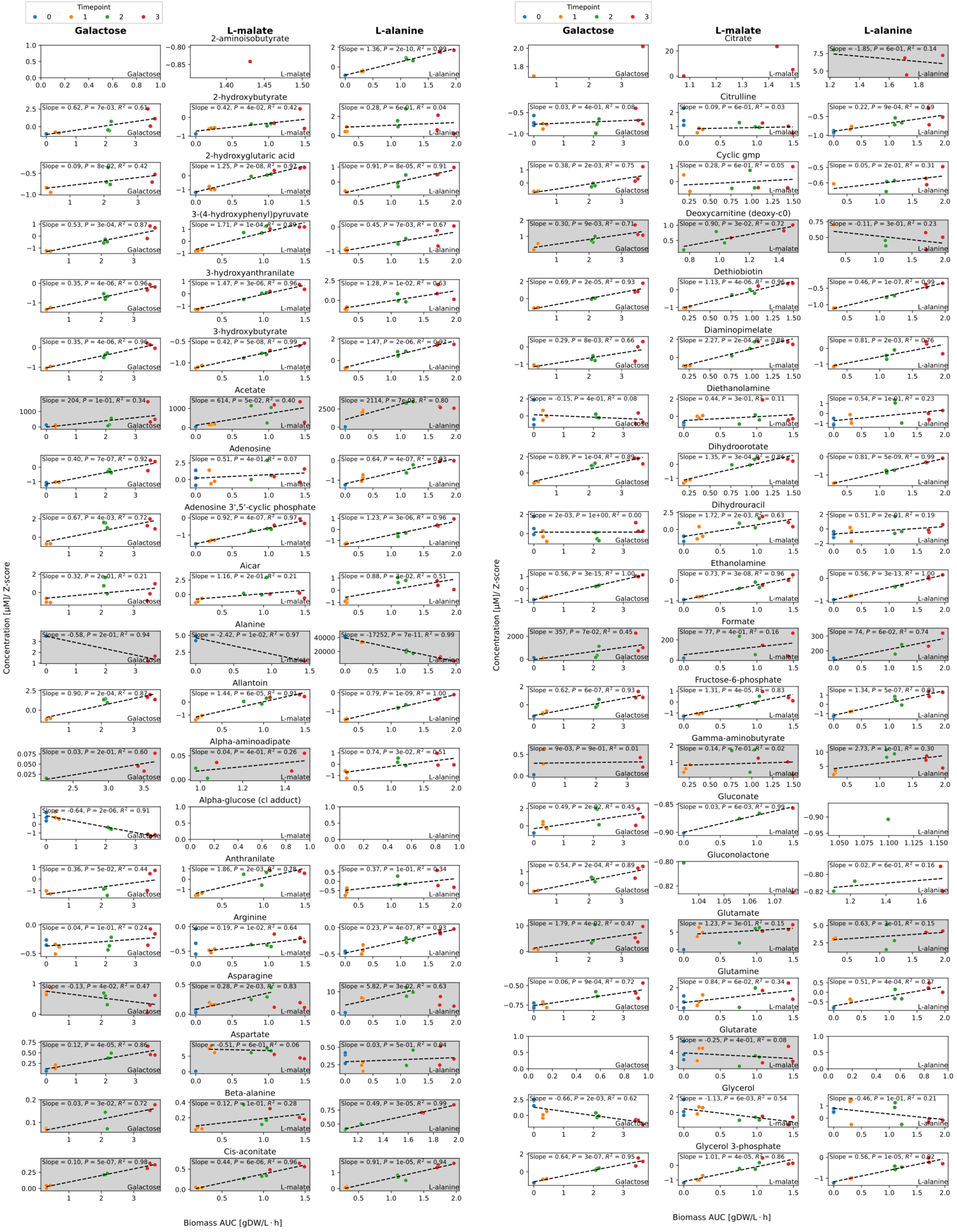


*Figure S7: Specific net uptake and release rates for E. coli cultivated in minimal medium containing galactose, L-malate, or L-alanine as the sole carbon source. Gray shading highlights rates calculated from absolute metabolite concentrations; unshaded panels indicate rates derived from relative (standardized peak area) measurements. This figure displays the first subset of measured metabolites (see Figs. S8 and S9 for the remaining data).*

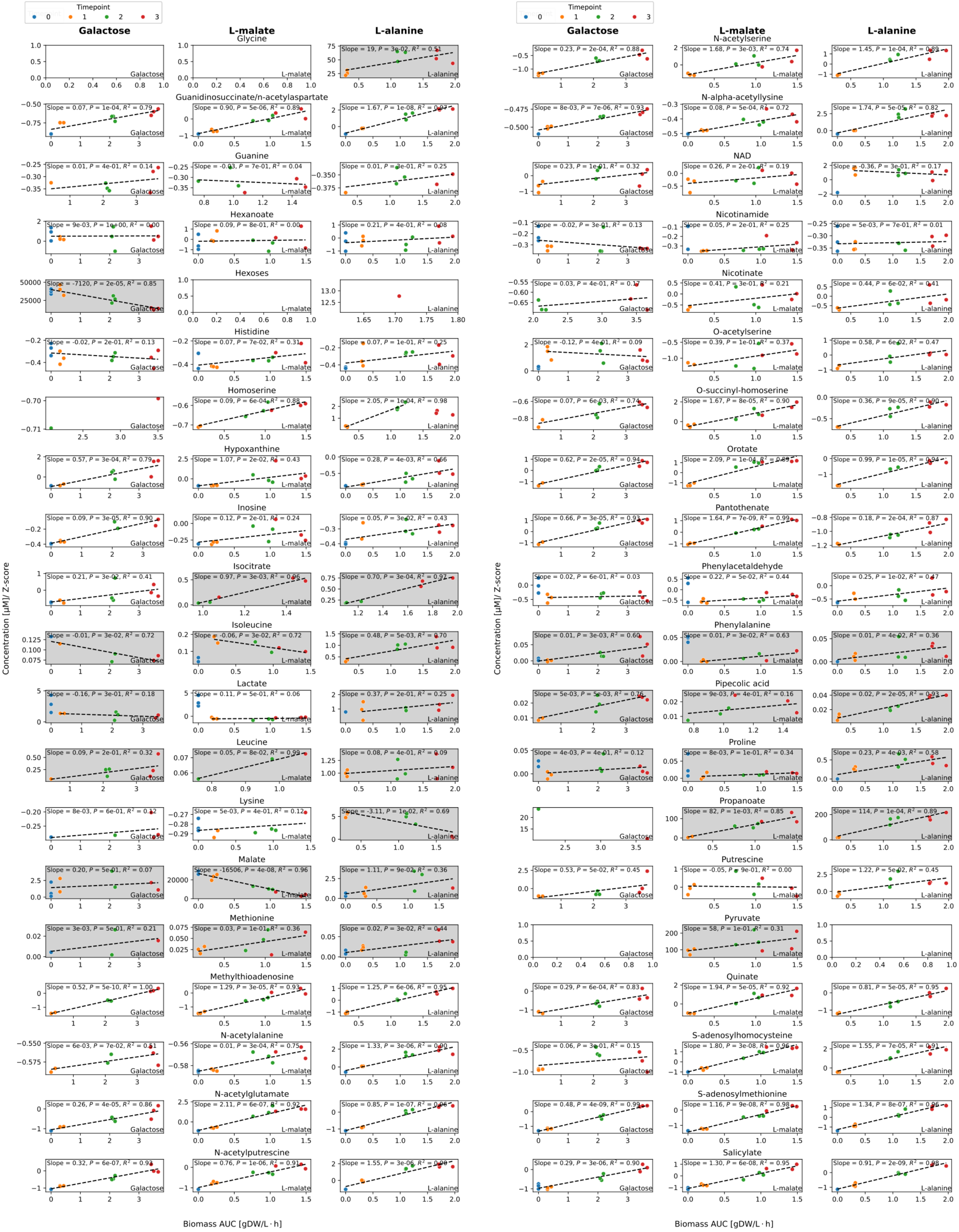


*Figure S8: Specific net uptake and release rates for E. coli cultivated in minimal medium containing galactose, L-malate, or L-alanine as the sole carbon source. Gray shading highlights rates calculated from absolute metabolite concentrations; unshaded panels indicate rates derived from relative (standardized peak area) measurements. This figure displays the second subset of measured metabolites (see Figs. S7 and S9 for the remaining data).*

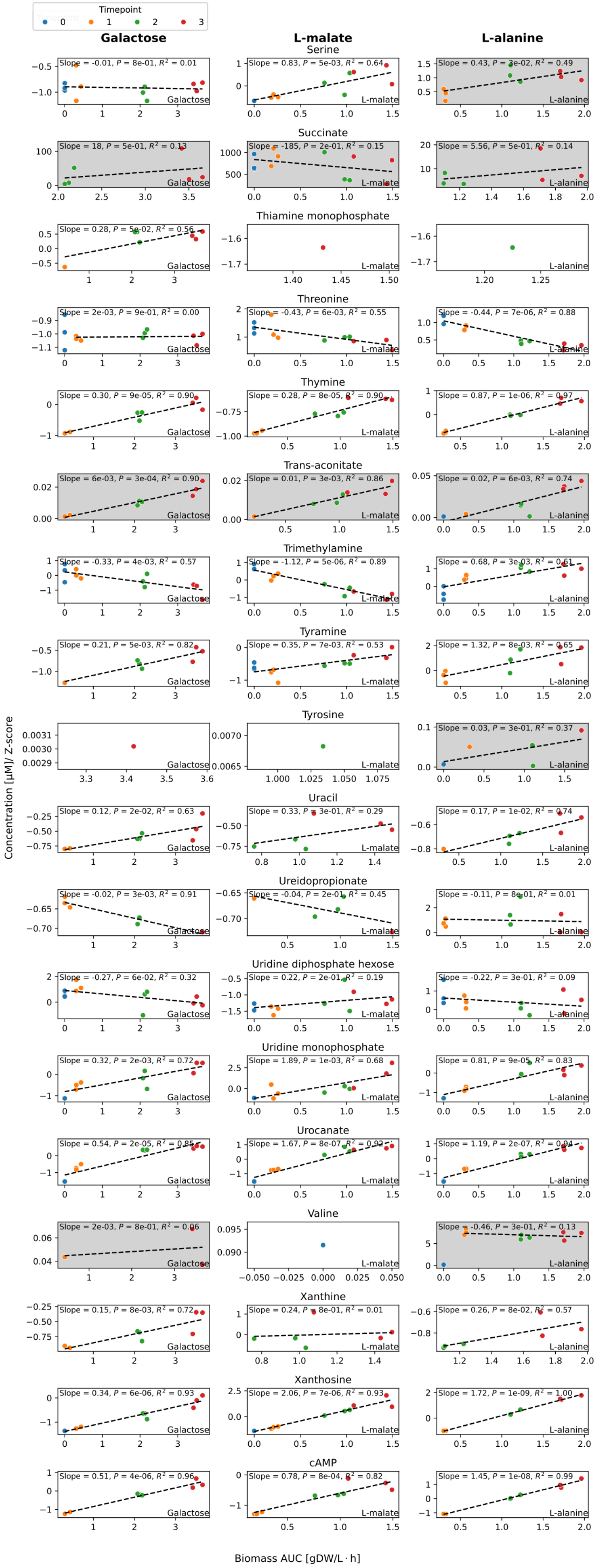


*Figure S9: Specific net uptake and release rates for E. coli cultivated in minimal medium containing galactose, L-malate, or L-alanine as the sole carbon source. Gray shading highlights rates calculated from absolute metabolite concentrations; unshaded panels indicate rates derived from relative (standardized peak area) measurements. This figure displays the third subset of measured metabolites (see Figs. S7 and S8 for the remaining data).*

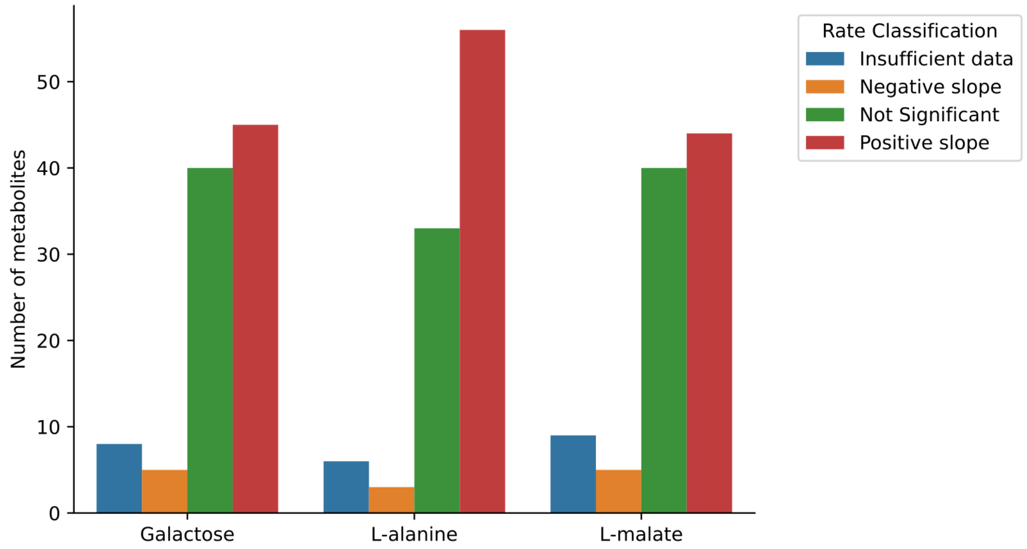


*Figure S10: Classification of estimated rates from E. coli batch cultures with galactose, L-malate or L-alanine as the carbon source (Figs. S7-S9). We find that 45-57% of metabolites have significant positive slopes (two-sided Wald test, FDR<0.05).*

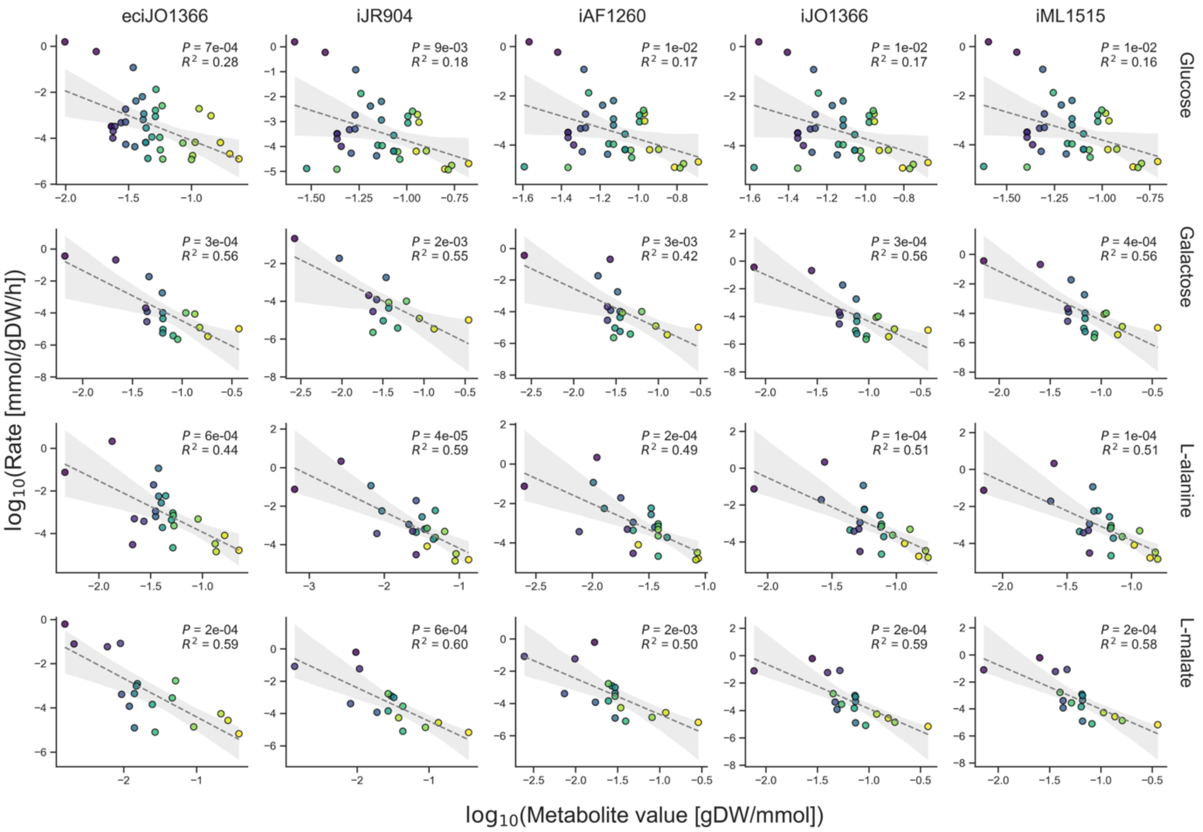


*Figure S11: Comparison of estimated metabolite values from different genome-scale metabolic models (GEM) of E. coli. The leftmost column displays the enzyme-constrained model eciJO1366 used in the main results of this work, and the other columns display different curated E. coli models covering an increasing number of genes (as indicated by the model names). Each row displays the results for each carbon source. The data points on each row are colored by the order of metabolite value as predicted by eciJO1366 in that condition to aid comparison of metabolite values between models.*

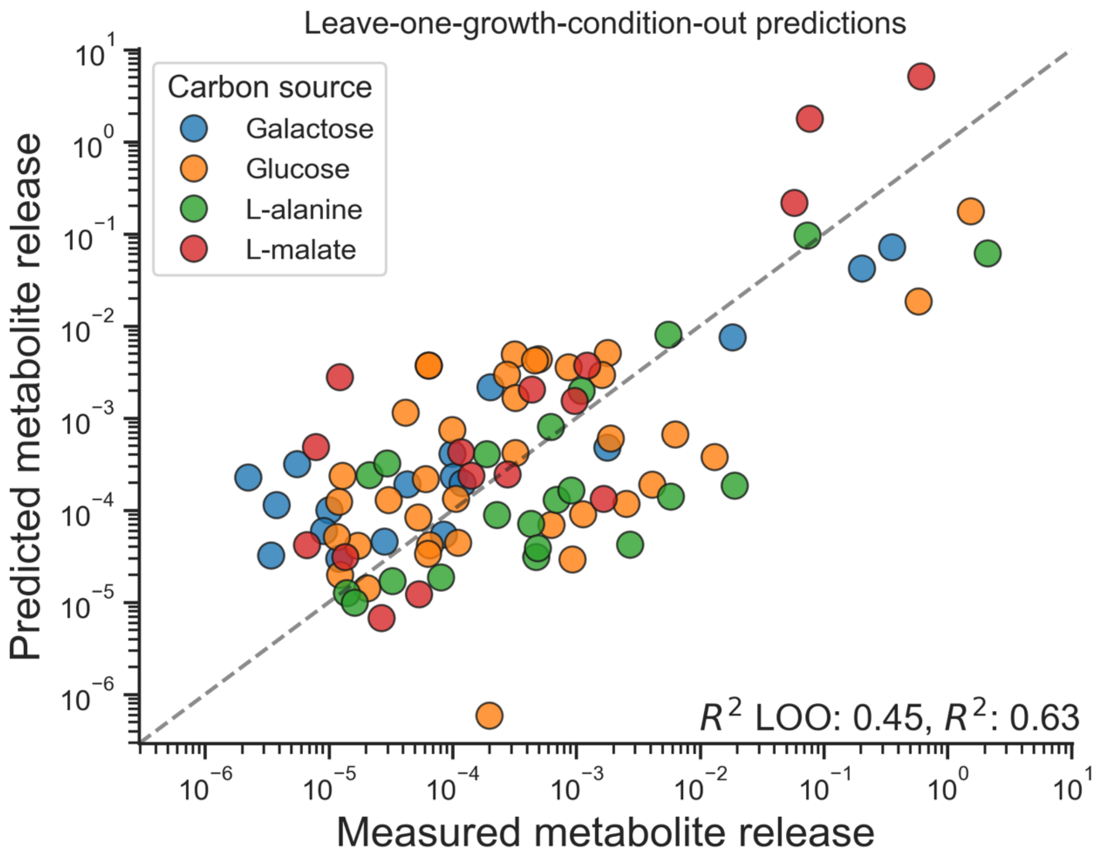


*Figure S12: Cross-validation of best linear model (ranked by BIC) for predicting metabolite release rates in E. coli, conducted by iteratively leaving out one of the four growth conditions and then predicting those release rates using a linear model trained using the three other ones. The best linear model is a four factor model that includes metabolite value, compound class, intracellular concentration and charge.*

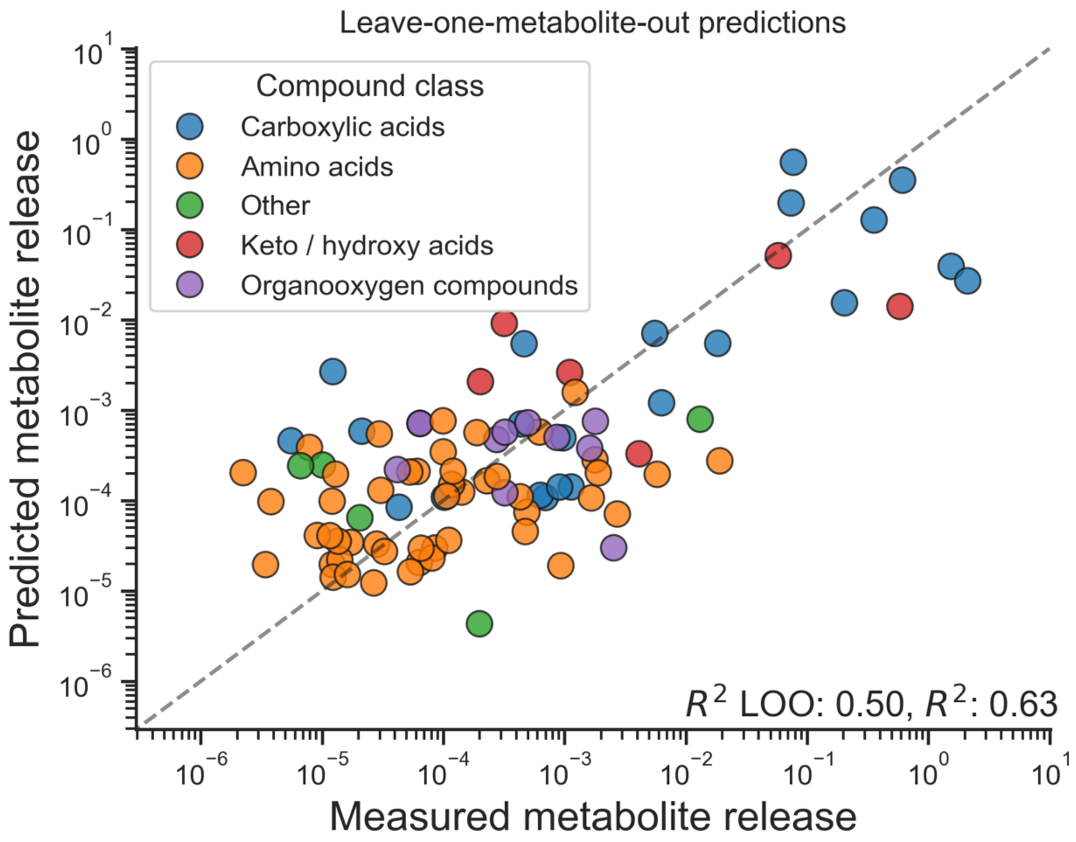


*Figure S13: Predicting release rates for out-of-sample metabolites using the best linear model for predicting release rates in E. coli (same as in Fig. S12).*

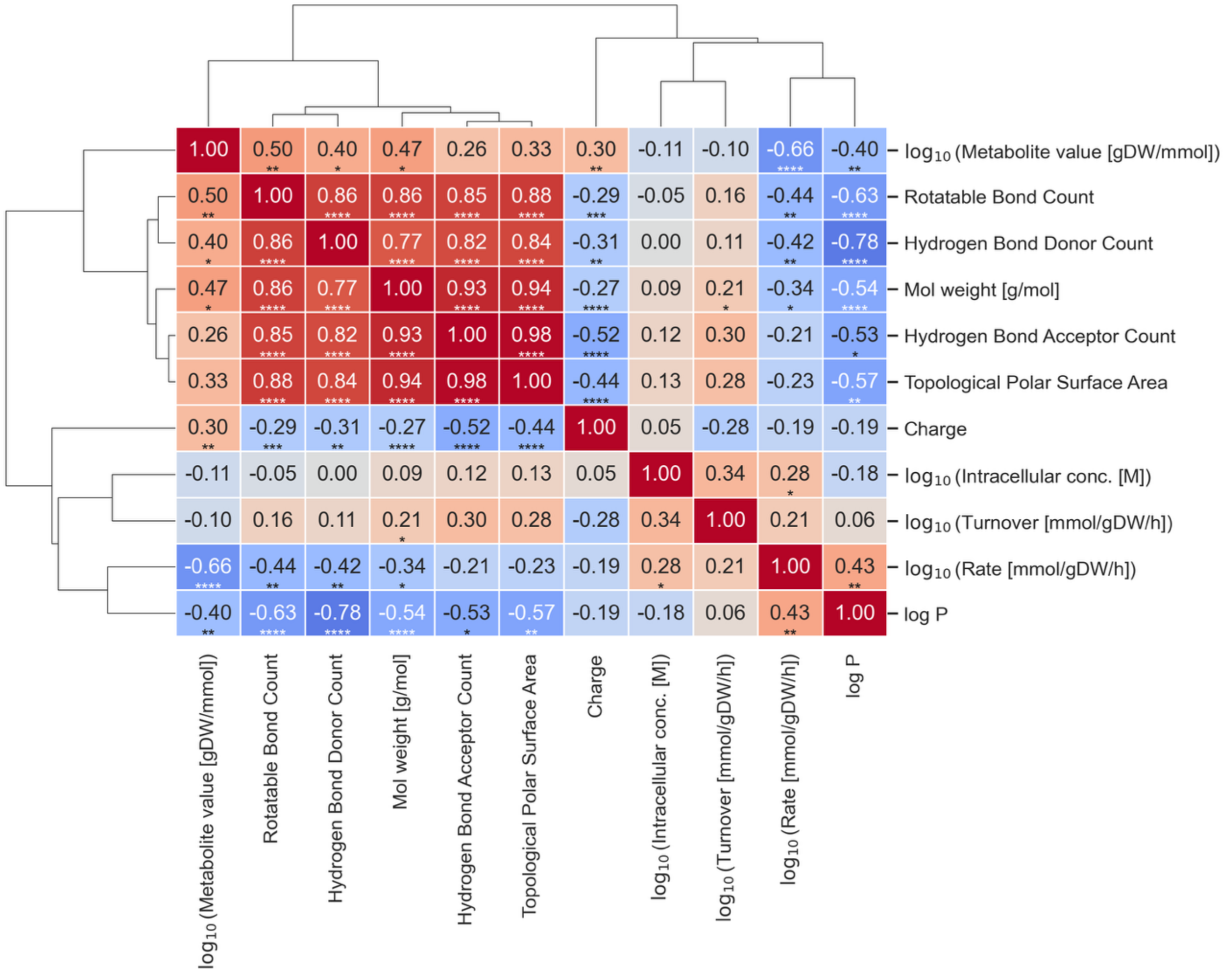


*Figure S14: Correlations between the factors used in Fig. 2A to explain metabolite release rates. The clustermap is based on the same data as Fig. 2A:* E. coli *grown in bioreactors in glucose (Paczia et al. 2012), galactose, L-alanine or L-malate (this study) minimal media (n = 92 observations of 48 metabolites). Values are Pearson correlation coefficients. Because several metabolites were measured in multiple conditions, P values were calculated using standard errors clustered by metabolite (*: P ≤ 0.05; **: P ≤ 0.01; ***: P ≤ 0.001; ****: P ≤ 0.0001). P values are not corrected for multiple testing.*

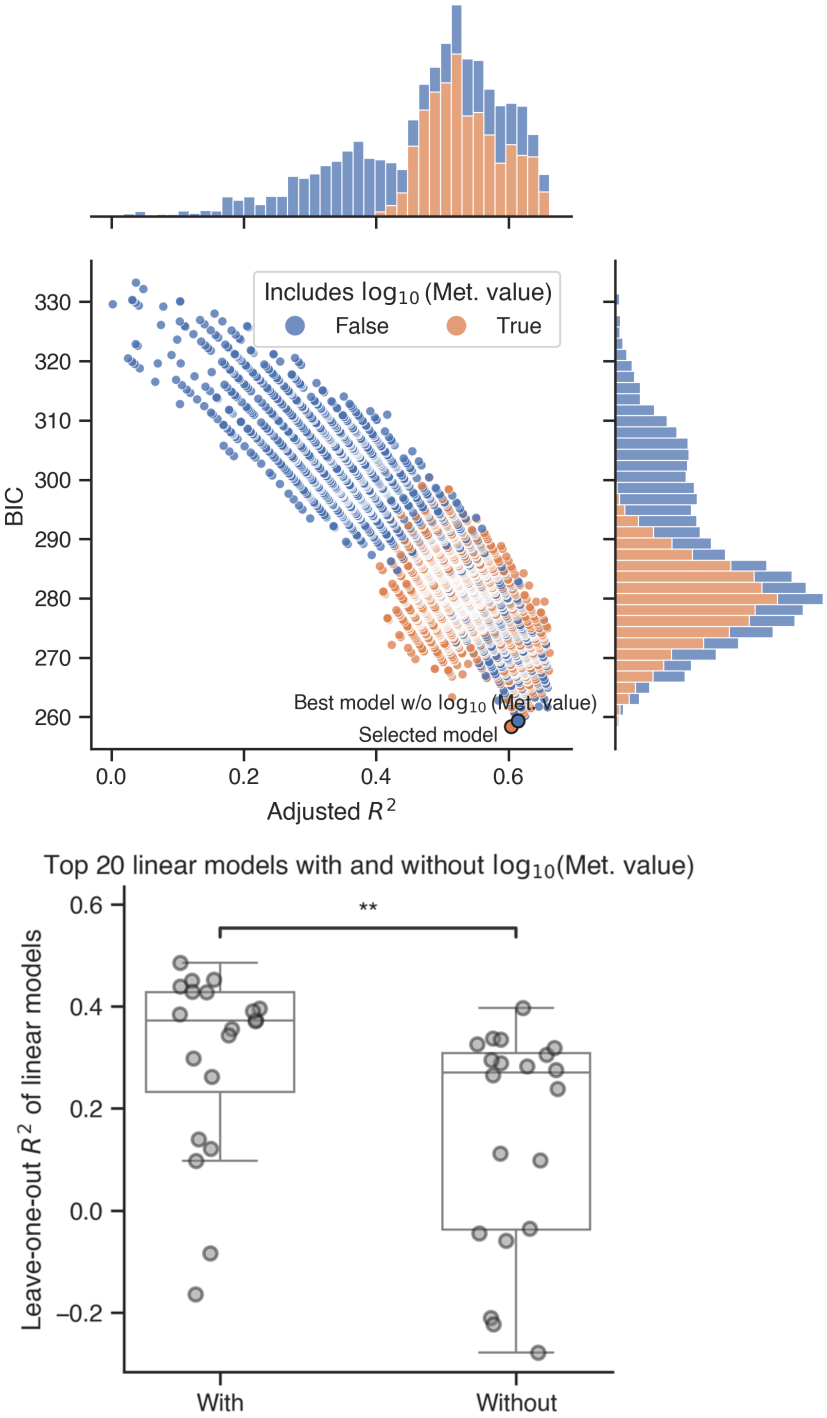


*Figure S15: Top panel shows the Bayesian information criterion (BIC) vs adjusted $R^2$ and their marginal distributions for all possible combinations of linear models. Although most of the best models contain log-transformed metabolite value as a factor, there are other models that obtain the same level of quality as measured by these two criteria. The model used in Figs. S12 and S13 and the best model without metabolite-value as a factor ranked by BIC are marked in orange and blue, respectively. Bottom panel: Out-of-sample predictability when leaving iteratively one of the 4 carbon sources (glucose, galactose, L-alanine, L-malate), as done in Figs. S12 and S13. The top 20 models ranked by BIC with log-transformed metabolite value as a factor are significantly better at predicting out-of-sample conditions ($P = 0.003$, one-sided Mann-Whitney U).*

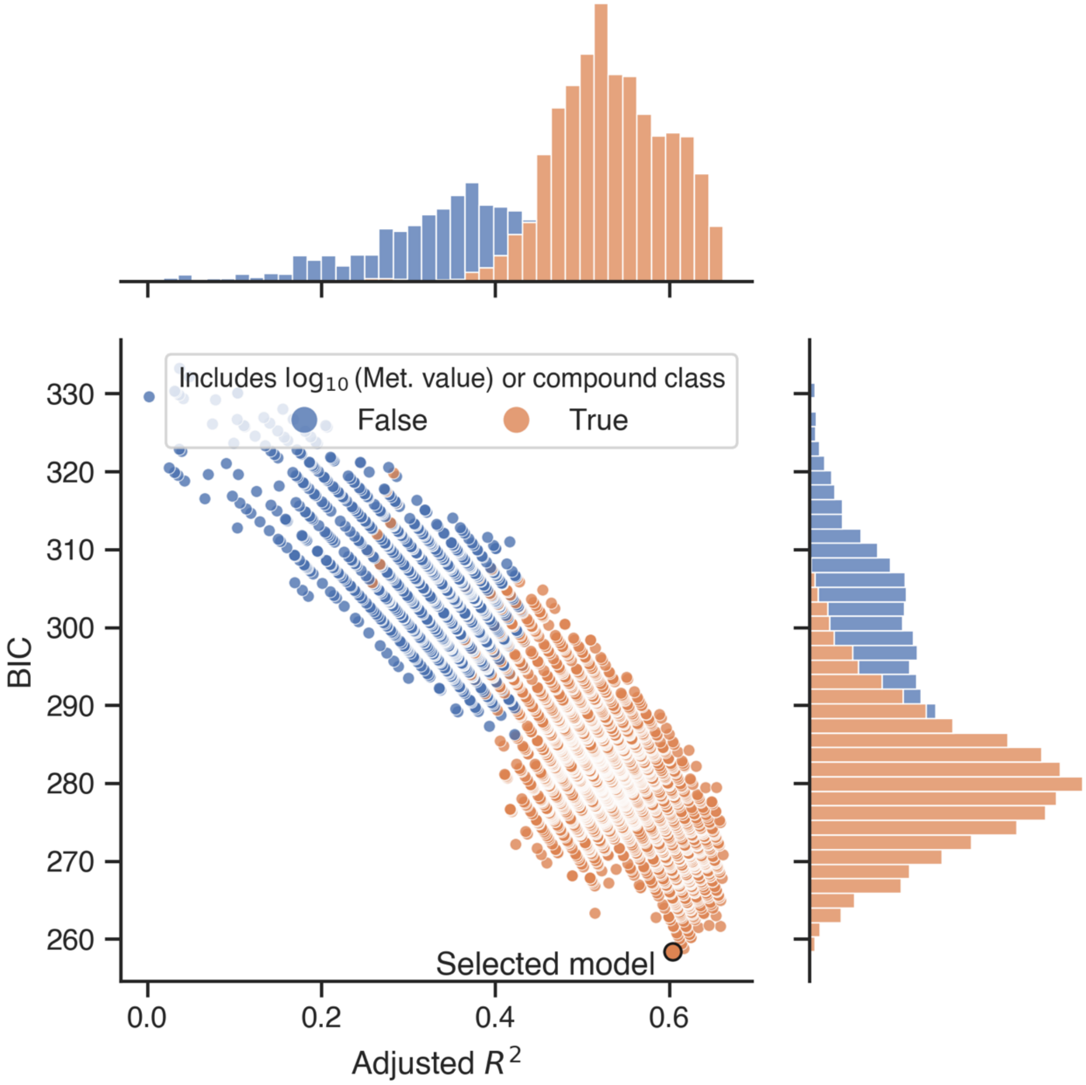


*Figure S16: A scatter plot of Bayesian information criterion (BIC) vs adjusted $R^2$ and their marginal distributions for all possible combinations of linear models. The models are grouped by whether they contain log-transformed metabolite value or compound class as one of the factors. All good models contain either of these factors. The model used in Figs. S12 and S13 is marked in orange.*

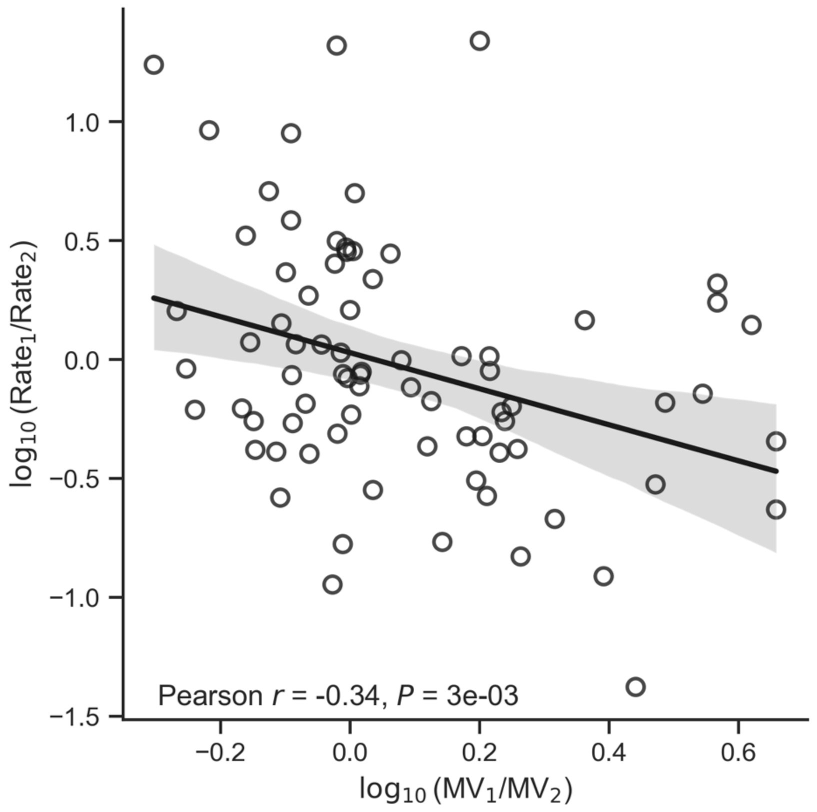


*Figure S17: Negative correlation between log-scaled ratios of metabolite values (x-axis) and release rates (y-axis). Each datapoint represents the comparison for one metabolite between two of the four conditions (glucose, galactose, L-alanine, L-malate) for E. coli, i.e. it is based on the same set of data as in Fig. 2A. However, only pairs of significantly positive release rates are included (Wald test, $P < 0.05$).*

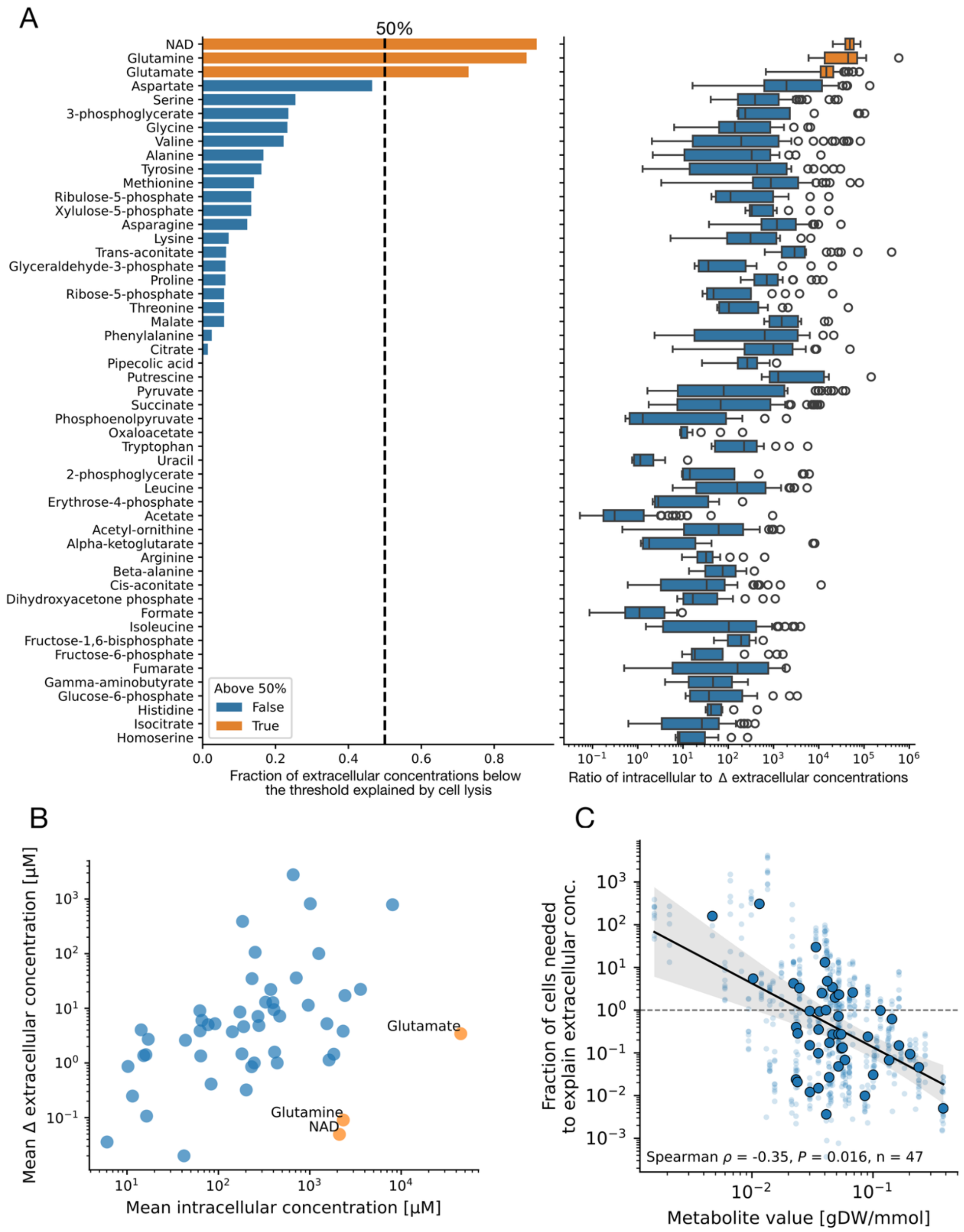


*Figure S18: Fraction of extracellular metabolite measurements in E. coli explained by cell lysis and the associated release of intracellular metabolite pools. A) Only glutamate, glutamine and NAD show the majority of data points in Fig. 3B below the maximum measured cell lysis fraction (2.3%). Note that this is a conservative threshold for the exponential phase, as the mean ± standard deviation of lysed cells is 0.6 ± 0.6% across the three conditions in Fig. 3A. These three metabolites have the highest ratios between intracellular concentrations and the change (Δ) in extracellular concentrations. B) These three metabolites have high ratios because the intracellular concentrations are high or extremely high, while the extracellular concentrations are moderate to low. C) Fraction of the cell population that would need to lyse to explain the measured extracellular concentrations during exponential growth, plotted against metabolite value. Data are from E. coli cultures in this study and from Paczia et al. (2012), as in Fig. 2. Light points show individual samples, and dark points show medians per metabolite. The line and shaded area show a linear regression with 95% confidence interval (slope P = 4e-7, standard errors clustered by metabolite). The dashed line marks the whole cell population. The Spearman correlation was calculated on per-metabolite medians. When also including data from Vila et al. (2023), which were obtained without pH and oxygen control, the relationship had the same direction but was weaker (regression slope P = 0.003; Spearman ρ = −0.12, P = 0.41).*

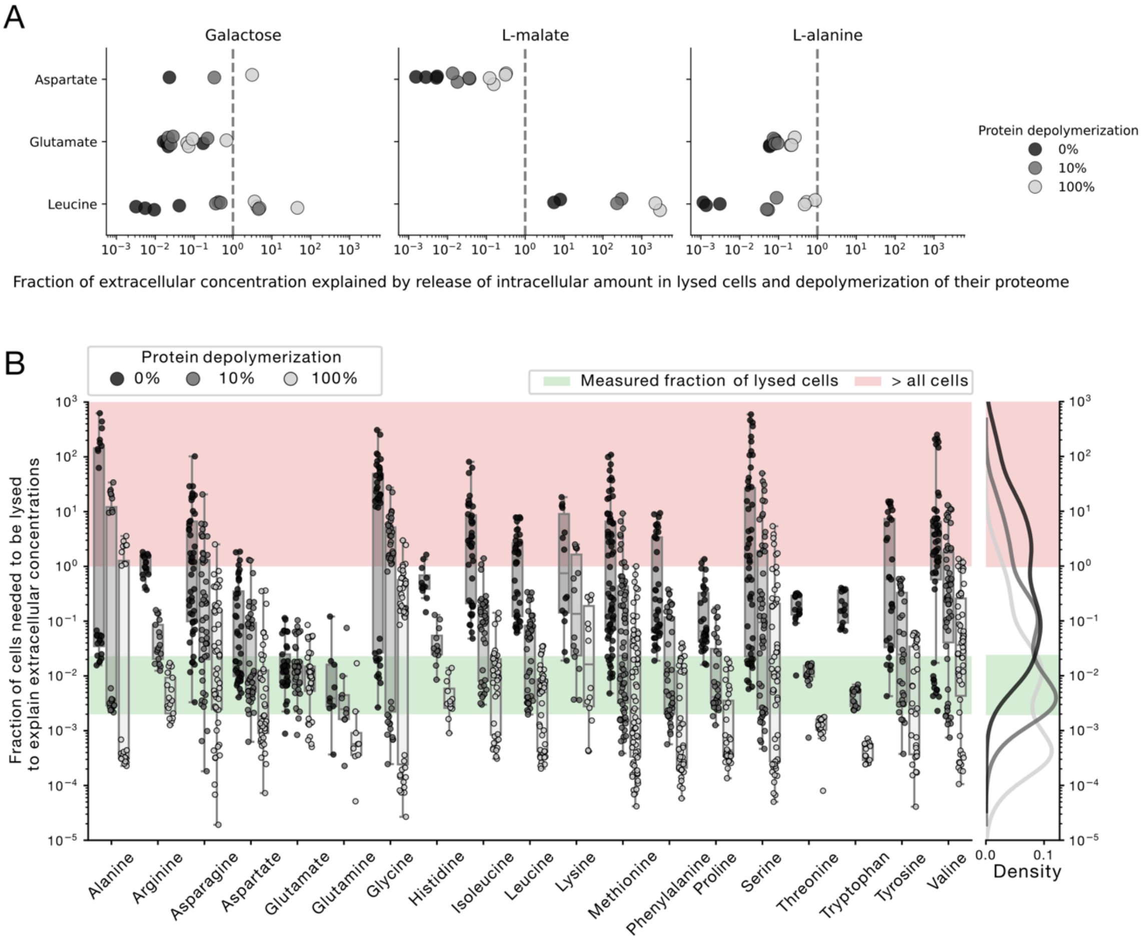


*Figure S19: The potential contribution from depolymerization of the proteome from lysed cells to extracellular amino acid concentrations. A) Estimated fraction of extracellular concentrations explained by intracellular concentrations, estimated fraction of lysed cells and 0%, 10% or 100% proteome depolymerization. B) Using the same levels of proteome degradation, we estimate how large a fraction of the cell population needs to be lysed to explain the extracellular concentrations when both proteome degradation and the release of intracellular concentrations are accounted for.*

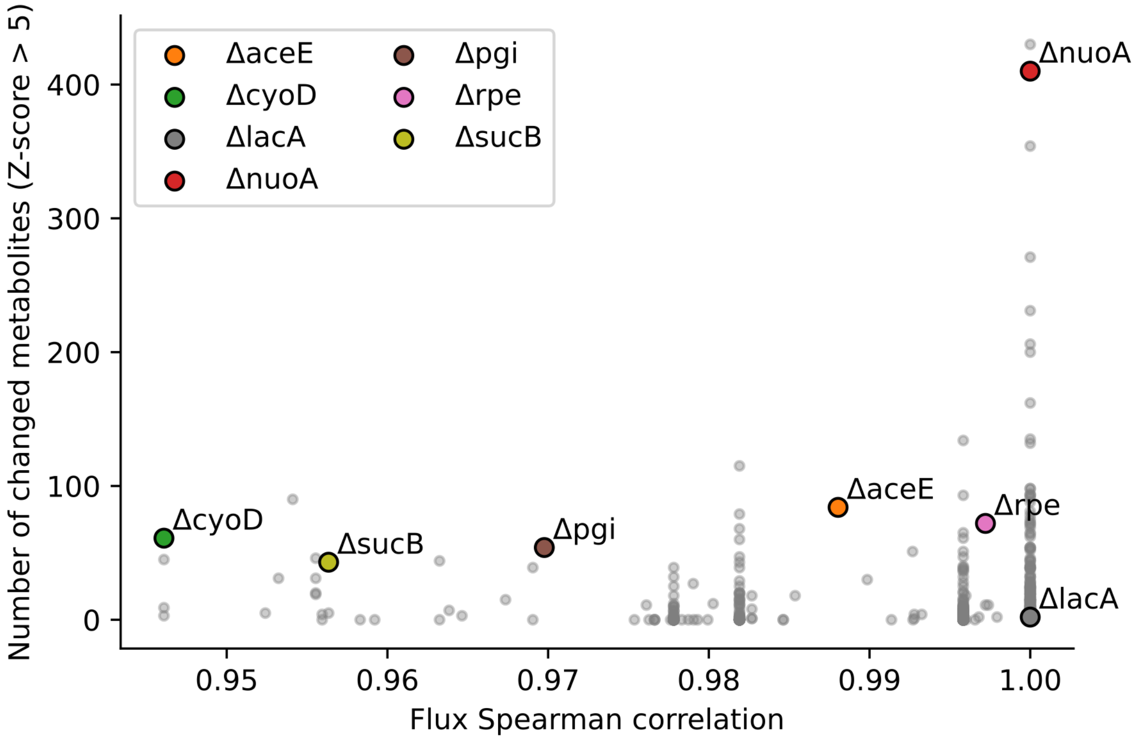


*Figure S20: Selection of knockout mutants that perturb intracellular metabolite fluxes. We used flux balance analysis to identify genes that are non-essential but substantially change intracellular fluxes when knocked out, measured as Spearman correlation against WT fluxes (x-axis). We then also used previous data[3] on intracellular metabolite changes caused by these knockouts to select gene knockout targets known to cause changes in intracellular metabolite levels (y-axis). We ultimately chose these six gene knockout strains affecting different parts of central carbon metabolism (see Fig. 4A), in addition to the negative control ΔlacA.*

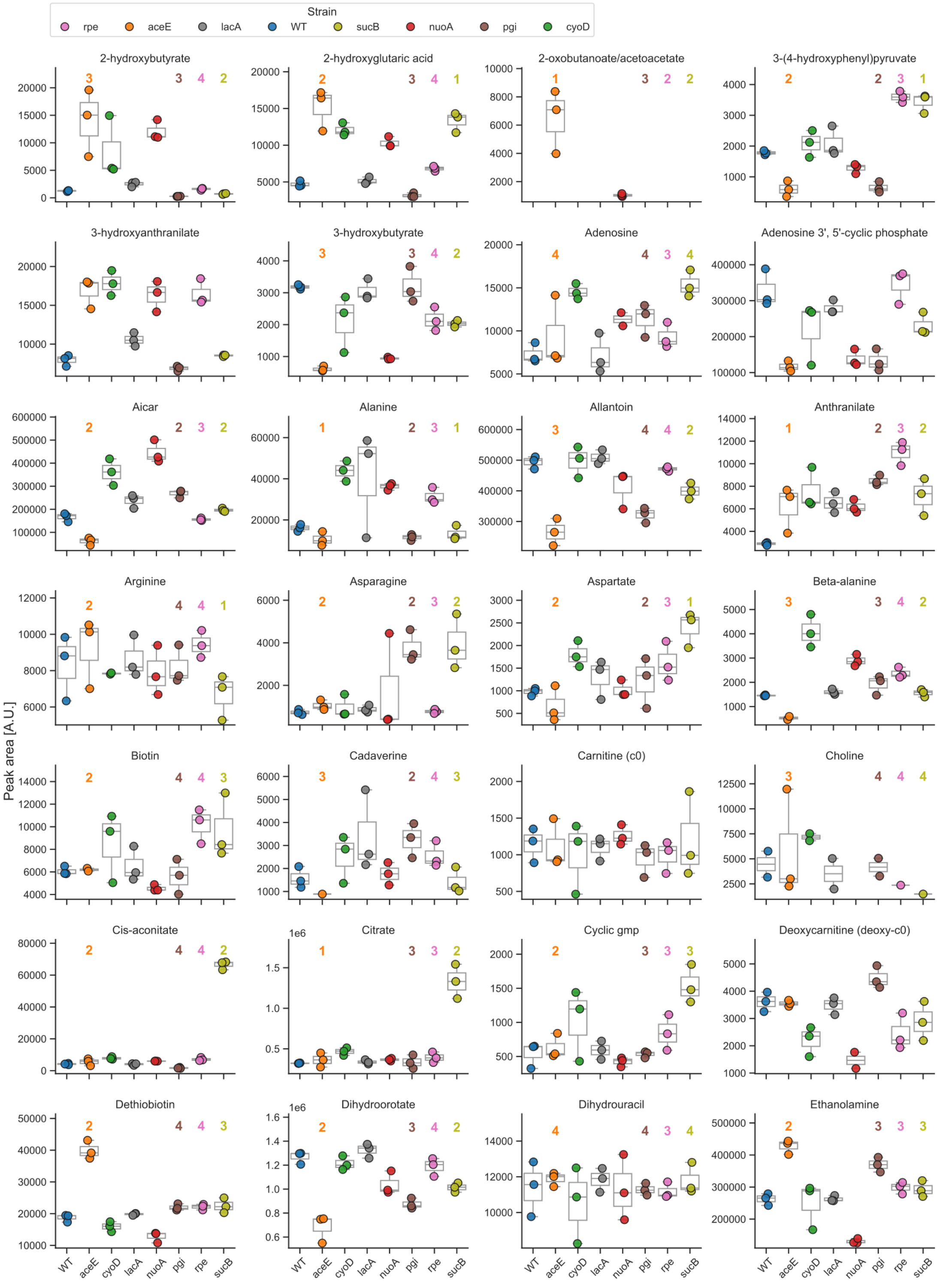


*Figure S21: First subset of the exometabolome data for E. coli WT and the KEIO knockout mutants selected to perturb intracellular metabolite fluxes. The metabolites are sorted alphabetically and the last two subsets of metabolites are presented in Figs. S22 and S23. The numbers above the box plot describe the distance – in terms of the number of reactions – between the metabolite and the reaction(s) affected by the gene knockout. We estimated distances based on the iML1515 GEM, ignoring currency metabolites and therefore also ΔnuoA and ΔcyoD which are related to electron transfer. Distances are not included for metabolites not mapping to iML1515 or for currency metabolites. We observed the most pronounced differences for metabolites in proximity (within 1-2 reactions) of the deleted reaction. For example, the sucB deletion caused the largest changes in the TCA cycle intermediates citrate, isocitrate (Fig. S22), and cis-aconitate, the aceE deletion caused the largest changes in pyruvate (Fig. S23) and lactate (Fig. S22), and the pgi deletion caused the largest changes in glucose 6-phosphate and fructose 6-phosphate (Fig. S22).*

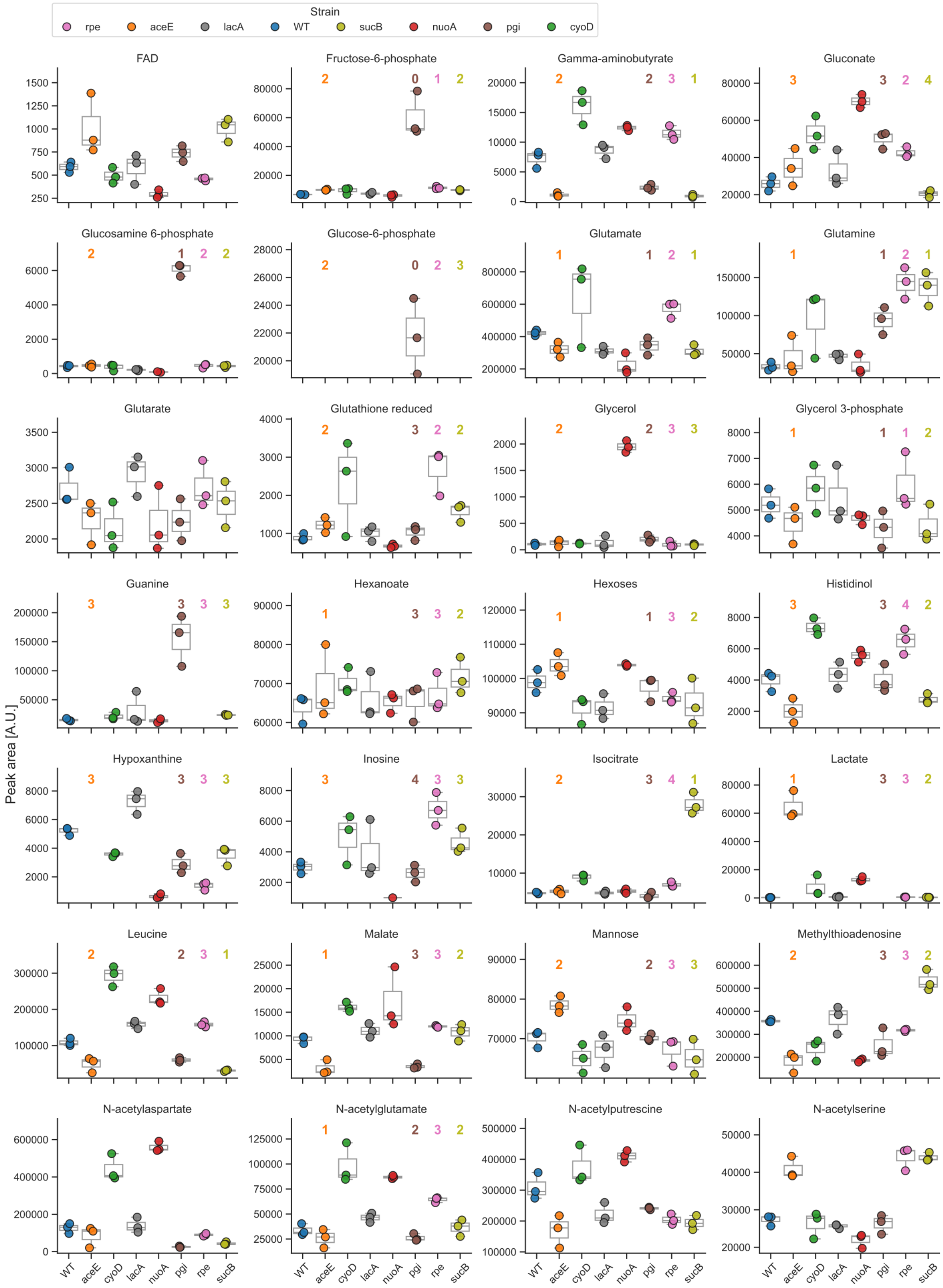


*Figure S22: Second subset of the exometabolome data for E. coli WT and the KEIO knockout mutants selected to perturb intracellular metabolite fluxes. The other two subsets of metabolites are presented in Figs. S21 and S23. The numbers above the box plot describe the distance – in terms of the number of reactions – between the metabolite and the reaction(s) affected by the gene knockout.*

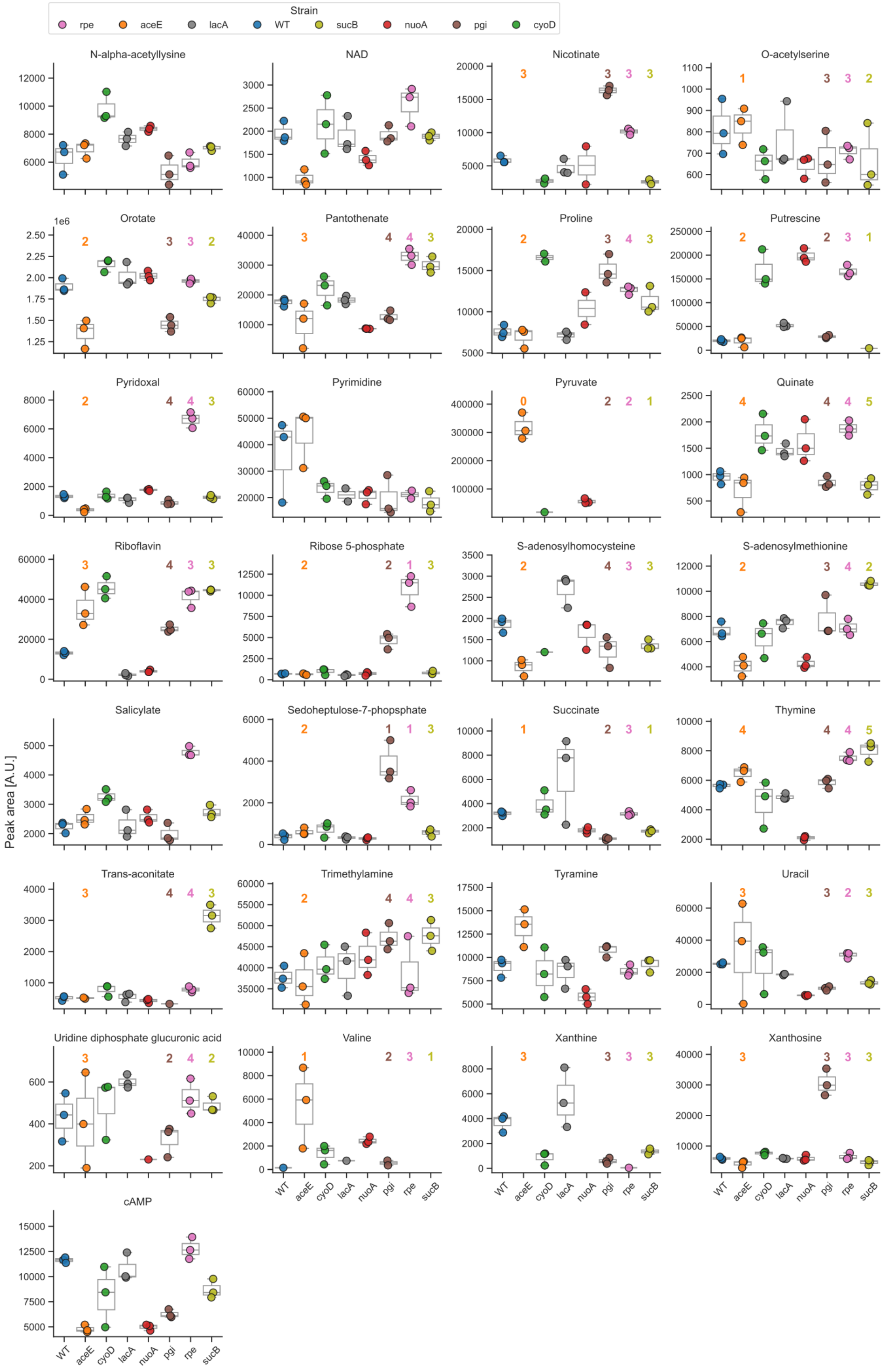


*Figure S23: Last subset of the exometabolome data for E. coli WT and the KEIO knockout mutants selected to perturb intracellular metabolite fluxes. The other two subsets of metabolites are presented in Figs. S21 and S22. The numbers above the box plot describe the distance – in terms of the number of reactions – between the metabolite and the reaction(s) affected by the gene knockout.*

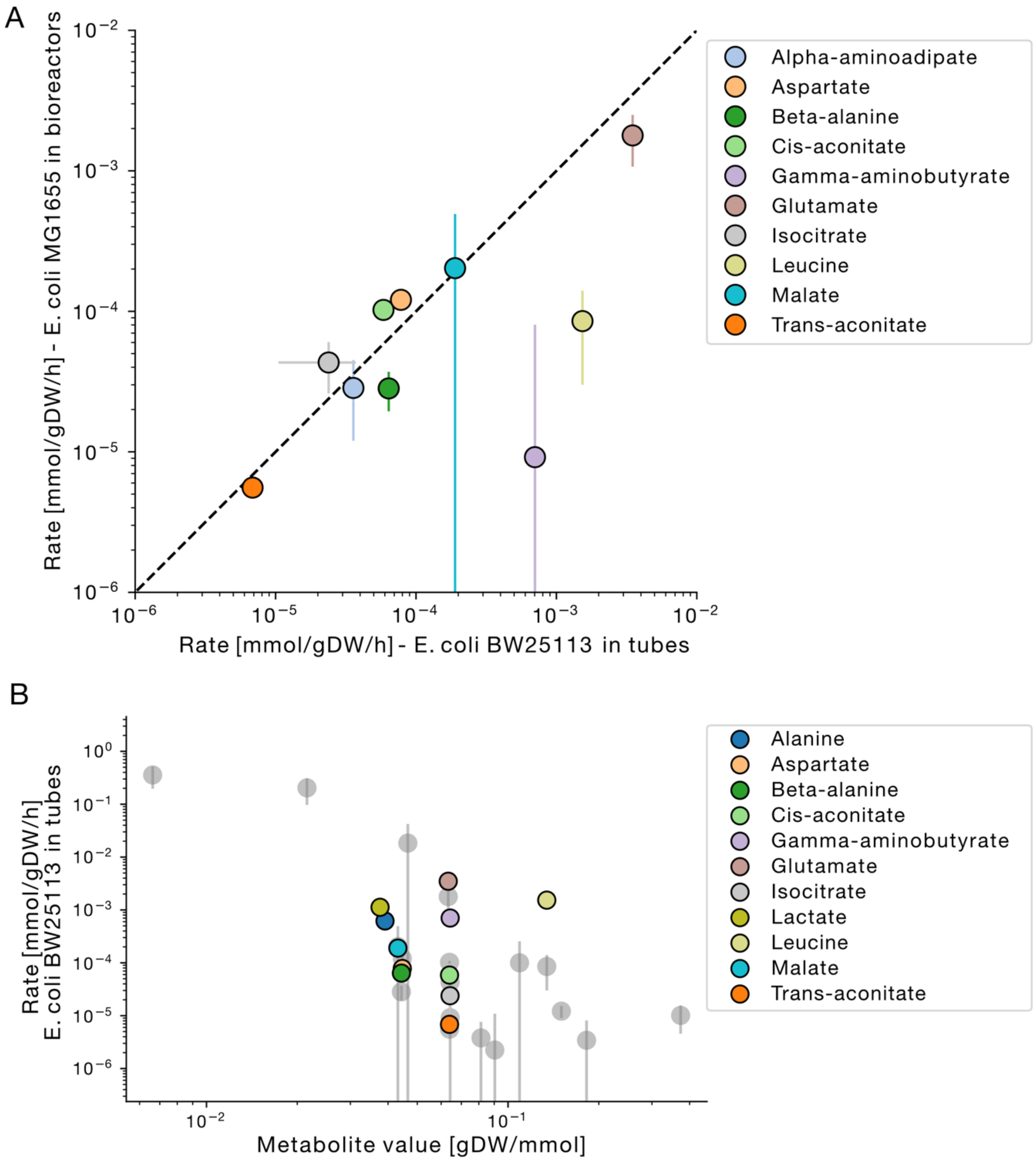


*Figure S24: Comparison of metabolite release rates for E. coli BW25113 with E. coli MG1655 rates presented in Fig. 1, both in M9 galactose medium (40 mM and 20 mM, respectively). A) E. coli BW25113 – cultivated in tubes in shake incubators – has higher release rates of gamma-aminobutyrate and leucine but otherwise similar rates to E. coli MG1655 cultivated in bioreactors with pH and oxygen control. In contrast to the E. coli MG1655 rates, the E. coli BW25113 rates are estimated from a single timepoint in late exponential phase and are thus less reliable than the MG1655 rates. The error bars for E. coli MG1655 represent standard errors of the slope estimate, while the error bars for BW25113 are standard errors of measured concentrations scaled by the AUC of biomass to correspond to release rates. The variation is larger in the slope estimates than in the single timepoint concentrations, explaining the apparently higher uncertainty associated with the MG1655 rates. B) The E. coli BW25113 rates plotted on top of the E. coli MG1655 rates (in gray, panel 1 of Fig. 1E). The metabolite values here are the same for E. coli MG1655 and E. coli BW25113.*

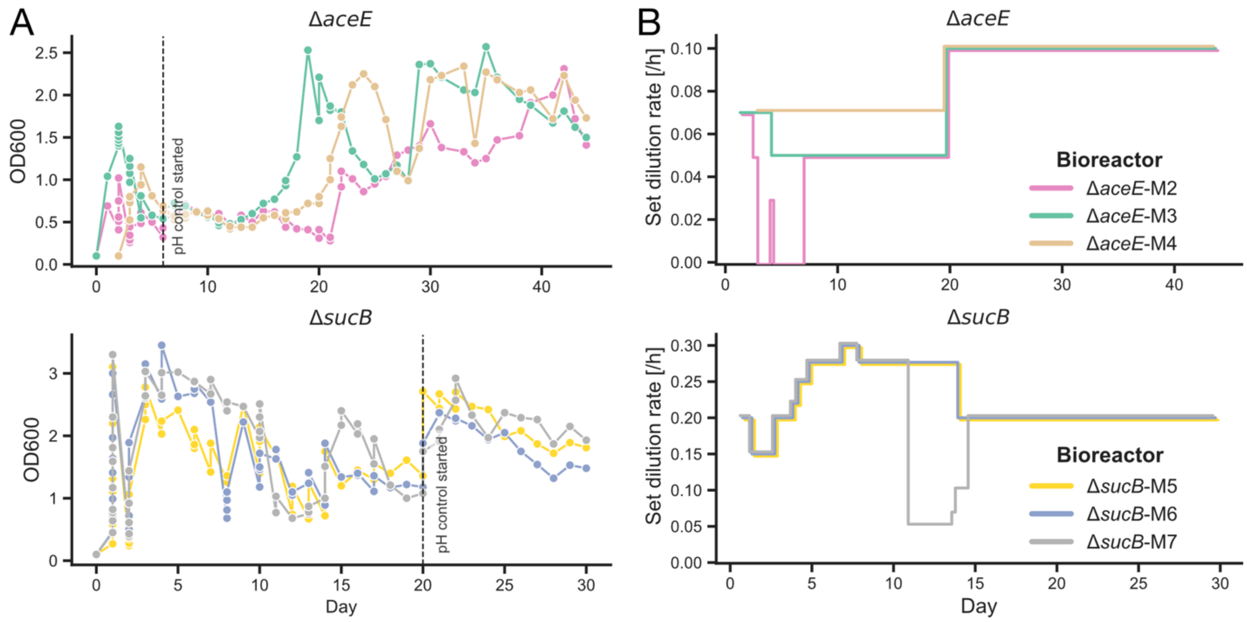


*Figure S25: Chemostat experiments of KEIO mutants ΔaceE and ΔsucB. A) Optical density ($OD_{600}$) in each bioreactor throughout the experiment. Changes in culture dynamics required adjustments of the dilution rate and the introduction of pH control to maintain the cultures. B) Set dilution rate of the six bioreactors throughout the experiment. The dilution rate was adjusted several times during the first half of the experiment and kept constant during the second half.*

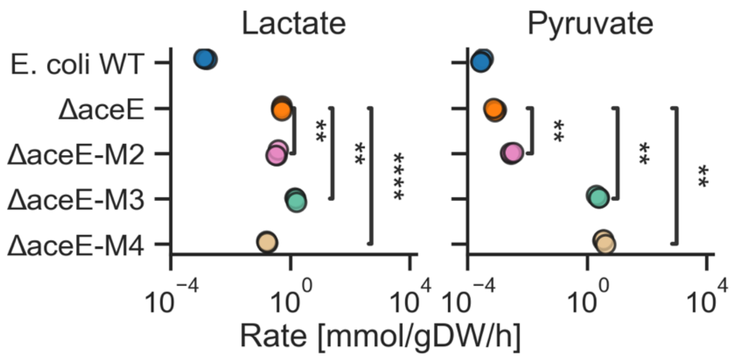


*Figure S26: Release rates of lactate and pyruvate in WT, ΔaceE ancestor, and evolved ΔaceE strains estimated from measured extracellular concentrations at $OD_{600}$~1 to account for differences in growth rate between strains. Statistical significance was assessed using Welch's t-test (ns: P > 0.05; *: P ≤ 0.05; **: P ≤ 0.01; ***: P ≤ 0.001; ****: P ≤ 0.0001).*

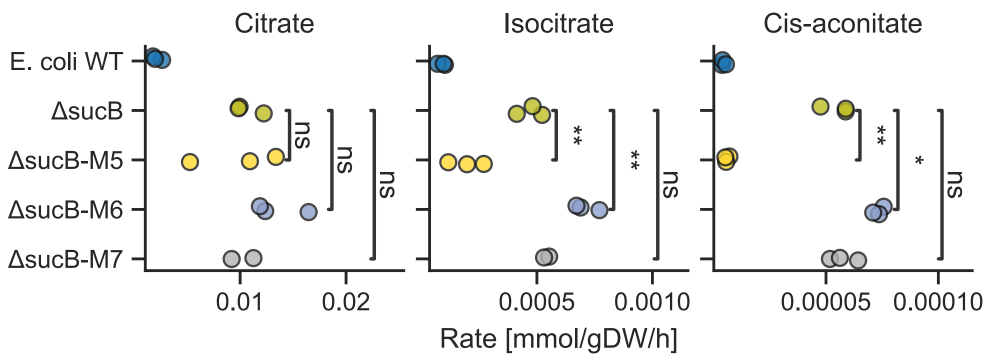


*Figure S27: Release rates of citrate, isocitrate and cis-aconitate in WT, ΔsucB ancestor, and evolved ΔsucB strains estimated from measured extracellular concentrations at $OD_{600}$~1 to account for differences in growth rate between strains. Statistical significance was assessed using Welch's t-test (ns: P > 0.05; *: P ≤ 0.05; **: P ≤ 0.01; ***: P ≤ 0.001; ****: P ≤ 0.0001).*

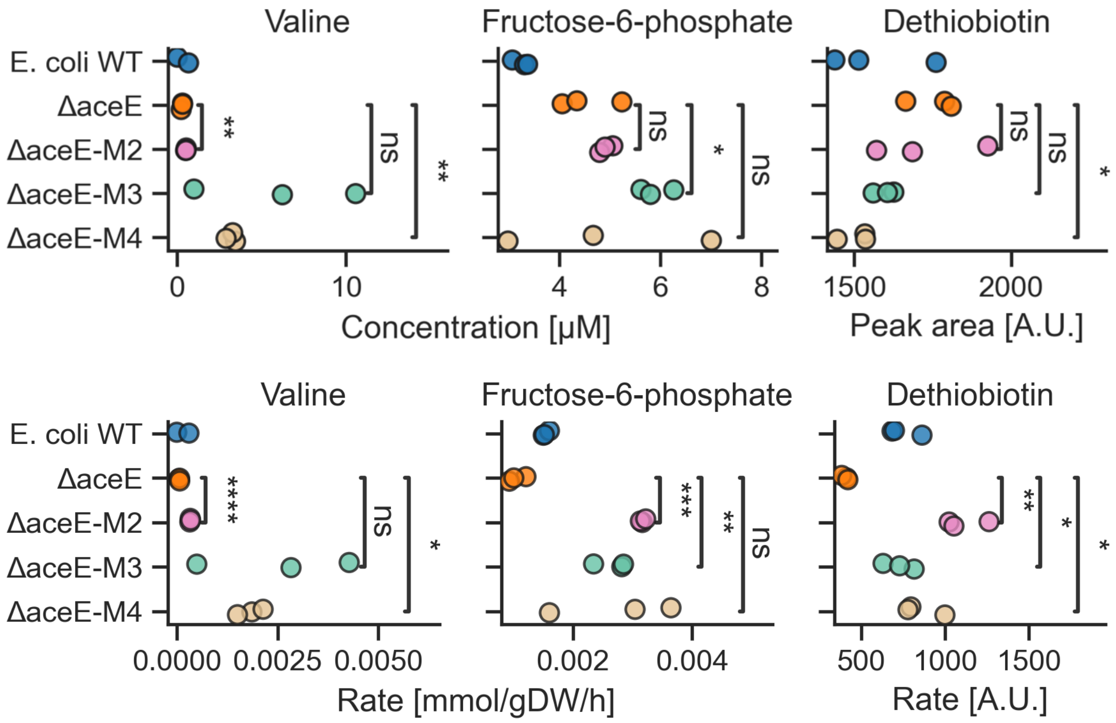


*Figure S28: The top row shows the extracellular concentrations of valine, fructose 6-phosphate and dethiobiotin in WT, ΔaceE ancestor, and evolved ΔaceE strains. The bottom row shows the same data but adjusted according to differences in growth by calculating mean release rates. Dethiobiotin could not be quantified to absolute levels and is therefore reported in arbitrary units [A.U.]. Statistical significance was assessed using Welch's t-test (ns: P > 0.05; *: P ≤ 0.05; **: P ≤ 0.01; ***: P ≤ 0.001; ****: P ≤ 0.0001).*

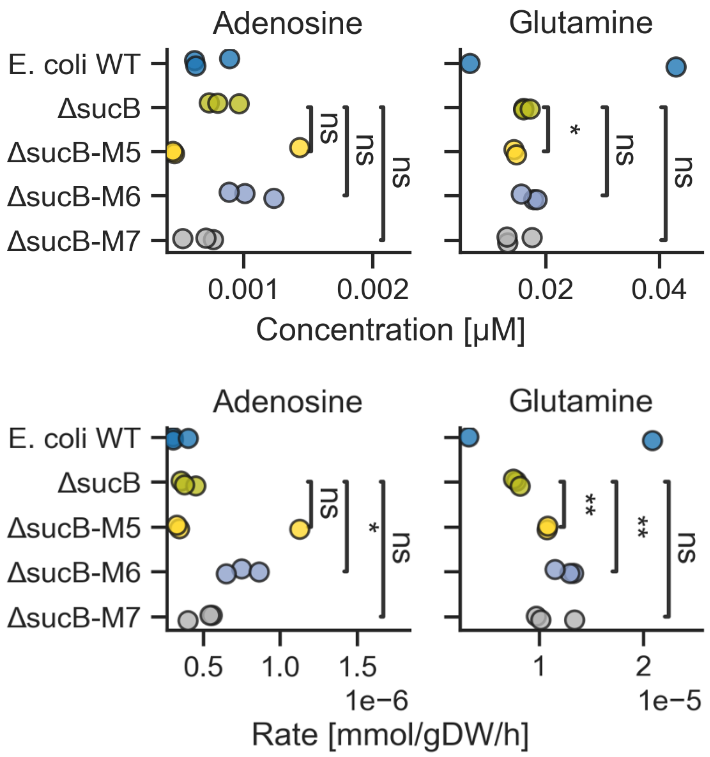


*Figure S29: The top row shows the extracellular concentrations of adenosine and glutamine in WT, ΔsucB ancestor, and evolved ΔsucB strains. The bottom row shows the same data but adjusted according to differences in growth by calculating mean release rates. Statistical significance was assessed using Welch's t-test (ns: P > 0.05; *: P ≤ 0.05; **: P ≤ 0.01; ***: P ≤ 0.001; ****: P ≤ 0.0001).*

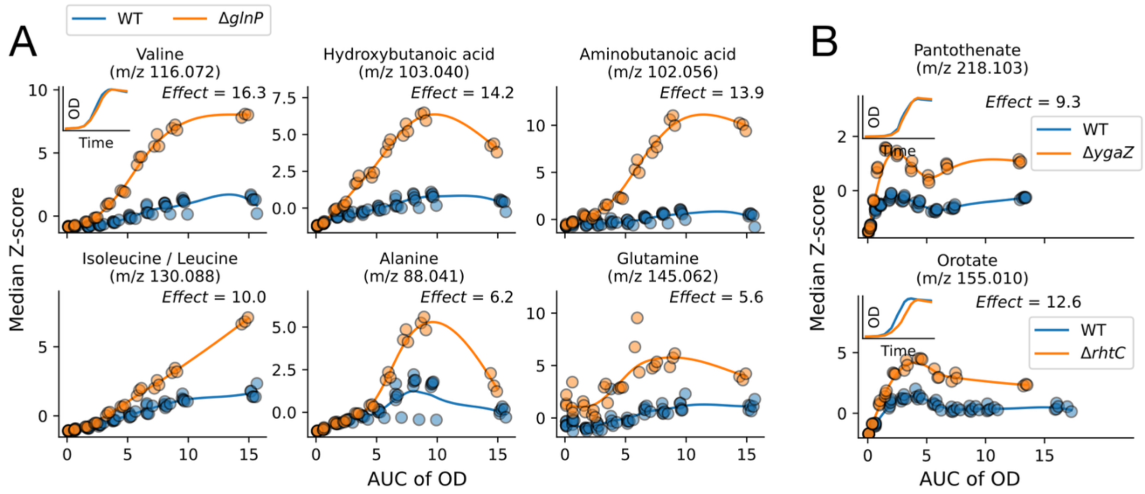


*Figure S30: A) Many positive and negative effects extended beyond known transporter annotations. For example, the knockout strain ΔglnP (lacking a subunit of a glutamine ABC transporter) increased, in addition to glutamine, also the extracellular concentrations of valine, hydroxybutanoic acid, aminobutanoic acid, isoleucine/leucine, and alanine (Effect > 3, P < 0.005). B) Positive effects were also observed for several transporters primarily associated with metabolite efflux, including YgaZ and RhtC.*

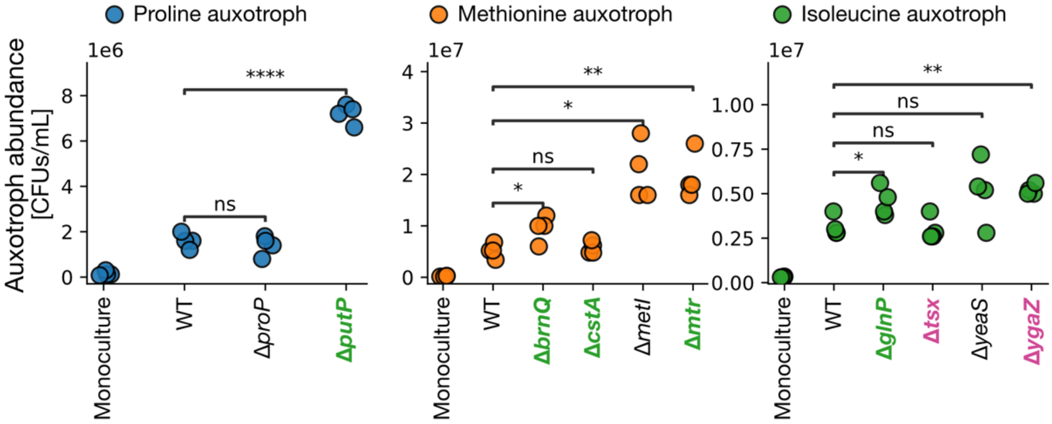


*Figure S31***:** *Growth of* E. coli *auxotrophs in coculture with transporter knockout strains in glucose minimal medium. Auxotroph abundance (CFU/mL) in monoculture and in coculture with the wild-type reference E. coli BW25113 (WT) or transporter knockout strains. In 4 of 5 cases, knockout strains with a strong positive effect on the amino acid required by the auxotroph (Effect > 3, P < 0.005, green labels) significantly improved auxotroph growth relative to coculture with WT. Contrary to expectations, ΔygaZ, one of the two strains with a strong negative effect (Effect < −3, P < 0.005, pink labels), supported growth of the isoleucine auxotroph better than WT. The remaining knockout strains had significant but weaker effects (ΔproP: 1.1, ΔmetI: 2.6, ΔyeaS: 2.5; P < 0.005). Statistical significance relative to WT was assessed using Welch's t-test (ns: P > 0.05; *: P ≤ 0.05; **: P ≤ 0.01; ****: P ≤ 0.0001).*

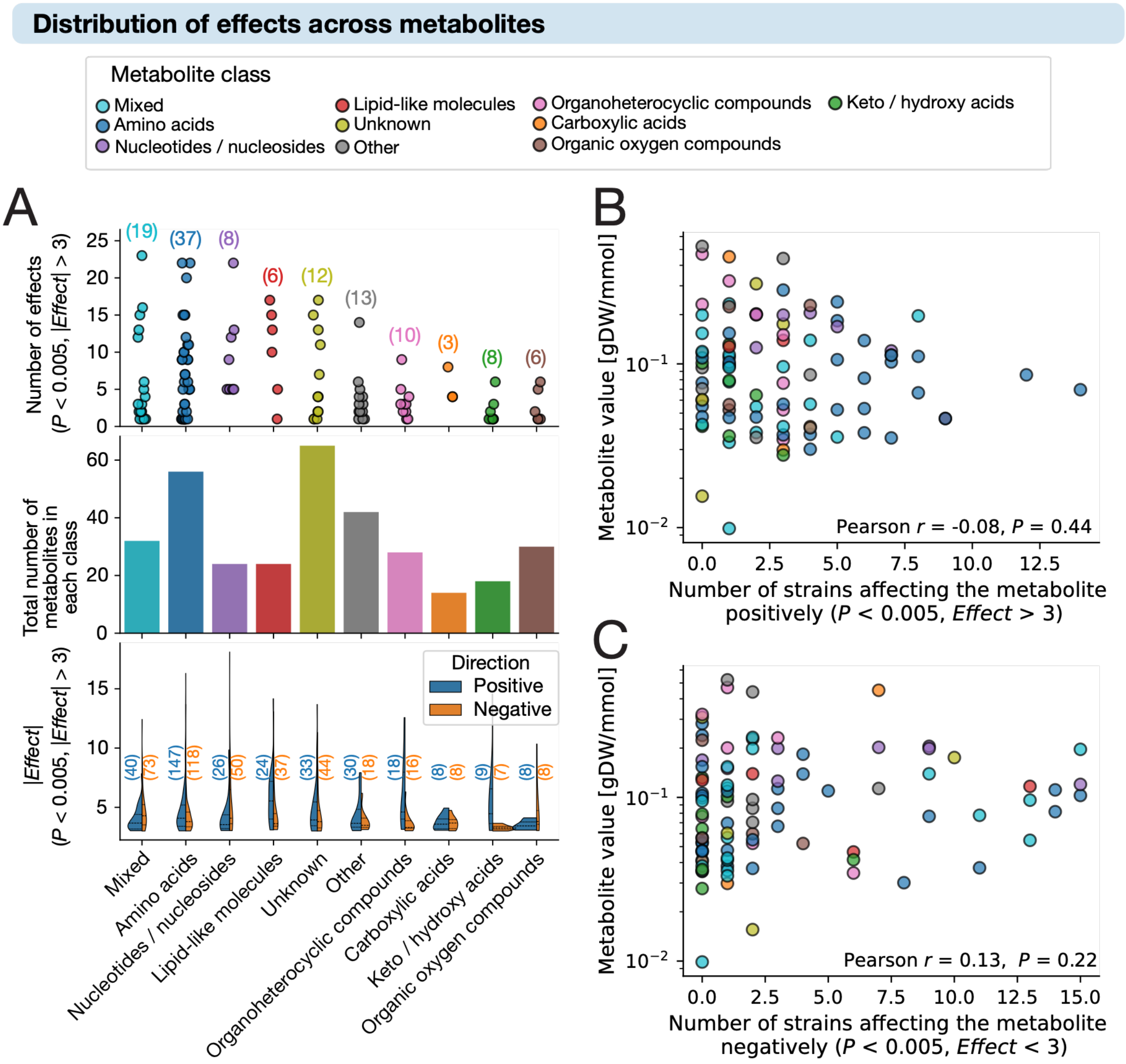


*Figure S32: Distribution of effects across metabolites and metabolite classes. A) Number of effects per metabolite (top) and distribution of effect sizes (bottom), grouped by metabolite class (P < 0.005, |Effect| > 3). Metabolites not affected by any transporter knockout are not included. No significant differences were observed across classes (ANOVA). B-C) No correlation was detected between metabolite value and how frequently it was affected by transporter knockout, either positively (B) or negatively (C).*

### Distribution of effects across knockout strains

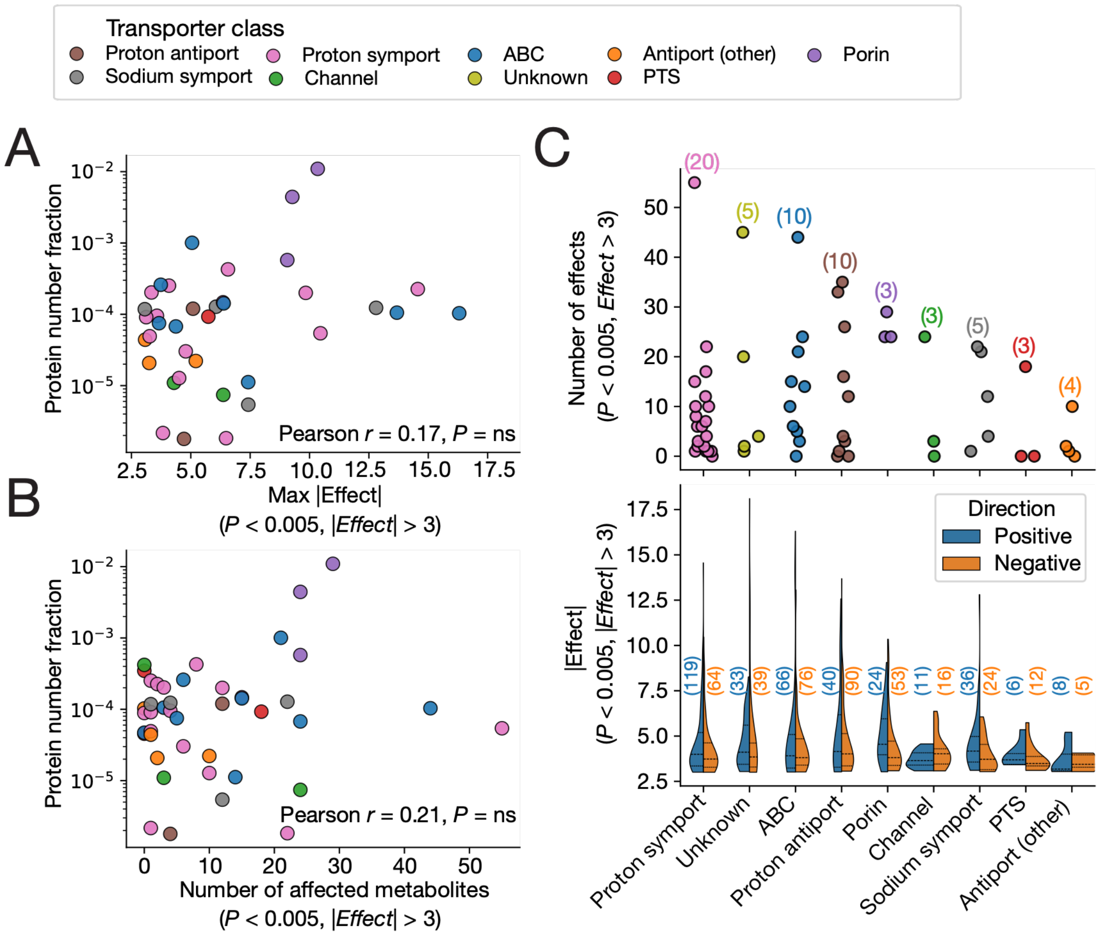


*Figure S33: Distribution of effects across knockout strains and transporter classes. A-B) No significant correlation was found between average protein expression level (mean protein number fraction across carbon-limited conditions)[4] and either the maximum effect size (A) or the number of affected metabolites (B). C) The number of effects per knockout strain (top) and distribution of effect sizes grouped by transporter class (bottom). No significant differences were observed across classes (ANOVA).*

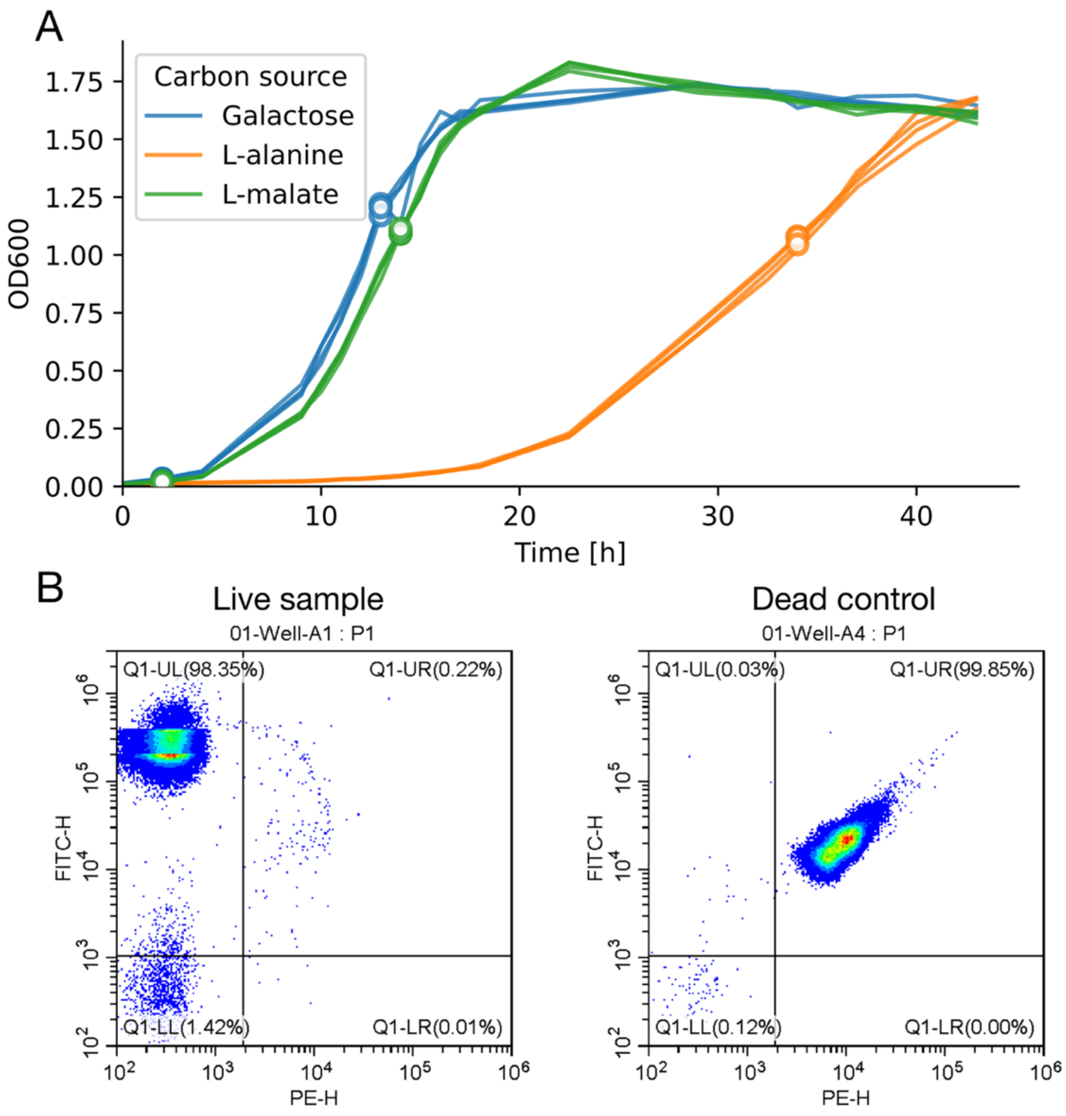


*Figure S34: A) Growth curves for E. coli K-12 MG1655 and sampling timepoints (marked by open circles) for paired intra- and extracellular metabolomics sampling and live/dead staining assays to quantify fraction of lysed cells. All replicates (four per carbon source) were sampled after two hours and at $OD_{600} \approx 1.1$. B) Example of gating strategy to quantify the fraction of live and lysed (dead) cells in samples. The left panel shows the live sample, and the right panel shows the lysed (dead) control used to define gating thresholds. The live population was quantified from the upper left quadrant, and the lysed population was quantified from the upper right quadrant.*
